# High-density linkage maps and chromosome level genome assemblies unveil direction and frequency of extensive structural rearrangements in wood white butterflies (*Leptidea* spp.)

**DOI:** 10.1101/2022.10.10.510802

**Authors:** L. Höök, K. Näsvall, R. Vila, C. Wiklund, N. Backström

**Author notes:** Correspondence: Lars Höök, Karin Näsvall, Evolutionary Biology Program, Department of Ecology and Genetics, Uppsala University. Norbyvägen 18D, 752 36 Uppsala, Sweden. Equal author contribution (L. Höök, K. Näsvall).

## Abstract

Karyotypes are generally conserved between closely related species and large chromosome rearrangements typically have negative fitness consequences in heterozygotes, potentially driving speciation. In the order Lepidoptera, most investigated species have the ancestral karyotype and gene synteny is often conserved across deep divergence, although examples of extensive genome reshuffling have recently been demonstrated. The genus *Leptidea* has an unusual level of chromosome variation and rearranged sex chromosomes, but the extent of restructuring across the rest of the genome is so far unknown. To explore the genomes of the wood white (*Leptidea*) species complex, we generated eight genome assemblies using a combination of 10X linked reads and HiC data, and improved them using linkage maps for two populations of the common wood white (*L. sinapis*) with distinct karyotypes. Synteny analysis revealed an extensive amount of rearrangements, both compared to the ancestral karyotype and between the *Leptidea* species, where only one of the three Z chromosomes was conserved across all comparisons. Most restructuring was explained by fissions and fusions, while translocations appear relatively rare. We further detected several examples of segregating rearrangement polymorphisms supporting a highly dynamic genome evolution in this clade. Fusion breakpoints were enriched for LINEs and LTR elements, which suggests that ectopic recombination might be an important driver in the formation of new chromosomes. Our results show that chromosome count alone may conceal the extent of genome restructuring and we propose that the amount of genome evolution in Lepidoptera might still be underestimated due to lack of taxonomic sampling.

## Introduction

The karyomorph is the highest order of organization of the genetic material and understanding the mechanistic underpinnings and micro- and macro-evolutionary effects of changes in chromosome numbers are long-standing goals in evolutionary biology (Mayrose & Lysak, 2021). Chromosome rearrangements have for example been suggested to be important drivers of speciation as a consequence of meiotic segregation problems and suppressed recombination in chromosomal heterozygotes (Faria & Navarro, 2010; Rieseberg, 2001). The number and structure of chromosomes are generally conserved within species and between closely related taxa, while substantial karyotypic changes can occur over longer evolutionary distances (Román-Palacios et al., 2021; Ruckman et al., 2020). In contrast, there are some examples of considerable chromosome rearrangement rate differences between closely related species, but the underlying reasons for why karyotypic change have occurred comparatively rapidly in some lineages are largely unexplored (de Vos et al., 2020; Ruckman et al., 2020; Sylvester et al., 2020). Chromosomal rearrangements, in particular fission and fusion events, are likely often underdominant and/or deleterious and the probability of their fixation in a population should therefore be higher in organisms with lower effective population size (*N_e_*) (Pennell et al., 2015), or with strong meiotic drive (Blackmon et al., 2019). Additionally, the consequences of fissions and fusions seem to depend on the types of chromosomes a species harbor. In species with holokinetic chromosomes (holocentric), the spindle apparatus can bind to multiple positions along the chromosomes, while the attachment is restricted to the specific centromere region in monocentric species (Melters et al., 2012). As a consequence, chromosome fissions and fusions do not necessarily lead to meiotic segregation problems in holocentric species (Faulkner, 1972; Lukhtanov et al., 2018). This has been hypothesized to lead to a higher rate of chromosome number evolution in holocentric compared to monocentric species. However, the evidence for such a rate difference have been mixed and a recent, large-scale meta-analyses across insects suggests that chromosome evolution is not significantly faster in holocentric species (Ruckman et al., 2020), and the explanation for rate differences between species might rather be lineage specific life-history traits or demographics (Kawakami et al., 2009; Larson et al., 1984; Petitpierre, 1987).

The order Lepidoptera, moths and butterflies, constitutes one of the most species rich and widespread animal groups and has for long been a popular study system in ecology and evolution, in large part due to the high diversity and the eye-catching colour pattern variations that have attracted both amateur naturalists and academic scholars for centuries (Boggs et al., 2003). Lepidoptera also share several key genetic features, like holocentric chromosomes (Suomalainen, 1953), and female heterogamety and achiasmy (Traut et al., 2007; Turner & Sheppard, 1975), which present interesting aspects regarding chromosomal rearrangements and their evolutionary consequences. The karyotype structure has generally been found to be conserved across lepidopteran genera, with most investigated taxa having the inferred ancestral haploid chromosome number of n = 31 (de Vos et al., 2020; Robinson, 1971). However, the genera *Agrodiaetus* (n = 10 - 134, Kandul et al., 2007) and *Godyris* (n = 13 - 120, Brown et al., 2004), for example, have been shown to have an elevated rate of karyotype evolution compared to other insects (Ruckman et al., 2020). On a more detailed level, genomic analyses have also unveiled a high level of conserved gene synteny between divergent lepidopteran lineages (Ahola et al., 2014; Davey et al., 2016; Pringle et al., 2007). Extensive, genome-wide chromosomal restructuring has so far only been detected in *Pieris napi / P. rapae* (Hill et al., 2019). However, since the analyses have been limited to a comparatively small set of taxonomic lineages where highly contiguous genome assemblies and / or linkage maps are available, the levels of both karyotype variation and intra-chromosomal rearrangements in Lepidoptera are likely underestimated (de Vos et al., 2020).

An attractive model system for studying karyotype evolution is the Eurasian wood white butterflies in the genus *Leptidea* (family Pieridae). The three species, common wood white (*Leptidea sinapis*), Réal’s wood white (*Leptidea reali*) and the cryptic wood white (*Leptidea juvernica*), form a species complex of morphologically nearly indistinguishable species that show remarkable inter- and intraspecific variation in chromosome numbers (Dincă et al., 2011; Lukhtanov et al., 2011). Cytogenetic analyses have shown that *L. reali* has rather few chromosomes and less intraspecific variation (n = 25 – 28), while *L. juvernica* has a considerably more fragmented and variable karyotype (n = 38 – 46) (Dincā et al., 2011; Šíchová et al., 2015). The most striking chromosome number variation has been found in *L. sinapis*, which has one of the most extreme non-polyploid, intraspecific karyotype clines of all eukaryotes, ranging from n ≈ 53 – 55 in southern Europe and gradually decreasing to n ≈ 28 – 29 in northern Europe and n ≈ 28 – 31 in central Asia (Lukhtanov et al., 2011, 2018). In addition, the common ancestor of the species complex has undergone translocations of autosomal genes to the sex chromosome(s) and extension of the ancestral Z chromosome that has resulted in a set of neo sex chromosomes (Šíchová et al., 2015; Yoshido et al., 2020). *Leptidea* butterflies have also experienced a relatively recent burst of transposable element activity (Talla et al., 2017), elements that could act as important drivers of chromosome rearrangements (Belyayev, 2014), but whether this has been important for karyotype evolution specifically in this genus remains to be explored. Altogether, these findings motivate detailed characterization of the rate and direction of the chromosome rearrangements and assessment of potential drivers of the rapid karyotype changes in this lineage.

In order to dissect the chromosome structure in detail and quantify rates and directions of chromosome rearrangements, we sequenced and assembled the genome of a male and female individual of *L. juvernica, L. reali* and the Swedish and Catalan populations of *L. sinapis* using a combination of 10X linked reads and HiC-scaffolding. In addition, we generated genetic maps for the two *L. sinapis* populations and used the linkage information to super-scaffold and correct the physical genome assemblies. The assembled chromosomes were compared between wood white species and with the inferred ancestral lepidopteran karyotype to characterize rates and patterns of fissions, fusions and intra-chromosomal rearrangements across the species complex. Furthermore, we quantified enrichment of different types of genetic elements in fission and fusion breakpoints to assess if specific genomic features have been associated with chromosome rearrangements.

## Results

The genome assemblies had extensive contiguity, high BUSCO scores (Supplementary table 1), few gaps and the majority of sequence contained in chromosome-sized scaffolds (Table 1, Supplementary figures 1 and 2). The implementation of linkage map information for the Swedish and Catalan *L. sinapis* (Figure 1) allowed for correction of these particular genome assemblies. Overall, we found a high level of collinearity between physical maps and linkage maps, but several large inversions were corrected (Supplementary figure 3). The Catalan *L. sinapis* male assembly was also compared to the DToL assembly of a *L. sinapis* male from Asturias at the north-western Iberian Peninsula (Lohse et al., 2022). Apart from differences which likely represent genuine rearrangements (see ‘Chromosome rearrangements’), there was a high level of collinearity between these two assemblies (Supplementary figure 4).

**Figure 1.**
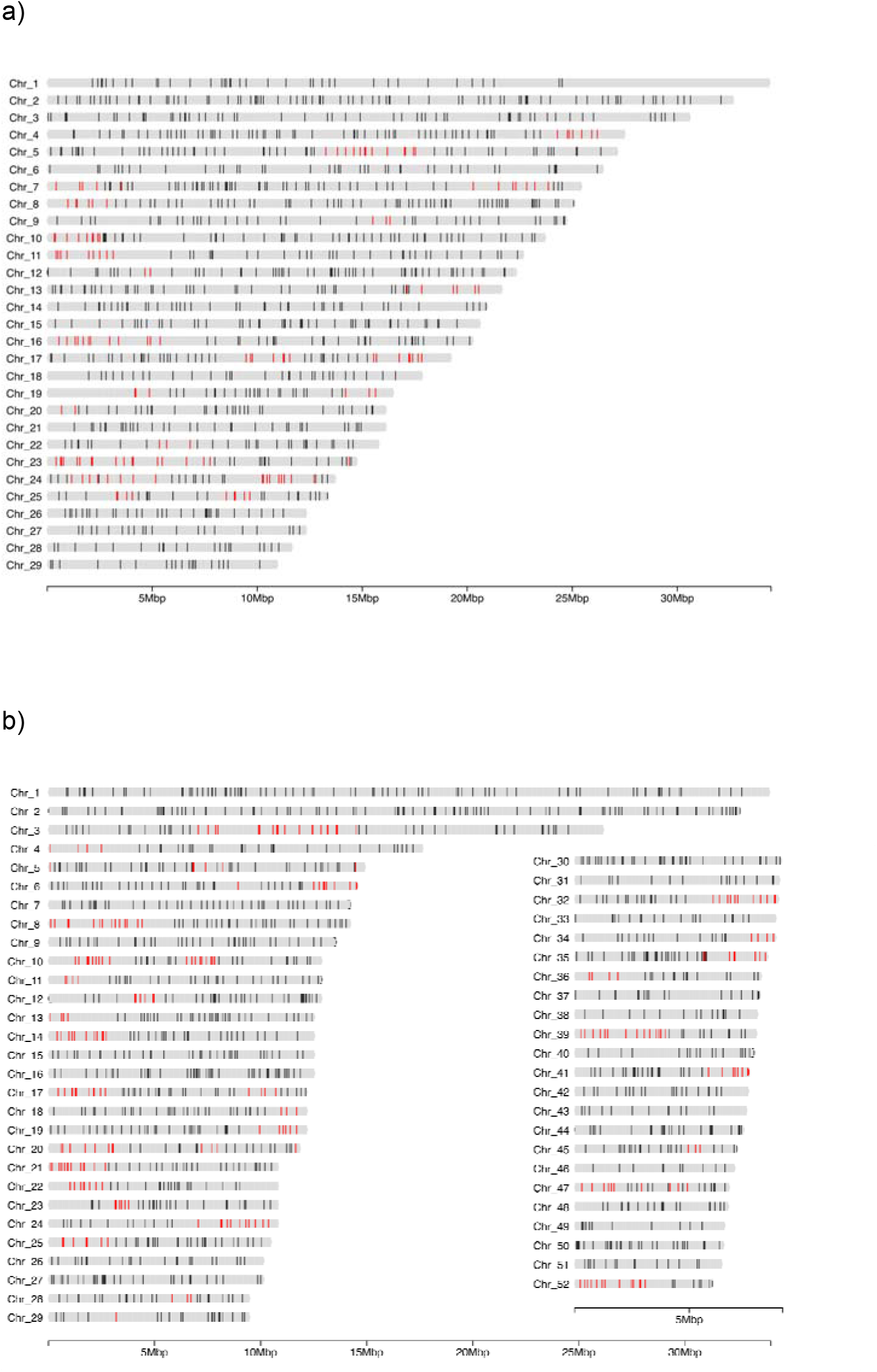
Linkage maps based on pedigrees for a) the Swedish, and b) the Catalan populations of L. sinapis. Vertical bars on each chromosome represent the position of linkage informative markers ordered according to the physical map (genome assembly). The x-axis represents the physical position of each marker in megabases (Mb). Red colour represents markers where the marker order in the linkage map was found to deviate from the order in the preliminary physical assembly.

**Table 1.** Estimated summary statistics for the 8 different *Leptidea* genome assemblies.

| Statistic | *L.*<br>*juvernica*<br>female | *L.*<br>*juvernica*<br>male | *L. reali*<br>female | *L. reali*<br>male | *L. sinapis*<br>Swedish<br>female | *L. sinapis*<br>Swedish<br>male | *L. sinapis*<br>Catalan<br>female | *L. sinapis*<br>Catalan<br>male |
| --- | --- | --- | --- | --- | --- | --- | --- | --- |
| Assembly size (Mb) | 675 | 660 | 670 | 642 | 685 | 664 | 672 | 655 |
| Sequence N's (%) | 3.4% | 4.1% | 3.7% | 2.4% | 5.4% | 5.6% | 3.3% | 3.8% |
| GC (%) | 35.6% | 35.5% | 35.4% | 35.3% | 35.4% | 35.3% | 35.4% | 35.4% |
| Predicted genes (N) | - | 16 149 | - | 15 689 | - | 15 915 | - | 16 478 |
| BUSCO genes (%) | 93.7% | 95.0% | 93.4% | 93.4% | 93.9% | 93.8% | 94.6% | 94.5% |
| Pseudo-chromosomes (N) | 43 | 42 | 26 | 26 | 28 | 29 | 52 | 52 |
| Sequence in Pseudo-chromosomes (%) | 83.3% | 90.7% | 88.0% | 88.4% | 89.2% | 90.2% | 89.2% | 90.4% |
| Read mapping (%) | 98.2% | 98.6% | 98.3% | 98.0% | 98.1% | 98.6% | 98.2% | 98.2% |
| Repeat content (%) | 54.6% | 52.0% | 52.0% | 54.1% | 52.0% | 51.3% | 53.6% | 53.2% |
| Scaffolds (N) | 20 731 | 16 073 | 16 210 | 15 448 | 17 536 | 16 022 | 17 757 | 16 290 |
| Scaffold N50 (Mb) | 15.087 | 15.702 | 22.890 | 21.966 | 24.480 | 21.838 | 11.055 | 11.044 |
| Contigs (N) | 36 909 | 30 088 | 29 081 | 28 137 | 31 643 | 29 387 | 31 619 | 31 643 |
| Contig N50 (kb) | 65.133 | 79.484 | 81.855 | 82.283 | 72.761 | 73.965 | 78.960 | 79.404 |

### Chromosome rearrangements

In order to characterize how chromosomes in the *Leptidea* clade correspond to the presumably nearly ancestral lepidopteran karyotypes of *B. mori* and *M. cinxia*, gene orders were compared between the different lineages (Figure 2). The synteny analysis revealed a considerable amount of large rearrangements where each *Leptidea* chromosome showed homology to 3 - 5 (median) *B. mori* chromosomes (Figure 2, Supplementary figure 5, Supplementary table 2). Not only have all chromosomes in *Leptidea* been restructured compared to the ancestral lepidopteran karyotype, but the frequency of rearrangements between *Leptidea* species was also high. Only one of the sex chromosomes (Z2) showed conserved synteny across all *Leptidea* species (Figure 2). When compared to *B. mori*, the *Leptidea* genomes contained 372 - 410 distinct synteny blocks. The synteny blocks ranged between 0.94 - 1.00 Mb, corresponding to 12.5 - 14.5 blocks per chromosome, and contained 24 - 25 genes (medians given; Supplementary table 2).

**Figure 2.**
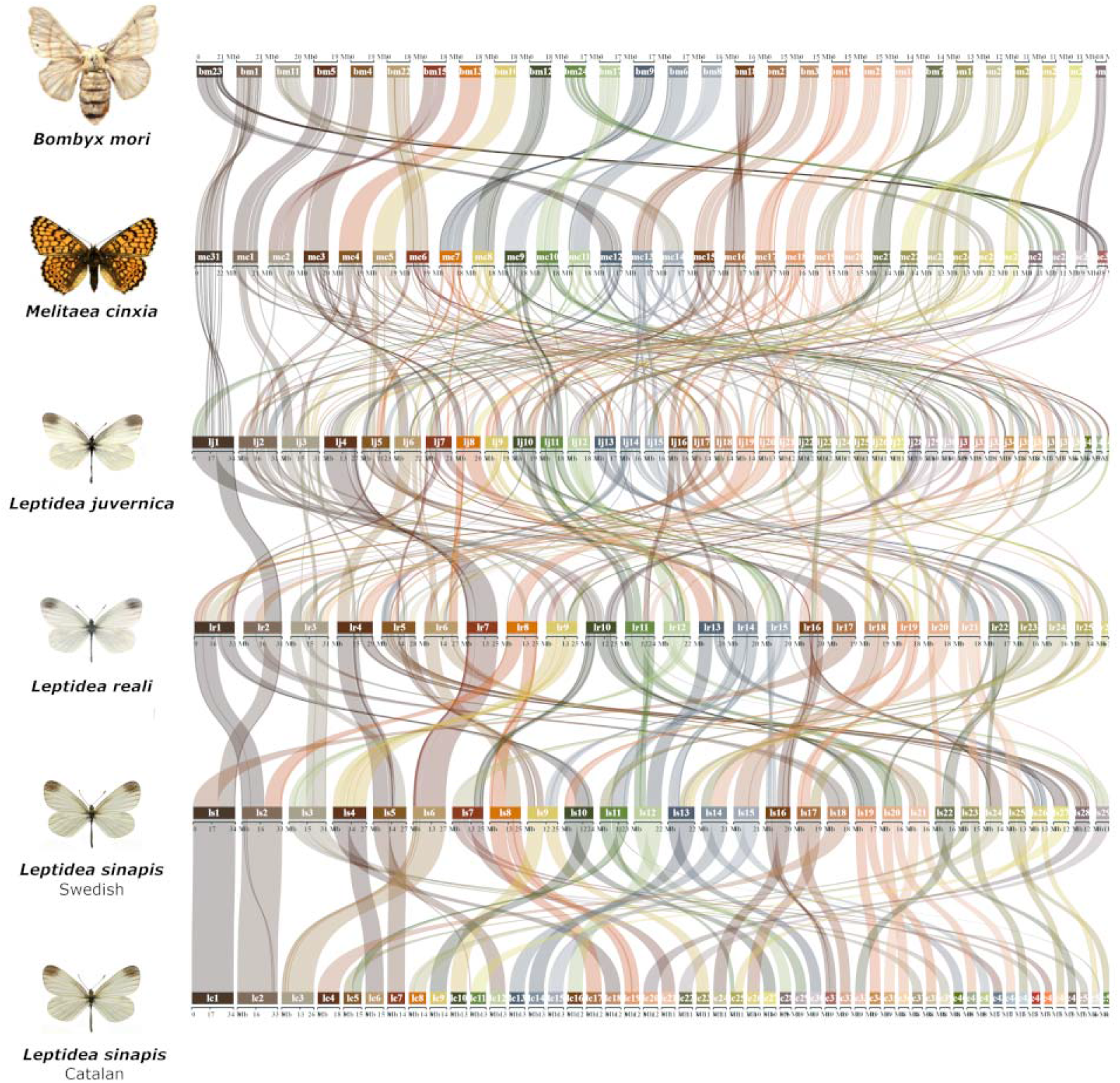
Comparison of synteny between the Leptidea species and the two references B. mori and M. cinxia. The karyotype of the latter species is assumed to represent the ancestral lepidopteran karyotype (Ahola et al., 2014). Chromosomes are ordered by size in each species. Pair-wise comparisons between B. mori and each respective Leptidea lineage are available in Supplementary Figure 5.

Since equivalent large-scale genome restructuring was recently shown in another pierid butterfly, *P. napi* (Hill et al., 2019), we also compared synteny between *P. napi* and the different *Leptidea* species. Apart from one syntenic chromosome pair (*P. napi* chromosome 18 and Catalan *L. sinapis* chromosome 31), all chromosome pairs in the comparisons contained a mix of synteny blocks from at least two different chromosomes, indicating that the absolute majority of rearrangements in *P. napi* and *Leptidea* have occurred independently in the two different lineages (Supplementary figure 6).

After discovering the considerable genomic reorganizations in *Leptidea*, we aimed at characterizing the different rearrangements in more detail. A phylogenetic approach was implemented to infer the most parsimonious scenario for intra-specific chromosome rearrangements. In *L. sinapis*, the data indicate that 9 fusions have occurred in the Swedish population (Supplementary figure 7) and 8 fissions (Supplementary figure 8) and two fusions (Supplementary figure 7) in the Catalan population. In addition, there were 8 cases of shared breakpoints between the Catalan *L. sinapis* and *L. juvernica* (Supplementary figure 8), which explain the majority of the remaining differences in chromosome numbers between Catalan and Swedish *L. sinapis*. Such shared breakpoints could either represent sorting of ancestral fissions, independent fissions at the same chromosome positions in Catalan *L. sinapis* and *L. juvernica*, or independent, identical fusions in *L. reali* and Swedish *L. sinapis*. If fusions tend to occur randomly, we expected that chromosome pairs that have fused independently in two different lineages could have different orientations. However, in all cases except one, where a small inversion was in close proximity to a breakpoint, the chromosomes were collinear between *L. reali* and Swedish *L. sinapis*, suggesting that the ancestral state has been retained in both species. The two *L. sinapis* populations also shared five ancestral fusions (Supplementary figure 7), but no shared fissions. In addition, four of the fissions identified in the Catalan *L. sinapis* were not present in the individual from Asturias (Supplementary figure 4). In *L. reali* we identified 21 fusions (Supplementary figure 9) and six fissions (Supplementary figure 10) in total. We cannot exclude that some of these events correspond to chromosome translocations rather than fissions and fusions, since all except one of the fissioned chromosomes also have been involved in fusions. Finally, in the comparison between *L. juvernica* on the one hand and *L. sinapis* / *L. reali* on the other, we found 51 chromosome breakpoints shared by *L. sinapis* / *L. reali* (fusion in *L. juvernica* or fission in *L. sinapis* / *L. reali*) and 44 unique breakpoints in *L. juvernica* (fission in *L. juvernica* or fusion in *L. sinapis* / *L. reali*).

The extreme rate of chromosomal rearrangements observed within the *Leptidea* clade motivated further comparison to the ancestral Lepidopteran karyotype. We therefore tested how often coordinates of the previously identified chromosome breakpoints overlapped with breaks in synteny between *M. cinxia* and *Leptidea*. Synteny breaks were inferred when consecutive alignment blocks switched from one chromosome to another in *M. cinxia*. This analysis showed that 33 of the 37 fusions previously called in *L. sinapis* and *L. reali* overlapped with synteny breakpoints in *M. cinxia* (33 / 37 overlapping fusions is significantly different from a random genomic occurrence of overlaps; randomization test, p-value < 1.0*10^−5^; Supplementary figure 11), verifying that these constitute novel rearrangements in *Leptidea*. The four exceptions included one fusion in *L. sinapis* and three fusions in *L. reali*. These cases could represent fissions in the ancestral *Leptidea* lineage followed by recurrent fusions in a specific species, or fissions that occurred independently in *L. juvernica* and *M. cinxia*. Three of 14 previously characterized fissions in *Leptidea* and four of 8 chromosome breakpoints shared between *L. juvernica* and Catalan *L. sinapis* overlapped synteny breaks in *M. cinxia* and therefore likely rather represent fusions in a *Leptidea* lineage than independent fissions in *M. cinxia* and one of the *Leptidea* species.

In cases where chromosome rearrangements within *Leptidea* were not supported based on homology information in *M. cinxia*, we analyzed the breakpoint regions for presence of telomeric repeats (TTAGG)n or (CCTAA)n, which should only be present if the rearranged region corresponds to a chromosome fusion. For the inferred fusion shared by both *L. sinapis* populations, we found telomeric repeats in both the Swedish and Catalan individuals, indicating that this represents a fusion back to the ancestral state. Conversely, the three fusions in *L. reali* did not contain any telomeric repeats which indicates that several fissions have occurred in the same chromosome region in different *Leptidea* lineages, or that ancestral fission/fusion polymorphisms have sorted differently within the *Leptidea* clade. Regions around one fission in Catalan *L. sinapis* and two fissions in *L. reali* that were not supported by homology information in *M. cinxia* also lacked telomeric repeats in the other *Leptidea* lineages. These three fissions therefore likely occurred within regions that represent chromosome fusions that pre-date the radiation of the *Leptidea* clade. Finally, two of the four shared fission breakpoints between *L. juvernica* and Catalan *L. sinapis*, which were fusion-like compared to *M. cinxia*, did not contain telomeric repeats in the other *Leptidea* samples and therefore likely represent fissions. All inferred fission and fusion events are summarized in Figure 3 and Supplementary table 3. We further polarized the chromosome breakpoints identified between *L. juvernica* and the other *Leptidea* species. Here, 46 of 51 *L. reali* / *L. sinapis* chromosome breakpoints (fusion in *L. juvernica* or fission in *L. reali* / *L. sinapis*) and 16 of 44 breakpoints in *L. juvernica* (fission in *L. juvernica* or fusion in *L. reali* / *L. sinapis*) had overlapping synteny breaks in *M. cinxia*, indicating that the majority (65.3%) represent fusions that have occurred within the *Leptidea* clade. The directions and frequencies of chromosome rearrangements in the different lineages suggest that the common ancestor of the three investigated *Leptidea* species had a haploid chromosome number (n ~ 51-53) close to that of present day Catalan *L. sinapis*.

**Figure 3.**
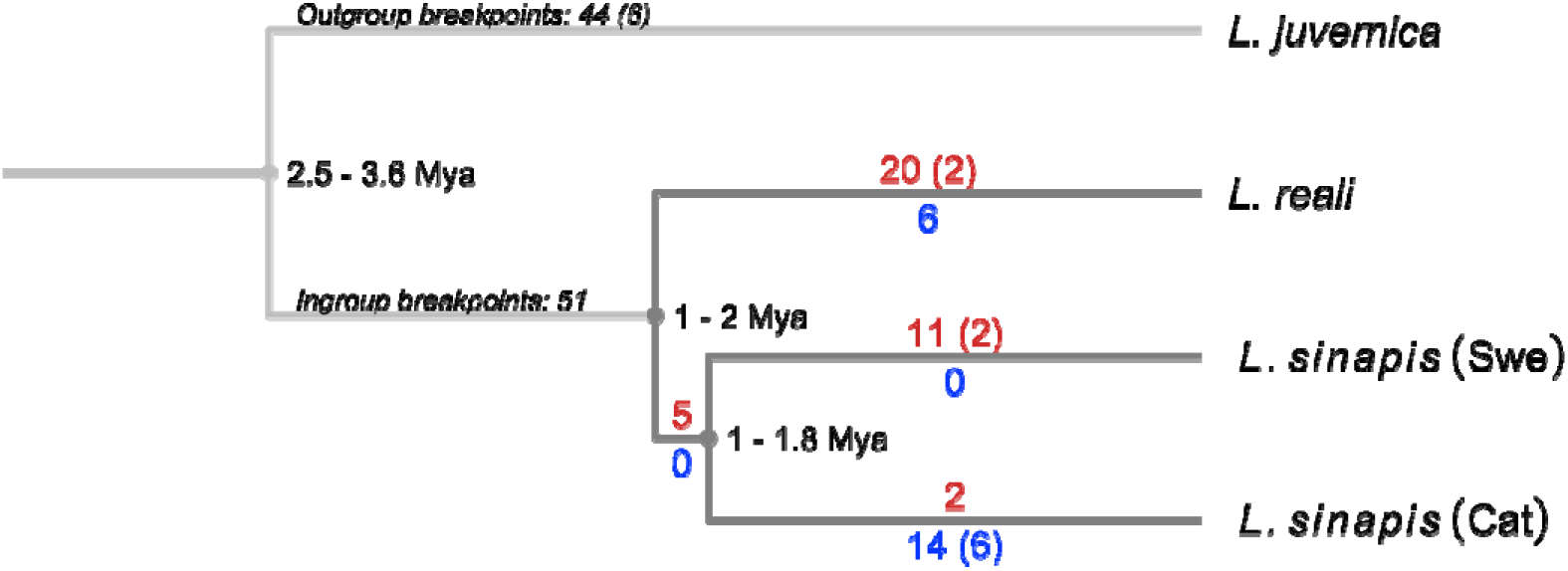
Estimated number of chromosomal rearrangement events in the different Leptidea species/populations. Fusions are highlighted in red and fissions in blue. Numbers show total counts for each branch and shared events are shown in parenthesis. Divergence times are based on Talla et al. (2017).

Next, we estimated how much of the observed rearrangements could alternatively be the result of translocations. First, we checked for exchange of homologous regions between chromosomes and species, which could be an indication of reciprocal translocations. For the ingroup species (*L. sinapis* and *L. reali*) we only detected one such case in Catalan *L. sinapis* involving chromosomes 25 and 51. Next, we counted how often a chromosome region was flanked by regions from one other chromosome in another species which could indicate a non-reciprocal, internal translocation event. Here, we also only detected one case, in *L. reali* chromosome 20. Finally, since translocations do not necessarily affect chromosome ends, we counted how many ancestral chromosome ends (compared to *B. mori*) were still present in *Leptidea* and how many of these were kept as ancestral pairs. This showed that few ancestral chromosome ends are maintained (*L. juvernica:* 10/84, *L. reali:* 7/52, Swedish *L. sinapis*: 11/56, Catalan *L. sinapis*: 18/104) and no ends remain paired in any of the species. Taken together, this suggests that translocations are rare and most rearrangements have occurred through fissions and fusions.

### Intraspecific rearrangements

The high frequency of interspecific chromosome rearrangements between *Leptidea* species spurred an additional set of analysis to assess occurrences and frequencies of fission / fusion polymorphisms segregating within the populations. While the physical assemblies of the male and female *L. reali* and Catalan *L. sinapis*, respectively, were collinear, we found evidence for fission / fusion polymorphisms segregating within both *L. sinapis* and *L. juvernica* (Supplementary figure 12). First, the Catalan (C) and Asturian (A) *L. sinapis* individuals, which presumably represent populations with recent shared ancestry, had highly similar karyotypes, but we found one fusion polymorphism (C 5 = A 6 + A 45; A 6 = C 5 + C 51; Supplementary figure 4) and a potential translocation involving chromosomes C 6 and C 18 + A 10 and A 18, respectively (Supplementary figure 4). Second, within the Swedish *L. sinapis*, the comparison of the male and the female assemblies revealed two segregating chromosome fusions (♀ chromosome 3 = ♂ chromosomes ♂ 19 + ♂ 25, and ♀ 6 = ♂ 21 + ♂ 28) (Supplementary figure 12 and 13) and two cases where the male and female were heterozygous for different rearrangement polymorphisms (♀ 4 = ♂ 5 + ♂ 27; ♂ 5 = ♀ 4 + ♀ 27, and ♀ 10 = ♂ 11 + ♂ 26; ♂ 11 = ♀ 10 + ♀ 26) (Figure 4, Supplementary figure 12). In the first case (♀ 4 = ♂ 5 + ♂ 27; ♂ 5 = ♀ 4 + ♀ 27), the fusion point between chromosomes ♀ 4 and ♂ 5 appears to be associated with a large inversion (supported by the linkage map data, see below), which connects the fused variants in different orientations. To understand the background of these two complex rearrangement polymorphisms, we analyzed the homologous regions in the other *Leptidea* populations / species. For the first case, we found that *L. reali* shared the fusion variant observed in the *L. sinapis* female (i.e. ♀ 4 = ♂ 5 + ♂ 27; likely the ancestral state) while the Catalan and Asturian *L. sinapis* shared the variant observed in the Swedish *L. sinapis* male (♂ 5 = ♀ 4 + ♀ 27), with an additional fission within the inverted region. In *L. juvernica*, the genomic regions involved in rearrangement polymorphisms in the Swedish *L. sinapis* were separate chromosomes. In the second case (♀ 10 = ♂ 11 + ♂ 26; ♂ 11 = ♀ 10 + ♀ 26), both *L. reali* (chromosome 14) and *L. juvernica* (5) shared the variant observed in the Swedish *L. sinapis* male (♂ 11 = ♀ 10 + ♀ 26) with several additional rearrangements around the fusion point. The constitution in Catalan and Asturian *L. sinapis* was also most similar to the variant observed in the Swedish *L. sinapis* male but with additional smaller rearrangements connected to it. Hence, the variant observed in the Swedish *L. sinapis* female (♀ 10 = ♂ 11 + ♂ 26) appears to be specific to this population (Figure 4).

**Figure 4.**
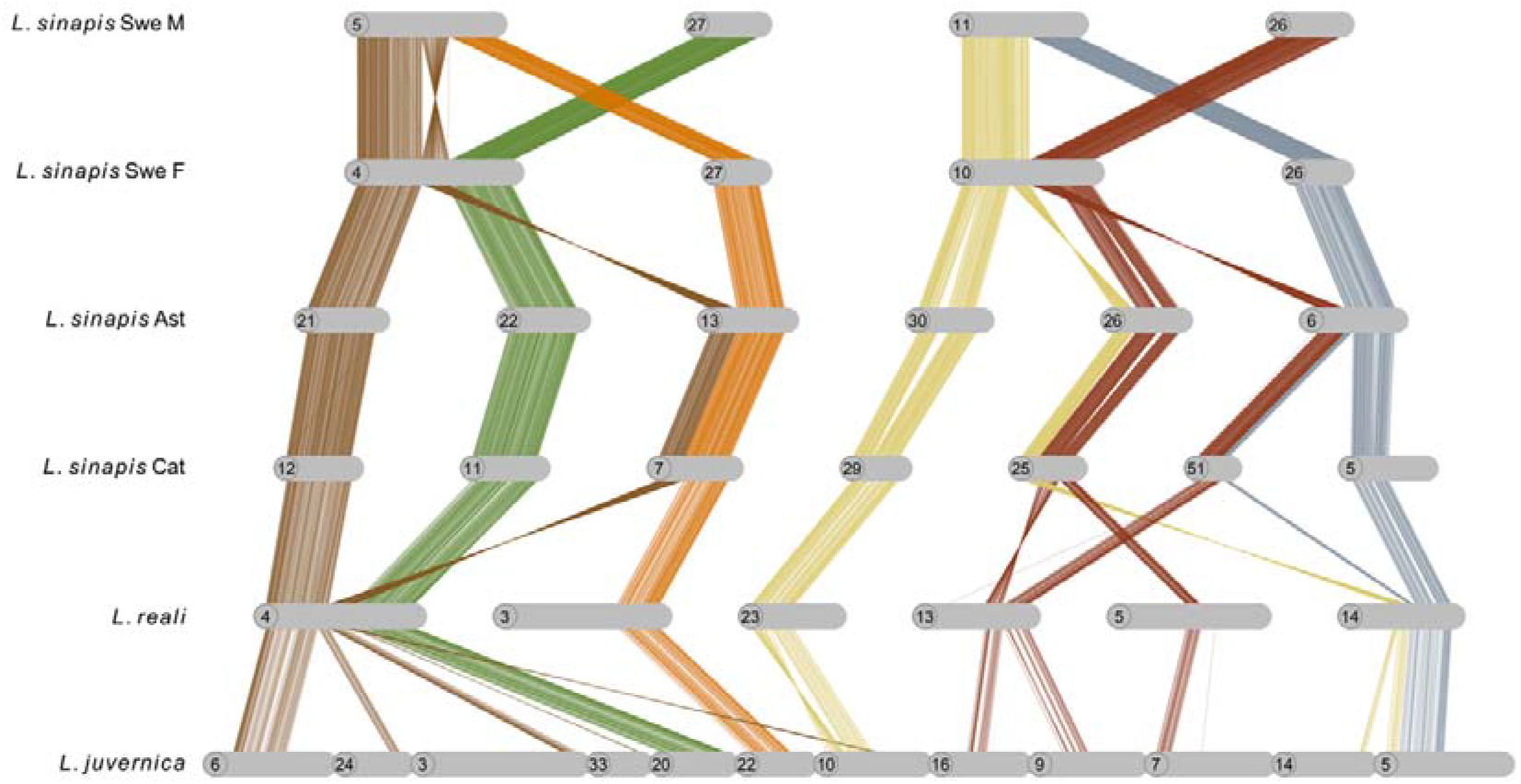
Fusion polymorphisms in Swedish L. sinapis and respective homologous regions in the other Leptidea species and populations. Lines show individual alignments (> 90% similarity) and colours represent homologous regions. Chromosomes have been rotated to enhance visualization. Note that chromosomes 26 and 27 are not homologous to the chromosome with the same number in the opposite sex in Swedish L. sinapis. Ast: Asturias population, Cat: Catalan population.

Finally, when comparing the male and the female *L. juvernica* assemblies, we also detected three segregating fusions (♀ 5 = ♂ 24 + ♂ 25; ♂ 17 = ♀ 28 + ♀ 38; ♂ 41 = ♀ 42 + ♀ 43; supplementary figure 14) and one rearrangement polymorphisms; ♀ 20 = ♂ 27 + ♂ 37; ♂ 27 = ♀ 20 + ♀ 40; Supplementary figure 15).

The intraspecific chromosome rearrangement polymorphisms observed with the HiC-maps obviously only reflects the variation between two individuals in each population. To provide information from more individuals in each population and assess the frequency of segregating rearrangement polymorphisms in more detail, we used the pedigrees to construct single family based linkage maps in the populations with the largest difference in chromosome count - the Swedish and the Catalan *L. sinapis*. This independent analysis verified the observations from the HiC-maps, and revealed additional segregating chromosome rearrangement polymorphisms. For the Swedish population, we could construct linkage maps for four independent families and they all had different karyotypes when compared to the Swedish *L. sinapis* male genome assembly. Family T4 had one of the chromosome fusions (♂ 21 + ♂ 28) and family T3 two additional fusions (♂ 27 + ♂ 5, ♂ 11 + ♂ 26) that were observed when comparing the male and female genome assemblies. In family T5 we observed the same fusions as in family T3 but with additional fissions involving ♂ 7 and ♂ 11. Some of the previously identified fusions and the fission of chromosome ♂ 7 were also observed in family T2. Hence, the linkage analysis in independent families confirmed the fission / fusion polymorphisms identified in the comparison between genome assemblies and revealed an additional fission of ♂ 7 in two families. For the Catalan families we could construct five independent maps. Here we observed three different karyotypes, all differing from the genome assembly of the Catalan *L. sinapis* male. A fission of chromosome ♂ 5 was present in all families, similar to the observation in the Asturian *L. sinapis*. In three families, 3c9, 4C and 9C, we found a fusion of ♂ 6 + ♂ 18 and in addition to ♂ 6 + ♂ 18 there was also one part of ♂ 5 fused to ♂ 45 in family C9. In summary, the genetic maps for independent families provide evidence for several chromosome rearrangement polymorphisms that are currently segregating in the different populations (Figure 5).

**Figure 5.**
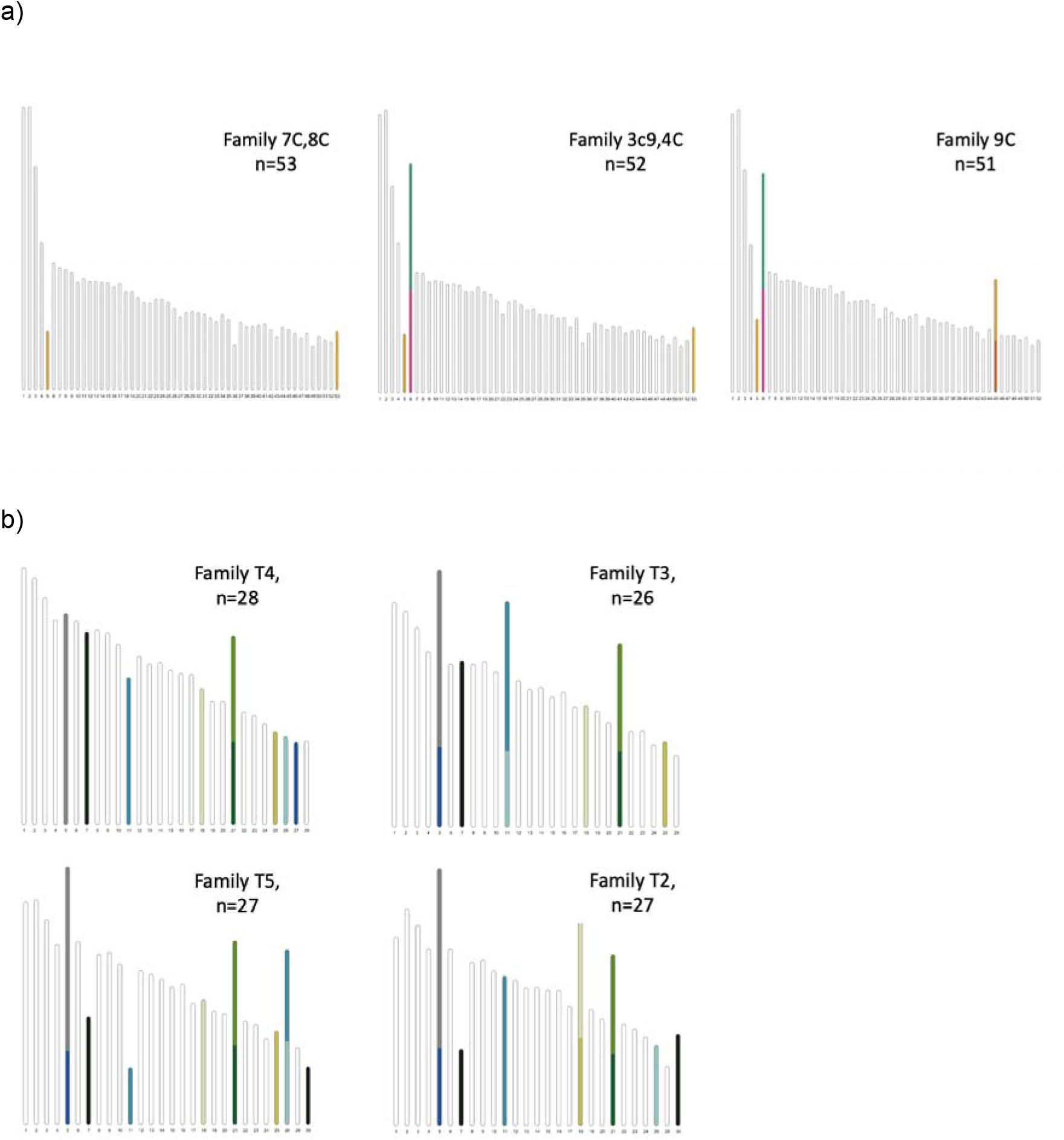
Linkage groups for a) Swedish families and b) Catalan L. sinapis families. Colours represent specific chromosomes in the male genome assembly for the Swedish (n = 26 - 28) and Catalan (n = 51 - 53) L. sinapis population, respectively.

### Structural variation in the sex chromosomes

The synteny of the Z chromosomes agreed with previous results (Yoshido et al., 2020) and in addition we detected previously unknown gene movement from *B. mori* autosomes 19, 26 and 28 to the sex chromosomes in *Leptidea* (Supplementary figure 16). In *L. sinapis*, we identified the three previously described Z chromosomes and a female specific ~ 4.4 Mb long scaffold, which likely represents (at least part of) the W chromosome. No equivalent W scaffold was observed in the Catalan *L. sinapis* female assembly. We noticed however that Z chromosome 3 in the two female *L. sinapis* assemblies and Z chromosome 1 in *L. reali* were several Mb longer than their male homologs and that they aligned less well compared to other chromosomes, indicating that these scaffolds likely are chimeras between the Z and W chromosome parts. In *L. reali*, we also discovered a previously unknown translocation event between what has previously been identified as Z chromosome 1 and Z chromosome 3 (Yoshido et al., 2020; Supplementary figure 16). One of these lineage specific Z chromosomes contains a major part of the ancestral Z which has fused with Z chromosome 3. The other contains the remaining ancestral and neo parts of Z chromosome 1. This was observed in both the male and the female.

### Sequence analysis of the fissioned and fused chromosome regions

In agreement with previous data (Talla et al., 2017), we found that the TE content was > 50% in all *Leptidea* assemblies (Table 1) and the majority of the TEs were long interspersed nuclear elements (LINEs; Supplementary figure 17). Since TEs may facilitate structural rearrangements (Miller & Capy, 2004), we assessed potential associations between specific sequence motifs and rearrangements by estimating the density of different repeat classes and coding sequences in the chromosome regions associated with fusions and fissions and comparing the densities to genomic regions not affected by rearrangements. We limited this analysis to *L. reali* and *L. sinapis* where we could polarize the rearrangements as fissions or fusions. In chromosome regions where fusions have occurred, there was a significant enrichment of both LINEs and LTRs (p-value < 2.0*10^−5^ in both cases), and a significant underrepresentation of SINEs (p-value < 2.0*10^−5^) and rolling-circle TEs (p-value = 1.0*10^−3^). Similarly, there was a significant enrichment of LINEs and a reduction in SINEs and rolling-circles (p < 2.0*10^−5^ in all cases) in chromosome ends in the lineages that lacked the fusion (queries), but in these regions LTRs were not significantly enriched (Figure 6A, Supplementary table 4). This shows that LTRs are more abundant where a fusion has occurred compared to homologous regions in species where a fusion has not taken place. In fission breakpoints in contrast, the only significant difference (p = 0.04) was found for rolling-circles, which were underrepresented in fissioned as compared to non-fissioned chromosomes (Figure 6B, Supplementary table 4).

**Figure 6.**
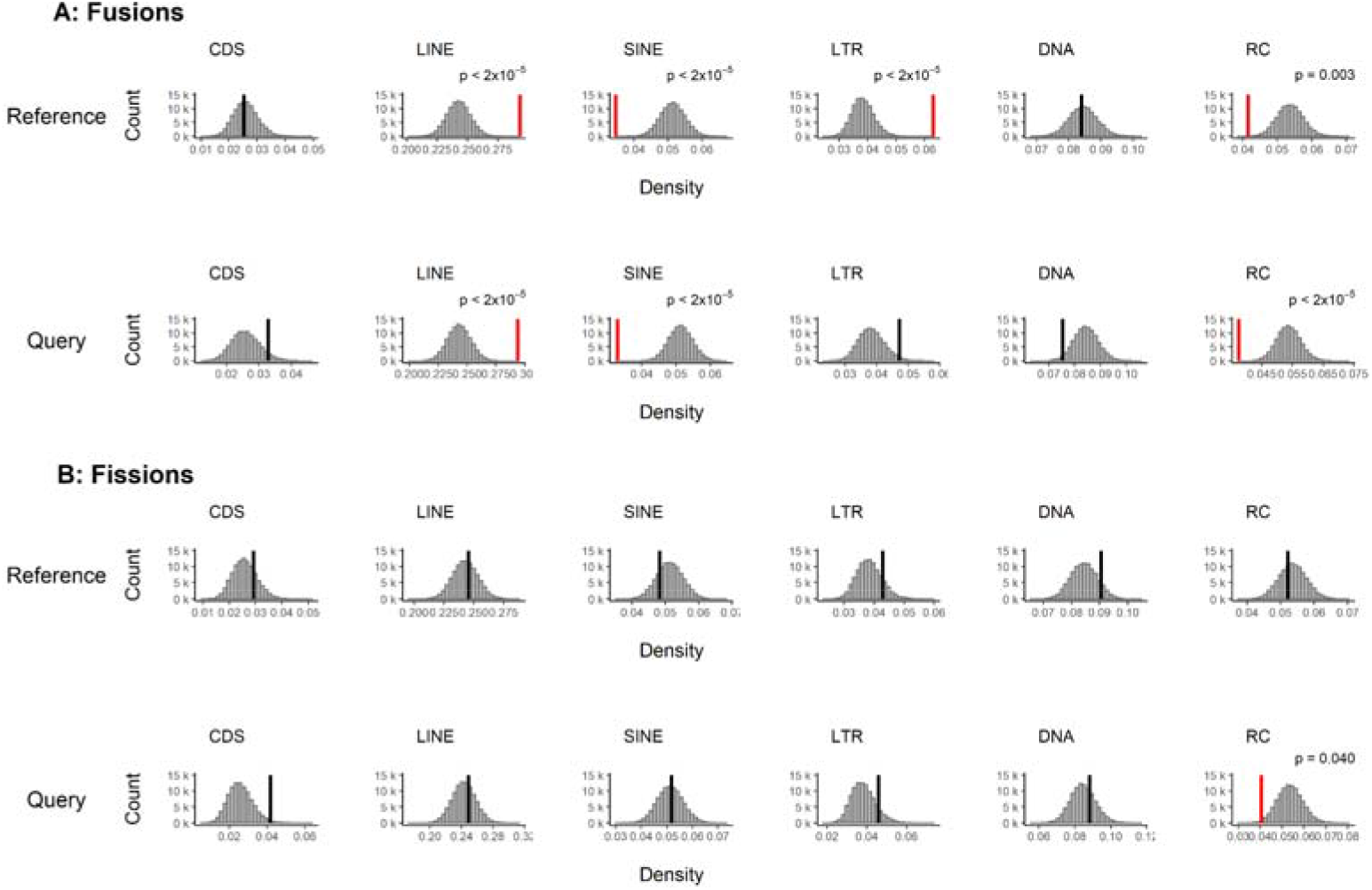
Composition of sequence elements in A) fusion and B) fission breakpoints as compared to the rest of the genomes for L. reali and L. sinapis (the outgroup L. juvernica was excluded from the analysis). Comparisons were performed separately for when the species were used as references or queries. The histograms show distributions of element densities generated by 100 k iterations of random genomic sampling with replacement. Vertical lines show mean density of elements in breakpoints, highlighted in red if significant and black if non-significant. FDR-adjusted p-values are indicated for significant tests. All p-values, means and standard deviations are reported in Supplementary table 4. CDS = coding sequence, LINE = long interspersed elements, SINE = short interspersed elements, LTR = long terminal repeats, DNA = DNA transposons, RC = rolling-circle TEs.

We found that the Asturian *L. sinapis* assembly contained more copies of the short telomeric repeat (TTAGG)n compared to the in-house developed assemblies, and that these were interspersed with certain LINE families. We therefore assessed if increased LINE content in fused chromosome regions could be explained by the presence of telomere associated LINEs. Two classes of telomere associated LINEs had a higher frequency in telomeric regions than in the rest of the genome in the Asturian genome assembly (Fisher’s exact test, fdr-adjusted p-value < 2.0*10^−57^, Supplementary table 5). However, these LINE classes made up only 5.47% of the total LINE content in fused chromosome regions in *Leptidea*, and LINEs in general were still significantly enriched (p-value = 8.0*10^−4^) in those regions after excluding this subset.

## Discussion

Here we present the results from an integrative approach, where we combine genome assembly and annotation with traditional linkage mapping, to characterize and quantify the directions and frequencies of large scale chromosome rearrangements in *Leptidea* butterflies. Our data showed lineage specific patterns of fissions, fusions (and potentially some translocations) and unveiled considerable directional variation in karyomorph change across species and populations. We also identified several segregating fission / fusion polymorphisms in the *Leptidea* populations and characterized specific repeat classes associated with chromosome regions involved in rearrangements. Since the extensive rearrangements have occurred over a comparatively short time scale in *Leptidea* (Talla et al., 2017), the system provides a unique opportunity for investigating the causes and consequences of rapid karyotype change in recently diverged species.

Based on current chromosome number variation within *Leptidea* and the observation that the inferred ancestral karyotype (n = 31) has been conserved across many divergent lepidopteran lineages (de Vos et al., 2020; Robinson, 1971), a straightforward expectation would be that *L. reali* (n = 25 - 28) and the northern populations of *L. sinapis* (n = 28 - 29) have, apart from a few fusion events, mainly retained the ancestral lepidopteran chromosome structures. This would mimic the rearrangements observed for *H. melpomene*, where 10 fusions have reduced the chromosome number (n = 21) compared to the ancestral karyotype (Davey et al., 2016). In line with this reasoning, the higher number of chromosomes in *L. juvernica* (n = 38 - 46) and Iberian *L. sinapis* (n ≈ 53 - 55) could simply be a consequence of chromosomal fissions, as observed in the lycaenid genus *Lysandra* (Pazhenkova & Lukhtanov, 2022). However, analogous to the organization of the genome structure in *P. napi* and *P. rapae* (Hill et al., 2019), our analyses reveal considerably more complex inter- and intra-chromosomal rearrangements in *Leptidea* than anticipated from comparisons of chromosome counts. These results confirm previous findings of a dynamic karyotype evolution in general in the species group (e.g. Dincā et al., 2011; Lukhtanov et al., 2011; Šíchová et al., 2015; Yoshido et al., 2020) and extends them by characterizing the specific chromosome rearrangements in detail and quantifying the differences in fission and fusion rates in different *Leptidea* species and populations. Despite the rather complex patterns of restructuring observed, we found some general trends of karyotype change between the species. For *L. reali* and the Swedish *L. sinapis* population, most species-specific chromosomes have been formed from fusions of chromosomes that segregate independently in Catalan *L. sinapis*. In Catalan *L. sinapis* on the other hand, lineage specific chromosomes have mainly formed through fissions of larger ancestral chromosomes. Since *L. juvernica* was used as an outgroup in the analysis, we could not infer direction in this lineage. However, comparisons with more divergent lepidopteran species suggest that both fissions and fusions have occurred at a high rate also in *L. juvernica*.

Our analyses show that the synteny blocks are short between *Leptidea* and the inferred ancestral karyotype - typically less than 1 Mb, which translates to 12 - 15 blocks per chromosome. Albeit less extensive, such a pattern has also been observed in *Pieris* sp. and has been suggested to reflect a history of recurrent reciprocal chromosome translocations (Hill et al., 2019). The synteny comparison between *Leptidea* species and *B. mori* showed that not a single chromosome in any *Leptidea* species has retained both chromosome ends and that only a minor fraction (~ 12 - 19%) have retained one of the ancestral ends. We also detect few signs of translocations and the high degree of synteny fragmentation in *Leptidea* is therefore probably a consequence of recurrent fissions and fusions in different chromosome regions and between different chromosome pairs, respectively.

The short synteny blocks also show that extensive chromosomal restructuring has occurred in the ancestral lineage of the *Leptidea* species included in our analyses and continued at a high rate in all species. Although we did not have data for all species in the genus, the inferred ancestral karyomorph of the analyzed species (n ~ 50) in combination with the high and variable chromosome numbers in more divergent *Leptidea* species - *L. amurensis* (n = 59 - 61), *L. duponcheli* (n = 102 - 104) and *L. morsei* (n = 54) (Robinson, 1971; Šíchová et al., 2016) - suggest that a high chromosome rearrangement rate is ubiquitous in wood whites. In *L. reali* for example, the chromosome count has decreased to n = 26 in 1 - 2 My since the split from *L. sinapis* (Talla et al., 2017). Even more striking is the rate of change in *L. sinapis* where chromosome counts range from n = 27 - 55 between recently diverged populations. This translates to a chromosome number evolutionary rate in both *L. sinapis* and *L. reali* that has been considerably faster than in for example *H. melpomene*, where 10 chromosome fusions have occurred over the last six million years (Davey et al., 2016). Additional support for an extreme rearrangement rate in the genus comes from the observations that several intraspecific fission / fusion polymorphisms are currently segregating within the different species (see also Lukhtanov et al., 2011; Šíchová et al., 2015) and that incomplete lineage sorting and / or recurrent rearrangements involving the same chromosome regions have been frequent in *Leptidea* historically.

Lepidoptera has traditionally been viewed as having a conserved genomic synteny. However, recent studies (Hill et al., 2019; Yoshido et al., 2020) and the results presented here add some doubt to this view. As mentioned above, an elevated rate of chromosome rearrangements has for example also been observed in the *Pieris napi / rapae* lineage (Hill et al., 2019), which belongs to the same family as *Leptidea* (Pieridae), but the two genera diverged approximately 80 Mya (Espeland et al., 2018) and our synteny analysis clearly show that the rearrangements have occurred independently. Given the limited availability of high-contiguity genome assemblies and / or high-resolution linkage maps, a more holistic view of inter- and intra-chromosomal rearrangement rates will have to await broader taxonomic sampling. Still, we can ask why some lepidopteran taxa are extremely conserved in terms of karyotype and synteny, while other lineages have accumulated a large number of fissions, fusions and translocations, and also why some chromosomes are more conserved than others. Holocentricity and female achiasmy may facilitate segregation and retention of polymorphic chromosomes (Melters et al., 2012), and consequently accelerate genome restructuring. However, a recent phylogenetic overview in insects showed that karyotype evolution is not accelerated in clades with holocentric chromosomes as compared to monocentric, although Lepidoptera appears to be an exception with higher rates of both fissions and fusions (Ruckman et al., 2020). Inverted (post-reductional) meiosis is another mechanism proposed to be important for mitigating the negative effects of chromosomal heterozygosity (Lukhtanov et al., 2018). Within Lepidoptera, this phenomenon has been observed in *L. sinapis* (Lukhtanov et al., 2018) and facultatively in *B. mori* (Banno et al., 1995) and *Polyommatus poseidonides* (Lukhtanov et al., 2020), but because of its association with holocentric chromosomes (Melters et al., 2012) it could potentially be more widespread. However, why do we not observe an elevated rate of chromosome rearrangements in butterflies and moths in general? *Bombyx mori*, for example, has a similar repeat density as *Leptidea* (Tang et al., 2021) and shares the potential for inverted meiosis (Banno et al., 1995). Still *B. mori*, and the majority of lepidopteran taxa with chromosome structure information, have retained the ancestral lepidopteran karyotype (Ahola et al., 2014; Pringle et al., 2007). One option is that chromosome rearrangements are dependent on the presence of specific features that generate *de novo* rearrangement mutations - i.e. that the rate of structural change is mutation limited. Within *Leptidea* for example, a relatively recent burst of transposable element activity has occurred (Talla et al., 2017). This increased activity of certain TE classes could potentially be an important driver of genomic restructuring. We found for example increased LINE and LTR density in fused chromosome regions. Previous data suggest that LINEs make up a considerable portion of the telomere regions in Lepidoptera (Okazaki et al., 1995; Takahashi et al., 1997), but the specific class we could associate with telomeres in *Leptidea* could not explain the general enrichment of LINEs in fused regions. LINEs have previously been associated with rearrangements in monocentric organisms, for example bats (Sotero-Caio et al., 2015) and gibbons (Carbone et al., 2014). Enrichment of LINEs and LTRs was similarly shown to occur in synteny breakpoints within the highly rearranged genome of the aphid *Myzus persicae* (Mathers et al., 2021), although in this case several other classes of TEs were also overrepresented. A plausible explanation is that an increase in ectopic recombination between similar copies of specific TE repeat classes located on different chromosomes (Almojil et al., 2021) can lead to rearrangements, but only in species where these specific classes have proliferated recently. In *Heliconius*, for example, there is a higher density of TEs in general in chromosome fusion points (Cicconardi et al., 2021). Here, we found that the enrichment of LINEs was significant both in species / regions where a chromosome fusion has taken place and in homologous chromosome regions in species where the fusion event has not occurred. This shows that the density of LINEs has been higher in chromosome regions where fusions have occurred rather than accumulating in the regions after the fusion event and indicates that recently proliferated LINE families could play an important role for rearrangements in *Leptidea*. However, we found no association between any of the investigated genomic features and chromosome fission events which indicates that chromosome breakage depends on a mechanism that we could not pick up with our data.

Although *Leptidea* has the most rearranged sex chromosomes of any Lepidopteran species described so far (Yoshido et al., 2020), our synteny analysis showed that the Z chromosomes are considerably more structurally conserved than the autosomes. In particular Z2, which is the only chromosome that has been completely conserved since the split of the *Leptidea* species. The gene content of the ancestral Z chromosome has also been maintained, although the gene order has been highly reshuffled from the ancestral state. A similar situation of sex chromosome conservation in the face of extensive genome restructuring was recently shown in the aphid *M. persicae* (Mathers et al., 2021) and the Z chromosome is highly conserved in Lepidoptera (Fraïsse et al., 2017; Sahara et al., 2012), even in rearranged genomes (Hill et al., 2019), but several cases of fusions with autosomes have been documented (Hill et al., 2019; Mongue et al., 2017; Nguyen et al., 2013). In systems where one sex chromosome is degenerated, for example the W chromosomes in lizards (Iannucci et al., 2019), snakes (Rovatsos et al., 2015) and butterflies (Lewis et al., 2021), the other sex-chromosome (here Z-chromosomes) is often highly conserved. One potential explanation for this is that translocation of genes with male-biased expression from the Z chromosome to an autosome likely would have deleterious effects in females (Vicoso, 2019). It has also been proposed that selection for maintaining linkage of genes with sex-biased expression can be a strong stabilizing force, as seen for example in birds (Nanda et al., 2008). Accumulation of male biased genes on the Z chromosome, as has been observed in many lepidopteran species (Arunkumar et al., 2009; Mongue & Walters, 2018) including *L. sinapis* (Höök et al., 2019), can therefore be a potential reason for the much more conserved Z chromosomes. The fact that the Z chromosome only recombines and spends relatively more time in male butterflies (Turner & Sheppard, 1975) should further strengthen this linkage. Translocation of genes from the sex chromosomes might also be selected against if it alters expression levels regulated by dosage compensation mechanisms, which has been observed in *L. sinapis* (Höök et al., 2019).

## Conclusion

Here we present female and male genome assemblies for three different *Leptidea* species and develop detailed linkage maps for two populations of *L. sinapis*. Synteny analysis revealed one of the most dramatic and rapid cases of chromosome evolution presented so far. The genus not only has one of the most variable intra- and interspecific chromosome numbers, but also, as shown here, potentially the most rearranged genomes across Lepidoptera. Our data suggests that fissions and fusions have been the main cause of the restructuring and that several rearrangement polymorphisms still segregate in the different species and populations. We further find an association between LINEs and LTR elements and fusion breakpoints which should be explored in more depth in future studies. The results presented here add another example of extensive genome reshuffling in Lepidoptera, which shows that the karyomorph does not necessarily predict the extent of chromosome rearrangements in a species.

## Methods

### Samples

Mated adult females of *L. sinapis* (Sweden and Catalonia), *L. reali* (Catalonia) and *L. juvernica* (Sweden) were sampled in the field 2019 and kept in the lab for egg laying. One male and one female offspring from each dam were sampled at the chrysalis stage and flash frozen in liquid nitrogen. Each sample was divided in two aliquots to allow for generation of both a 10X Genomics Chromium Genome- and a Dovetail HiC-library from each individual. For 10X sequencing, DNA was extracted using a modified high molecular weight salt extraction method (Aljanabi & Martinez, 1997). Tissues were homogenized for HiC sequencing using a mortar and pestle in liquid nitrogen and sent to the National Genomics Infrastructure (NGI, Stockholm) for library preparation and sequencing.

### Sequencing and assembly

Library preparations, sequencing and genome assembly was performed by NGI Stockholm using the Illumina NovaSeq6000 technology with 2 x 151 bp read length. 10X linked reads were assembled with 10X Genomics Supernova v. 2.1.0 (Weisenfeld et al., 2017). HiC reads were processed with Juicer v. 1.6 (Durand et al., 2016a) and assemblies were scaffolded with 3DDNA v.180922 (Dudchenko et al., 2017). Resulting assemblies were corrected in several consecutive steps with Juicebox v. 1.11.08 (Durand et al., 2016b). First, all obvious scaffolding errors were corrected using the HiC contact information. Next, linkage information (see subheading “Linkage maps”) was used to identify and correct technical inversions and translocations in the *L. sinapis* assemblies. Finally, we used pairwise alignments (see Genome alignments) between i) the male and female from each respective population, and ii) all assemblies, including an assembly of an Asturian *L. sinapis* individual from the Darwin Tree of Life (DToL) initiative (Lohse et al., 2022), and used visual inspection to detect deviating scaffold orders or orientations. Manual corrections were done in cases when the orientation was not in conflict with the HiC-signal (Supplementary figure 1). After manual curation, the number of chromosome sized scaffolds for each assembly (Swedish *L. sinapis* n = 28 - 29, Catalan *L. sinapis* n = 52, *L. reali* n = 26, L. juvernica n = 42 - 43) was within the expected karyotype range (Dincā et al., 2011; Lukhtanov et al., 2011; Šíchová et al., 2015) and these scaffolds contained the majority of the total sequence content (Table 1, Supplementary figure 1 and 2). In addition, collinearity between male and female assemblies improved after manual curation (Supplementary figure 12). Redundant scaffolds (100% identical duplicates and scaffolds contained within others) were removed using ‘dedupe.sh’ in BBTools v. 38.61b (Bushnell, 2019). The core gene completeness of the different assemblies was assessed with BUSCO v. 3.0.2 (Simão et al., 2015) using the insecta_odb9 data set. The percent of complete BUSCO insect orthologs ranged between 93.4% in *L. reali* male to 95.0% in *L. juvernica* male (Table 1, Supplementary table 1). Potential contaminant scaffolds were identified and removed using BlobTools v. 1.1.1 (Laetsch & Blaxter, 2017).

#### Linkage map

##### Sampling, DNA-extraction and sequencing protocol

Offspring from wild caught females from two populations of *L. sinapis* with different karyotypes (Sweden n = 28 - 29; Catalonia n = 53 - 55) were reared on cuttings of *Lotus corniculatus*. The pedigrees consisted of 6 dams and 184 and 178 offspring from the Swedish and Catalan population, respectively (Supplementary table 6). The offspring were sampled at larval instar V, snap frozen in liquid nitrogen and stored at −20°C. DNA-extractions from caudal abdominal segments were performed with a modified Phenol-Chloroform extraction (batch1; Sambrook & Russell, 2006) or high salt extraction (batch2; Aljanabi & Martinez, 1997) after standard Proteinase-K digestion overnight. The amount and quality of the DNA was analyzed with Nanodrop (Thermo Fisher Scientific) and Qubit (Thermo Fisher Scientific). The DNA was digested with the *EcoR1* enzyme according to the manufacturer’s protocol, using 16 hours digestion time (Thermo Fisher Scientific). The efficiency of the digestion was determined by visual inspection of the fragmentation using gel electrophoresis (1% agarose gel). The fragmented DNA samples were sent for RAD-seq library preparation and paired-end sequencing on Illumina HiSeq2500 (batch 1) and NovaSeq600 (batch 2) at the National Genomics Infrastructure, SciLife, Stockholm.

##### Data processing

The quality of the raw reads was initially assessed with FastQC (Andrews, 2010). Duplicate removal and quality filtering were performed with clone_filter and process_radtags from Stacks2 (Catchen et al., 2013). We applied the quality filtering options -q to filter out reads with > phred score 10 (90% probability of correct base called) in windows 15% of the length of the read, -c to remove all reads with unassigned bases, and --disable_rad_chec to keep reads without complete RAD-tags. All reads were truncated to 120 bp. The filtered reads were mapped to the male genome assembly from each population using bwa mem with default options, and sorted with samtools sort (Li et al., 2009). The bam-files were further filtered with samtools view –q 30 option and a custom script to only retain reads with unique mapping positions. The mapping coverage was analyzed with Qualimap (Okonechnikov et al., 2016) and individual coverage was visualized as mean coverage per chromosome divided by mean coverage per individual. The sex of offspring was set as female if the normalized coverage on the Z1- and Z2-chromosomes was < 75% compared to the autosomes. We used Samtools mpileup for variant calling with minimum mapping quality (-q) 10 and minimum base quality (-Q) 10 (Li, 2011). The variants were converted to likelihoods using Pileup2Likelihoods in LepMap3 with default settings (Rastas, 2017), minimum coverage 3 per individual (minCoverage = 3) and 30% of the individuals allowed to have lower coverage than minimum coverage (numLowerCoverage = 0.3). To verify that the pedigree was correct, the relatedness coefficients of the samples were estimated with the module IBD in LepMap3 using 10% of the markers and a multiple dimensional scaling/principal coordinate analysis based on a distance matrix inferred with Tassel 5 v. 20210210 (Bradbury et al., 2007).

##### Linkage map construction

LepMap3 was used to construct the linkage maps (Rastas, 2017). Informative parental markers were called with module ParentCall using default values, except that non-informative markers were remove and we applied the setting zLimit = 2 to detect markers segregating as sex chromosomes. This module also uses genotype likelihood information from the offspring to impute missing or erroneous parental markers. Markers that did not map to the chromosome-size scaffolds in the physical assembly were removed. The module Filtering2 was applied to remove markers with high segregation distortion (dataTolerance = 0.00001) and markers that were missing in more than 30% of the individuals in each family (missingLimit = 0.3). In addition, only markers present in at least five families were retained (familyInformativeLimit = 5). The markers were binned over stretches of 10 kb with a custom script and binned markers with more than five SNPs per bin were removed. The module OrderMarkers2 with the option outputPhasedData = 4 was used to phase all binned data. The final binning and filtering resulted in 3,237 and 4,207 retained markers in the Swedish and Catalan pedigree, respectively.

The markers were assigned to linkage groups using the module SeparateChromosomes2, with an empirically estimated lodLimit. The lodLimit is the threshold for the logarithm of the odds that two markers are inherited together (LOD-score), i.e. belonging to the same linkage group. We evaluated a range of lodLimits (1 - 30) and finally set it to 10 for the Catalan and 12 for the Swedish families - settings that resulted in approximately the number of linkage groups expected from karyotype data (50 linkage groups in the Catalan and 23 linkage groups in the Swedish families, respectively). To assign additional unlinked markers to the linkage groups, JoinSingles was run with lodLimits 5 for the Catalan map and 8 for the Swedish. Since butterflies have female heterogamety and female achiasmy, the three different Z-chromosomes were clustered in one linkage group and had to be split in separate linkage groups manually (based on information from the physical genome assemblies). To correct for interference and multiple recombination events per linkage group we applied the Kosambi correction. To account for female achiasmy, the recombination rate in females was set to zero (recombination2 = 0). The linkage maps were refined by manually removing non-informative markers at the ends of each map. The trimmed map was re-evaluated with OrderMarkers with the options evaluateOrder and improveOrder = 1. Remaining unlinked markers at map ends were manually removed after visual inspection and the maps were once again re-evaluated with OrderMarkers. The genetic distances and marker orders were compared to the physical positions along each chromosome and the physical coordinates for potential rearrangements were used for re-evaluation of the HiC-maps (Figure 1). The collinearity of the genetic and physical maps was assessed using Spearman’s rank correlation.

##### Read mapping

10X raw reads were first processed with Long ranger basic v. 2.2.2 (Marks et al., 2019). Reads were then trimmed for low quality bases and adapters with Cutadapt v. 2.3 (Martin, 2011) in TrimGalore v. 0.6.1 (Krueger, 2019) using the NovaSeq filter (--nextseq 30) and discarding trimmed reads shorter than 30 bp. Fastqscreen v. 0.11.1 (Wingett & Andrews, 2018) was used to screen and filter libraries from common contaminants (*A. thaliana*, *D. melanogaster*, *E. coli, S. cerevisiae, H. sapiens, C. familiaris, M. musculus, Wolbachia* and *L. corniculatus*, downloaded from NCBI 2021-03-05). Reads were mapped with BWA mem v. 0.7.17 (Li, 2013) resulting in a high mapping rate across all assemblies (98.0 - 98.6 %; Table 1). Mapped reads were then filtered for supplementary and secondary alignments with a custom script, and low-quality alignments (mapq < 30) with samtools v1.10 (Li et al., 2009). Duplicate reads were removed using MarkDuplicates in GATK v. 4.1.1.0 (McKenna et al., 2010). Resulting read mappings were assessed with samtools flagstats (Li et al., 2009) and QualiMap v. 2.2 (Okonechnikov et al., 2016).

##### MtDNA

Circularized mitochondrial genomes for each sample were assembled de novo from processed 10X reads using NOVOplasty v. 4.2 (Dierckxsens et al., 2017) with the *COX1* gene from *Leptidea morsei* as seed (downloaded from NCBI 2020-08-29). The mitochondrial genomes were annotated with MITOS (Bernt et al., 2013), using custom scripts to set the gene *TRNM* as starting position. To identify and remove partial mtDNA scaffolds from the nuclear genome assemblies, we aligned the mtDNA genomes to the assemblies with nucmer in MUMmer v. 4.0.0rc1 (Marçais et al., 2018) using default settings and filtering the output with delta-filter -g. Scaffolds aligning by > 95% of their length and with > 95% identity were filtered out using a custom script. After filtering away identified partial mtDNA scaffolds we reintroduced the mtDNA as a separate scaffold to each assembly.

##### Sex chromosome identification

Sex chromosomes were identified by homology and read coverage. We used gene synteny (see synteny analysis) between *B. mori* and the sequenced *Leptidea* species to verify previously characterized sex and neo-sex chromosomes (Yoshido et al., 2020). As an independent validation we analyzed read depth (see read mapping) for chromosome sized scaffolds across the female and male assemblies with QualiMap v. 2.2 (Okonechnikov et al., 2016). For scaffolds identified as sex chromosomes, the mean read coverage in females ranged between 49.36 - 74.44% of mean assembly coverage (Supplementary table 7, Supplementary figure 18). Sex chromosomes were also verified visually in non-normalized HiC heatmaps, where female samples had a clearly reduced contact between identified sex chromosomes and autosomes (Supplementary figure 1) as compared to the background noise.

##### Gene and repeat annotation

Repeat libraries were generated *de novo* with RepeatModeler v. 1.0.11 (Smit & Hubley, 2017) using default settings. Repeat families classified as unknown by RepeatModeler were additionally screened against Repbase (Jurka, 1998) using CENSOR (Kohany et al., 2006) with the options ‘sequence source - Eukaryota’ and ‘report simple repeat’. The highest scoring hit for each query was integrated with the RepeatModeler output using custom scripts. Repeats were then annotated and quantified using RepeatMasker v. 4.1.0 (Smit et al., 2019). Genes were annotated for the male genome assemblies of each species with the MAKER pipeline v. 3.01.04 (Cantarel et al., 2008). First, both translated protein and coding nucleotide sequences were used as input evidence. Protein sequences were a combination of previous *L. sinapis* annotations (Talla et al., 2019), reviewed Lepidoptera proteins from uniprot (downloaded 2021-04-02) and a set of Lepidoptera core orthologs from Kawahara & Breinholt, 2014. Coding sequences were a combination of *L. sinapis* (Talla et al., 2019) and *L. juvernica* (Yoshido et al., 2020) transcripts. In addition, the output of RepeatModeler was used to mask repeat sequences within MAKER. Resulting gene models were used to train *ab initio* gene predictors in snap (Korf, 2004) and Augustus v. 3.4.0 (Stanke et al., 2008). The gff-file generated in the first round and the gene predictors were then used jointly in a second round of gene prediction. The number of resulting annotations ranged between 15 689 - 17 229 and were of expected quality and size (Supplementary table 8, Supplementary figures 19 and 20). The final gene models were functionally annotated with Interproscan v. 5.30-69.0 (Jones et al., 2014) and blast searches against Swiss-Prot.

##### Synteny analysis

Gene synteny was compared between the male assemblies from each of the sequenced *Leptidea* species and the two reference species *Bombyx mori* and *Melitaea cinxia*, and in addition *Pieris napi* (assemblies and annotations downloaded from NCBI 2021-04-25). First, reciprocal protein alignments were generated with blastp. The blast output was then trimmed to include the top five best hits per query. Finally, synteny blocks were built with MCScanX (Wang et al., 2012) using default settings but restricting the maximum gene gap size to 10 to reduce the amount of overlapping synteny blocks. The results were visualized with Synvisio (Bandi & Gutwin, 2020) and Circos v. 0.69-9 (Krzywinski et al., 2009).

##### Genome alignments

To guide manual curation, estimate collinearity and detect rearrangements, all assemblies were aligned to the male assembly of each respective species with nucmer in MUMmer v. 4.0.0rc1 (Marçais et al., 2018), using default settings. Alignments were restricted to the chromosome sized scaffolds of each assembly. The resulting alignments were filtered with delta-filter −1 to get 1-to-1 alignments including rearrangements. Male to female alignments for each species were visualized with dotplots using a modified version of the script ‘mummerCoordsDotPlotly.R’ from dotPlotly (Poorten, 2018).

##### Rearrangement analysis

Large scale rearrangements were identified from breakpoints in alignments, when queries changed from aligning against one reference scaffold to another. First, blocks of consecutive alignments (> 90% similarity) between the homologous scaffolds were built using a custom script, removing singleton queries against other scaffolds and only keeping blocks > 100 kb. Rearrangements were then classified using a phylogenetic approach, by finding unique or shared breakpoints between assemblies when aligned against the same reference using bedtools v. 2.29.2 (Quinlan & Hall, 2010). Each male assembly was separately used as reference, complemented by the alignment of the female assembly of the same species as an additional control for interspecific variation. Unique breakpoints were called as fissions in the query species while breakpoints shared by all query species were called as fusions in the reference. Following the same logic, we searched for shared breakpoints between all possible combinations of species pairs to find potential cases of incomplete lineage sorting, reuse of breakpoints or rearrangements in ancestral *L. sinapis* (shared by both populations). The output of each separate comparison was checked and manually curated for any discrepancies resulting from using different references. In addition, based on the identified breakpoints, we quantified potential translocations as any case where two chromosomes in one species contained the same combination of parts from two chromosomes in another species (reciprocal) and cases where one chromosome had one alignment block flanked by two blocks from one other chromosome in another species (non-reciprocal).

##### Breakpoint content analysis

The composition of genetic elements in chromosome breakpoints was analyzed by comparing the observed mean density (here defined as proportion of base pairs) of coding sequence and different transposable element (TE) classes to random resampling distributions taken from the rest of the genomes. TEs from the same category and with overlapping coordinates were merged before analysis. Resampling (with replacement, 100 k iterations) was performed by taking random non-overlapping windows of the same size and number as identified fusions or fissions, and calculating the mean density of sequence elements across the sampled windows. The empirical means were compared to the generated resampling distributions using a two-tailed significance test. The false discovery rate was controlled for by adjusting p-values with the Benjamini-Hochberg method (Benjamini & Hochberg, 1995). Fusion and fission breakpoints were analyzed separately, only including events identified in the ingroup species (*L. sinapis* and *L. reali*) and excluding cases with ambiguous polarity. To account for variation in breakpoint size, breakpoints were standardized to 100 kb around their midpoint. In addition, separate tests were made for corresponding homologous regions in the alignment queries. Since there is no shared midpoint coordinate between two query chromosomes flanking a breakpoint, we instead selected two 50 kb windows starting from the last query coordinate of each alignment block and extending towards a theoretical point of breakage. The terminal 50 kb were selected in cases where windows extended beyond the end of the query scaffold. Since the same set of query fissions were scored against two references and not necessarily overlapped, we selected the breakpoints with the coordinates closest to the terminal ends. Similarly, when the same chromosome end was associated with two different fusion events, we selected the outermost coordinates to represent the region. When analyzing query sequences, we excluded internal breakpoints, defined as occurring > 1 Mb from the chromosome terminal ends, as these cases indicate either fissions followed by subsequent fusions (not necessarily representative of a fission) or that different fusions have occurred in the reference and the query and are therefore already counted when each respective species is used as reference. The analysis was performed using bedtools v2.29.2 (Quinlan & Hall, 2010) and custom scripts.

Fusion breakpoints were investigated for the presence and accumulation of telomere associated LINE elements. Putative telomeric LINEs were first identified in the terminal 250 kb of scaffolds of the DToL *L. sinapis* assembly (Lohse et al., 2022). Enrichment of specific LINEs in telomeric regions were then called with Fisher’s exact test using an alpha level of 0.05 and adjusting p-values with the Benjamini-Hochberg method (Benjamini & Hochberg, 1995), and requiring at least the same count of each element in telomeric regions as the number of telomeres (n = 96) and presence in telomeric regions of at least half of the chromosomes. Homologous LINEs in the other assemblies (Swedish + Catalan *L. sinapis* and *L. reali*) were then identified with reciprocal blast alignments, keeping all hits as putative telomere specific LINEs. Finally, accumulation was quantified by calculating the summed fraction of putative telomeric LINEs out of the total LINE density in fusion breakpoints. All statistical tests were performed in R (R Core Team, 2019) unless otherwise noted.

## Data availability

All raw sequence data have been deposited at the European Nucleotide Archive under accession ENAXXXX. All in-house developed scripts and pipelines are available in GitHub (https://github.com/EBC-butterfly-genomics-team).

## Acknowledgements

This work was funded by the Swedish Research Council (VR research grant #019-04791 to N.B.). The authors acknowledge support from the National Genomics Infrastructure in Stockholm funded by Science for Life Laboratory, the Knut and Alice Wallenberg Foundation and the Swedish Research Council, and SNIC/Uppsala Multidisciplinary Center for Advanced Computational Science for assistance with massively parallel sequencing and access to the UPPMAX computational infrastructure. We also acknowledge SciLifeLab in Uppsala for long-term bioinformatics support via the WABI initiative. R.V. was supported by project PID2019-107078GB-100, funded by Ministerio de Ciencia e Innovación (MCIN)/Agencia Estatal de Investigación (AEI)/ 10.13039/501100011033.

## References

Ahola, V., Lehtonen, R., Somervuo, P., Salmela, L., Koskinen, P., Rastas, P., Välimäki, N., Paulin, L., Kvist, J., Wahlberg, N., Tanskanen, J., Hornett, E. A., Ferguson, L. C., Luo, S., Cao, Z., de Jong, M. A., Duplouy, A., Smolander, O.-P., Vogel, H., … Hanski, I. (2014). The Glanville fritillary genome retains an ancient karyotype and reveals selective chromosomal fusions in Lepidoptera. Nature Communications, 5(1), 4737. https://doi.org/10.1038/ncomms5737

Aljanabi, S. M., & Martinez, I. (1997). Universal and rapid salt-extraction of high quality genomic DNA for PCR-based techniques. Nucleic Acids Research, 25(22), 4692–4693. https://doi.org/10.1093/nar/25.22.4692

Almojil, D., Bourgeois, Y., Falis, M., Hariyani, I., Wilcox, J., & Boissinot, S. (2021). The Structural, Functional and Evolutionary Impact of Transposable Elements in Eukaryotes. Genes, 12(6), 918. https://doi.org/10.3390/genes12060918

Andrews, S. (2010). Babraham Bioinformatics—FastQC A Quality Control tool for High Throughput Sequence Data. https://www.bioinformatics.babraham.ac.uk/projects/fastqc/

Arunkumar, K. P., Mita, K., & Nagaraju, J. (2009). The Silkworm Z Chromosome Is Enriched in Testis-Specific Genes. Genetics, 182(2), 493–501. https://doi.org/10.1534/genetics.108.099994

Bandi, V., & Gutwin, C. (n.d.). Interactive Exploration of Genomic Conservation. 10.

Banno, Y., Kawaguchi, Y., Koga, K., & Doira, H. (1995). Postreductional meiosis revealed in males of the mutant with chromosomal aberration “T (23;25) Nd” of the silkworm Bombyx mori. The Journal of Sericultural Science of Japan, 64(5), 410–414. https://doi.org/10.11416/kontyushigen1930.64.410

Belyayev, A. (2014). Bursts of transposable elements as an evolutionary driving force. Journal of Evolutionary Biology, 27(12), 2537–2584. https://doi.org/10.1111/jeb.12513

Benjamini, Y., & Hochberg, Y. (1995). Controlling the False Discovery Rate: A Practical and Powerful Approach to Multiple Testing. Journal of the Royal Statistical Society. Series B (Methodological), 57(1), 289–300.

Bernt, M., Donath, A., Jühling, F., Externbrink, F., Florentz, C., Fritzsch, G., Pütz, J., Middendorf, M., & Stadler, P. F. (2013). MITOS: Improved de novo metazoan mitochondrial genome annotation. Molecular Phylogenetics and Evolution, 69(2), 313–319. https://doi.org/10.1016/j.ympev.2012.08.023

Blackmon, H., Justison, J., Mayrose, I., & Goldberg, E. E. (2019). Meiotic drive shapes rates of karyotype evolution in mammals. Evolution, 73(3), 511–523. https://doi.org/10.1111/evo.13682

Boggs, C. L., Watt, W. B., Ehrlich, P. R., Ehrlich, P. R., & Ehrlich, P. R. (2003). Butterflies: Ecology and Evolution Taking Flight. University of Chicago Press.

Bradbury, P. J., Zhang, Z., Kroon, D. E., Casstevens, T. M., Ramdoss, Y., & Buckler, E. S. (2007). TASSEL: Software for association mapping of complex traits in diverse samples. Bioinformatics, 23(19), 2633–2635. https://doi.org/10.1093/bioinformatics/btm308

Brown, K. S., Jr, Von Schoultz, B., & Suomalainen, E. (2004). Chromosome evolution in Neotropical Danainae and Ithomiinae (Lepidoptera). Hereditas, 141(3), 216–236. https://doi.org/10.1111/j.1601-5223.2004.01868.x

Bushnell, B. (2019). BBMap. SourceForge. https://sourceforge.net/projects/bbmap/

Cantarel, B. L., Korf, I., Robb, S. M. C., Parra, G., Ross, E., Moore, B., Holt, C., Sánchez Alvarado, A., & Yandell, M. (2008). MAKER: An easy-to-use annotation pipeline designed for emerging model organism genomes. Genome Research, 18(1), 188–196. https://doi.org/10.1101/gr.6743907

Carbone, L., Alan Harris, R., Gnerre, S., Veeramah, K. R., Lorente-Galdos, B., Huddleston, J., Meyer, T. J., Herrero, J., Roos, C., Aken, B., Anaclerio, F., Archidiacono, N., Baker, C., Barrell, D., Batzer, M. A., Beal, K., Blancher, A., Bohrson, C. L., Brameier, M.,… Gibbs, R. A. (2014). Gibbon genome and the fast karyotype evolution of small apes. Nature, 513(7517), 195–201. https://doi.org/10.1038/nature13679

Catchen, J., Hohenlohe, P. A., Bassham, S., Amores, A., & Cresko, W. A. (2013). Stacks: An analysis tool set for population genomics. Molecular Ecology, 22(11), 3124–3140. https://doi.org/10.1111/mec.12354

Cicconardi, F., Lewis, J. J., Martin, S. H., Reed, R. D., Danko, C. G., & Montgomery, S. H. (2021). Chromosome Fusion Affects Genetic Diversity and Evolutionary Turnover of Functional Loci but Consistently Depends on Chromosome Size. Molecular Biology and Evolution, 38(10), 4449–4462. https://doi.org/10.1093/molbev/msab185

Davey, J. W., Chouteau, M., Barker, S. L., Maroja, L., Baxter, S. W., Simpson, F., Merrill, R. M., Joron, M., Mallet, J., Dasmahapatra, K. K., & Jiggins, C. D. (2016). Major Improvements to the Heliconius melpomene Genome Assembly Used to Confirm 10 Chromosome Fusion Events in 6 Million Years of Butterfly Evolution. G3: Genes, Genomes, Genetics, 6(3), 695–708. https://doi.org/10.1534/g3.115.023655

de Vos, J. M., Augustijnen, H., Bätscher, L., & Lucek, K. (2020). Speciation through chromosomal fusion and fission in Lepidoptera. Philosophical Transactions of the Royal Society B: Biological Sciences, 375(1806), 20190539. https://doi.org/10.1098/rstb.2019.0539

Dierckxsens, N., Mardulyn, P., & Smits, G. (2017). NOVOPlasty: De novo assembly of organelle genomes from whole genome data. Nucleic Acids Research, 45(4), e18. https://doi.org/10.1093/nar/gkw955

Dincā, V., Lukhtanov, V. A., Talavera, G., & Vila, R. (2011). Unexpected layers of cryptic diversity in wood white Leptidea butterflies. Nature Communications, 2, 324. https://doi.org/10.1038/ncomms1329

Dudchenko, O., Batra, S. S., Omer, A. D., Nyquist, S. K., Hoeger, M., Durand, N. C., Shamim, M. S., Machol, I., Lander, E. S., Aiden, A. P., & Aiden, E. L. (2017). De novo assembly of the Aedes aegypti genome using Hi-C yields chromosome-length scaffolds. Science, 356(6333), 92–95. https://doi.org/10.1126/science.aal3327

Durand, N. C., Robinson, J. T., Shamim, M. S., Machol, I., Mesirov, J. P., Lander, E. S., & Aiden, E. L. (2016). Juicebox Provides a Visualization System for Hi-C Contact Maps with Unlimited Zoom. Cell Systems, 3(1), 99–101. https://doi.org/10.1016/j.cels.2015.07.012

Durand, N. C., Shamim, M. S., Machol, I., Rao, S. S. P., Huntley, M. H., Lander, E. S., & Aiden, E. L. (2016). Juicer Provides a One-Click System for Analyzing Loop-Resolution Hi-C Experiments. Cell Systems, 3(1), 95–98. https://doi.org/10.1016/j.cels.2016.07.002

Espeland, M., Breinholt, J., Willmott, K. R., Warren, A. D., Vila, R., Toussaint, E. F. A., Maunsell, S. C., Aduse-Poku, K., Talavera, G., Eastwood, R., Jarzyna, M. A., Guralnick, R., Lohman, D. J., Pierce, N. E., & Kawahara, A. Y. (2018). A Comprehensive and Dated Phylogenomic Analysis of Butterflies. Current Biology: CB, 28(5), 770–778.e5. https://doi.org/10.1016/j.cub.2018.01.061

Faria, R., & Navarro, A. (2010). Chromosomal speciation revisited: Rearranging theory with pieces of evidence. Trends in Ecology & Evolution, 25(11), 660–669. https://doi.org/10.1016/j.tree.2010.07.008

Faulkner, J. S. (1972). Chromosome studies on Carex section Acutae in north-west Europe. Botanical Journal of the Linnean Society, 65(3), 271–301. https://doi.org/10.1111/j.1095-8339.1972.tb00120.x

Fraïsse, C., Picard, M. A. L., & Vicoso, B. (2017). The deep conservation of the Lepidoptera Z chromosome suggests a non-canonical origin of the W. Nature Communications, 8(1), 1486. https://doi.org/10.1038/s41467-017-01663-5

Hill, J., Rastas, P., Hornett, E. A., Neethiraj, R., Clark, N., Morehouse, N., Celorio-Mancera, M. de la P., Cols, J. C., Dircksen, H., Meslin, C., Keehnen, N., Pruisscher, P., Sikkink, K., Vives, M., Vogel, H., Wiklund, C., Woronik, A., Boggs, C. L., Nylin, S., & Wheat, C. W. (2019). Unprecedented reorganization of holocentric chromosomes provides insights into the enigma of lepidopteran chromosome evolution. Science Advances, 5(6), eaau3648. https://doi.org/10.1126/sciadv.aau3648

Höök, L., Leal, L., Talla, V., & Backström, N. (2019). Multilayered Tuning of Dosage Compensation and Z-Chromosome Masculinization in the Wood White (Leptidea sinapis) Butterfly. Genome Biology and Evolution, 11(9), 2633–2652. https://doi.org/10.1093/gbe/evz176

Iannucci, A., Altmanová, M., Ciofi, C., Ferguson-Smith, M., Milan, M., Pereira, J. C., Pether, J., Rehák, I., Rovatsos, M., Stanyon, R., Velenský, P., Ráb, P., Kratochvíl, L., & Johnson Pokorná, M. (2019). Conserved sex chromosomes and karyotype evolution in monitor lizards (Varanidae). Heredity, 123(2), 215–227. https://doi.org/10.1038/s41437-018-0179-6

Jones, P., Binns, D., Chang, H.-Y., Fraser, M., Li, W., McAnulla, C., McWilliam, H., Maslen, J., Mitchell, A., Nuka, G., Pesseat, S., Quinn, A. F., Sangrador-Vegas, A., Scheremetjew, M., Yong, S.-Y., Lopez, R., & Hunter, S. (2014). InterProScan 5: Genome-scale protein function classification. Bioinformatics, 30(9), 1236–1240. https://doi.org/10.1093/bioinformatics/btu031

Jurka, J. (1998). Repeats in genomic DNA: Mining and meaning. Current Opinion in Structural Biology, 8(3), 333–337. https://doi.org/10.1016/s0959-440x(98)80067-5

Kandul, N. P., Lukhtanov, V. A., & Pierce, N. E. (2007). Karyotypic diversity and speciation in Agrodiaetus butterflies. Evolution; International Journal of Organic Evolution, 61(3), 546–559. https://doi.org/10.1111/j.1558-5646.2007.00046.x

Kawahara, A. Y., & Breinholt, J. W. (2014). Phylogenomics provides strong evidence for relationships of butterflies and moths. Proceedings of the Royal Society B: Biological Sciences, 281(1788), 20140970. https://doi.org/10.1098/rspb.2014.0970

Kawakami, T., Butlin, R. K., Adams, M., Paull, David. J., & Cooper, S. J. B. (2009). Genetic Analysis of a Chromosomal Hybrid Zone in the Australian Morabine Grasshoppers (vandiemenella, Viatica Species Group). Evolution, 63(1), 139–152. https://doi.org/10.1111/j.1558-5646.2008.00526.x

Kohany, O., Gentles, A. J., Hankus, L., & Jurka, J. (2006). Annotation, submission and screening of repetitive elements in Repbase: RepbaseSubmitter and Censor. BMC Bioinformatics, 7(1), 474. https://doi.org/10.1186/1471-2105-7-474

Korf, I. (2004). Gene finding in novel genomes. BMC Bioinformatics, 5(1), 59. https://doi.org/10.1186/1471-2105-5-59

Krueger, F. (2019). Babraham Bioinformatics—Trim Galore! https://www.bioinformatics.babraham.ac.uk/projects/trim_galore/

Krzywinski, M., Schein, J., Birol, I., Connors, J., Gascoyne, R., Horsman, D., Jones, S. J., & Marra, M. A. (2009). Circos: An information aesthetic for comparative genomics. Genome Research, 19(9), 1639–1645. https://doi.org/10.1101/gr.092759.109

Laetsch, D. R., & Blaxter, M. L. (2017). BlobTools: Interrogation of genome assemblies (6:1287). F1000Research. https://doi.org/10.12688/f1000research.12232.1

Larson, A., Prager, E. M., & Wilson, A. C. (1984). Chromosomal evolution, speciation and morphological change in vertebrates: The role of social behaviour. In M. D. Bennett, A. Gropp, & U. Wolf (Eds.), Chromosomes Today: Volume 8 Proceedings of the Eighth International Chromosome Conference held in Lübeck, West Germnay, 21–24 September 1983 (pp. 215–228). Springer Netherlands. https://doi.org/10.1007/978-94-010-9163-3_20

Lewis, J. J., Cicconardi, F., Martin, S. H., Reed, R. D., Danko, C. G., & Montgomery, S. H. (2021). The Dryas iulia Genome Supports Multiple Gains of a W Chromosome from a B Chromosome in Butterflies. Genome Biology and Evolution, 13(7), evab128. https://doi.org/10.1093/gbe/evab128

Li, H. (2011). Improving SNP discovery by base alignment quality. Bioinformatics, 27(8), 1157–1158. https://doi.org/10.1093/bioinformatics/btr076

Li, H. (2013). Aligning sequence reads, clone sequences and assembly contigs with BWA-MEM (arXiv:1303.3997). arXiv. https://doi.org/10.48550/arXiv.1303.3997

Li, H., Handsaker, B., Wysoker, A., Fennell, T., Ruan, J., Homer, N., Marth, G., Abecasis, G., Durbin, R., & 1000 Genome Project Data Processing Subgroup. (2009). The Sequence Alignment/Map format and SAMtools. Bioinformatics (Oxford, England), 25(16), 2078–2079. https://doi.org/10.1093/bioinformatics/btp352

Lohse, K., Höök, L., Näsvall, K., & Backström, N. (2022). The genome sequence of the wood white butterfly, Leptidea sinapis (Linnaeus, 1758). Welcome Open Research, Submitted.

Lukhtanov, V. A., Dincā, V., Friberg, M., Šíchová, J., Olofsson, M., Vila, R., Marec, F., & Wiklund, C. (2018). Versatility of multivalent orientation, inverted meiosis, and rescued fitness in holocentric chromosomal hybrids. Proceedings of the National Academy of Sciences, 115(41), E9610–E9619. https://doi.org/10.1073/pnas.1802610115

Lukhtanov, V. A., Dincā, V., Friberg, M., Vila, R., & Wiklund, C. (2020). Incomplete Sterility of Chromosomal Hybrids: Implications for Karyotype Evolution and Homoploid Hybrid Speciation. Frontiers in Genetics, 11. https://doi.org/10.3389/fgene.2020.583827

Lukhtanov, V. A., Dincā, V., Talavera, G., & Vila, R. (2011). Unprecedented within-species chromosome number cline in the Wood White butterfly Leptidea sinapis and its significance for karyotype evolution and speciation. BMC Evolutionary Biology, 11, 109. https://doi.org/10.1186/1471-2148-11-109

Marçais, G., Delcher, A. L., Phillippy, A. M., Coston, R., Salzberg, S. L., & Zimin, A. (2018). MUMmer4: A fast and versatile genome alignment system. PLoS Computational Biology, 14(1), e1005944. https://doi.org/10.1371/journal.pcbi.1005944

Marks, P., Garcia, S., Barrio, A. M., Belhocine, K., Bernate, J., Bharadwaj, R., Bjornson, K., Catalanotti, C., Delaney, J., Fehr, A., Fiddes, I. T., Galvin, B., Heaton, H., Herschleb, J., Hindson, C., Holt, E., Jabara, C. B., Jett, S., Keivanfar, N.,… Church, D. M. (2019). Resolving the full spectrum of human genome variation using Linked-Reads. Genome Research, 29(4), 635–645. https://doi.org/10.1101/gr.234443.118

Martin, M. (2011). Cutadapt removes adapter sequences from high-throughput sequencing reads. EMBnet.Journal, 17(1), 10–12. https://doi.org/10.14806/ej.17.1.200

Mathers, T. C., Wouters, R. H. M., Mugford, S. T., Swarbreck, D., van Oosterhout, C., & Hogenhout, S. A. (2021). Chromosome-Scale Genome Assemblies of Aphids Reveal Extensively Rearranged Autosomes and Long-Term Conservation of the X Chromosome. Molecular Biology and Evolution, 38(3), 856–875. https://doi.org/10.1093/molbev/msaa246

Mayrose, I., & Lysak, M. A. (2021). The Evolution of Chromosome Numbers: Mechanistic Models and Experimental Approaches. Genome Biology and Evolution, 13(2), evaa220. https://doi.org/10.1093/gbe/evaa220

McKenna, A., Hanna, M., Banks, E., Sivachenko, A., Cibulskis, K., Kernytsky, A., Garimella, K., Altshuler, D., Gabriel, S., Daly, M., & DePristo, M. A. (2010). The Genome Analysis Toolkit: A MapReduce framework for analyzing next-generation DNA sequencing data. Genome Research, 20(9), 1297–1303. https://doi.org/10.1101/gr.107524.110

Melters, D. P., Paliulis, L. V., Korf, I. F., & Chan, S. W. L. (2012). Holocentric chromosomes: Convergent evolution, meiotic adaptations, and genomic analysis. Chromosome Research, 20(5), 579–593. https://doi.org/10.1007/s10577-012-9292-1

Miller, W. J., & Capy, P. (2004). Mobile genetic elements as natural tools for genome evolution. Methods in Molecular Biology (Clifton, N.J.), 260, 1–20. https://doi.org/10.1385/1-59259-755-6:001

Mongue, A. J., Nguyen, P., Voleníková, A., & Walters, J. R. (2017). Neo-sex Chromosomes in the Monarch Butterfly, Danaus plexippus. G3 Genes|Genomes|Genetics, 7(10), 3281–3294. https://doi.org/10.1534/g3.117.300187

Mongue, A. J., & Walters, J. R. (2018). The Z chromosome is enriched for sperm proteins in two divergent species of Lepidoptera. Genome, 61(4), 248–253. https://doi.org/10.1139/gen-2017-0068

Nanda, I., Schlegelmilch, K., Haaf, T., Schartl, M., & Schmid, M. (2008). Synteny conservation of the Z chromosome in 14 avian species (11 families) supports a role for Z dosage in avian sex determination. Cytogenetic and Genome Research, 122(2), 150–156. https://doi.org/10.1159/000163092

Nguyen, P., Sýkorová, M., Šíchová, J., Kůta, V., Dalíková, M., Čapková Frydrychová, R., Neven, L. G., Sahara, K., & Marec, F. (2013). Neo-sex chromosomes and adaptive potential in tortricid pests. Proceedings of the National Academy of Sciences, 110(17), 6931–6936. https://doi.org/10.1073/pnas.1220372110

Okazaki, S., Ishikawa, H., & Fujiwara, H. (1995). Structural analysis of TRAS1, a novel family of telomeric repeat-associated retrotransposons in the silkworm, Bombyx mori. Molecular and Cellular Biology, 15(8), 4545–4552. https://doi.org/10.1128/MCB.15.8.4545

Okonechnikov, K., Conesa, A., & García-Alcalde, F. (2016). Qualimap 2: Advanced multi-sample quality control for high-throughput sequencing data. Bioinformatics, 32(2), 292–294. https://doi.org/10.1093/bioinformatics/btv566

Pazhenkova, E. A., & Lukhtanov, V. A. (2022). Chromosomal conservatism vs chromosomal megaevolution: Enigma of karyotypic evolution in Lepidoptera (p. 2022.06.05.494852). bioRxiv. https://doi.org/10.1101/2022.06.05.494852

Pennell, M. W., Kirkpatrick, M., Otto, S. P., Vamosi, J. C., Peichel, C. L., Valenzuela, N., & Kitano, J. (2015). Y Fuse? Sex Chromosome Fusions in Fishes and Reptiles. PLOS Genetics, 11(5), e1005237. https://doi.org/10.1371/journal.pgen.1005237

Petitpierre, E. (1987). Why beetles have strikingly different rates of chromosomal evolution? Elytron. Bulletin of the European Association of Coleopterology, 1, 25–32.

Poorten, T. (2018). DotPlotly [HTML]. https://github.com/tpoorten/dotPlotly (Original work published 2017)

Pringle, E. G., Baxter, S. W., Webster, C. L., Papanicolaou, A., Lee, S. F., & Jiggins, C. D. (2007). Synteny and Chromosome Evolution in the Lepidoptera: Evidence From Mapping in Heliconius melpomene. Genetics, 177(1), 417–426. https://doi.org/10.1534/genetics.107.073122

Quinlan, A. R., & Hall, I. M. (2010). BEDTools: A flexible suite of utilities for comparing genomic features. Bioinformatics (Oxford, England), 26(6), 841–842. https://doi.org/10.1093/bioinformatics/btq033

R Core Team. (2019). R: The R Project for Statistical Computing. https://www.r-project.org/

Rastas, P. (2017). Lep-MAP3: Robust linkage mapping even for low-coverage whole genome sequencing data. Bioinformatics, 33(23), 3726–3732. https://doi.org/10.1093/bioinformatics/btx494

Rieseberg, L. H. (2001). Chromosomal rearrangements and speciation. Trends in Ecology & Evolution, 16(7), 351–358. https://doi.org/10.1016/s0169-5347(01)02187-5

Robinson, R. (1971). Lepidoptera Genetics. Pergamon Press.

Román-Palacios, C., Medina, C. A., Zhan, S. H., & Barker, M. S. (2021). Animal chromosome counts reveal a similar range of chromosome numbers but with less polyploidy in animals compared to flowering plants. Journal of Evolutionary Biology, 34(8), 1339–1339. https://doi.org/10.1111/jeb.13884

Rovatsos, M., Vukiċ, J., Lymberakis, P., & Kratochvíl, L. (2015). Evolutionary stability of sex chromosomes in snakes. Proceedings. Biological Sciences, 282(1821), 20151992. https://doi.org/10.1098/rspb.2015.1992

Ruckman, S. N., Jonika, M. M., Casola, C., & Blackmon, H. (2020). Chromosome number evolves at equal rates in holocentric and monocentric clades. PLOS Genetics, 16(10), e1009076. https://doi.org/10.1371/journal.pgen.1009076

Sahara, K., Yoshido, A., & Traut, W. (2012). Sex chromosome evolution in moths and butterflies. Chromosome Research, 20(1), 83–94. https://doi.org/10.1007/s10577-011-9262-z

Sambrook, J., & Russell, D. W. (2006). Purification of nucleic acids by extraction with phenol:chloroform. CSH Protocols, 2006(1), pdb.prot4455. https://doi.org/10.1101/pdb.prot4455

Šíchová, J., Ohno, M., Dincā, V., Watanabe, M., Sahara, K., & Marec, F. (2016). Fissions, fusions, and translocations shaped the karyotype and multiple sex chromosome constitution of the northeast-Asian wood white butterfly, Leptidea amurensis. Biological Journal of the Linnean Society, 118(3), 457–471. https://doi.org/10.1111/bij.12756

Šíchová, J., Voleníková, A., Dincā, V., Nguyen, P., Vila, R., Sahara, K., & Marec, F. (2015). Dynamic karyotype evolution and unique sex determination systems in Leptidea wood white butterflies. BMC Evolutionary Biology, 15, 89. https://doi.org/10.1186/s12862-015-0375-4

Simão, F. A., Waterhouse, R. M., Ioannidis, P., Kriventseva, E. V., & Zdobnov, E. M. (2015). BUSCO: Assessing genome assembly and annotation completeness with single-copy orthologs. Bioinformatics, 31(19), 3210–3212. https://doi.org/10.1093/bioinformatics/btv351

Smit, A., & Hubley, R. (2017). RepeatModeler. https://www.repeatmasker.org/RepeatModeler/

Smit, A., Hubley, R., & Green, P. (2019). RepeatMasker. https://www.repeatmasker.org/

Sotero-Caio, C. G., Volleth, M., Hoffmann, F. G., Scott, L., Wichman, H. A., Yang, F., & Baker, R. J. (2015). Integration of molecular cytogenetics, dated molecular phylogeny, and model-based predictions to understand the extreme chromosome reorganization in the Neotropical genus Tonatia (Chiroptera: Phyllostomidae). BMC Evolutionary Biology, 15(1), 220. https://doi.org/10.1186/s12862-015-0494-y

Stanke, M., Diekhans, M., Baertsch, R., & Haussler, D. (2008). Using native and syntenically mapped cDNA alignments to improve de novo gene finding. Bioinformatics, 24(5), 637–644. https://doi.org/10.1093/bioinformatics/btn013

Suomalainen, E. (1953). The Kinetochore and the Bivalent Structure in the Lepidoptera. Hereditas, 39(1–2), 88–96. https://doi.org/10.1111/j.1601-5223.1953.tb03403.x

Sylvester, T., Hjelmen, C. E., Hanrahan, S. J., Lenhart, P. A., Johnston, J. S., & Blackmon, H. (2020). Lineage-specific patterns of chromosome evolution are the rule not the exception in Polyneoptera insects. Proceedings of the Royal Society B: Biological Sciences, 287(1935), 20201388. https://doi.org/10.1098/rspb.2020.1388

Takahashi, H., Okazaki, S., & Fujiwara, H. (1997). A New Family of Site-Specific Retrotransposons, SART1, Is Inserted into Telomeric Repeats of the Silkworm, Bombyx Mori. Nucleic Acids Research, 25(8), 1578–1584. https://doi.org/10.1093/nar/25.8.1578

Talla, V., Soler, L., Kawakami, T., Dincā, V., Vila, R., Friberg, M., Wiklund, C., & Backström, N. (2019). Dissecting the Effects of Selection and Mutation on Genetic Diversity in Three Wood White (Leptidea) Butterfly Species. Genome Biology and Evolution, 11(10), 2875–2886. https://doi.org/10.1093/gbe/evz212

Talla, V., Suh, A., Kalsoom, F., Dinca, V., Vila, R., Friberg, M., Wiklund, C., & Backström, N. (2017). Rapid Increase in Genome Size as a Consequence of Transposable Element Hyperactivity in Wood-White (Leptidea) Butterflies. Genome Biology and Evolution, 9(10), 2491–2505. https://doi.org/10.1093/gbe/evx163

Tang, M., He, S., Gong, X., Lü, P., Taha, R. H., & Chen, K. (2021). High-Quality de novo Chromosome-Level Genome Assembly of a Single Bombyx mori With BmNPV Resistance by a Combination of PacBio Long-Read Sequencing, Illumina Short-Read Sequencing, and Hi-C Sequencing. Frontiers in Genetics, 12, 718266. https://doi.org/10.3389/fgene.2021.718266

Traut, W., Sahara, K., & Marec, F. (2007). Sex Chromosomes and Sex Determination in Lepidoptera. Sexual Development, 1(6), 332–346. https://doi.org/10.1159/000111765

Turner, J. R. G., & Sheppard, P. M. (1975). Absence of crossing-over in female butterflies (Heliconius). Heredity, 34(2), 265–269. https://doi.org/10.1038/hdy.1975.29

Vicoso, B. (2019). Molecular and evolutionary dynamics of animal sex-chromosome turnover. Nature Ecology & Evolution, 3(12), 1632–1641. https://doi.org/10.1038/s41559-019-1050-8

Wang, Y., Tang, H., Debarry, J. D., Tan, X., Li, J., Wang, X., Lee, T., Jin, H., Marler, B., Guo, H., Kissinger, J. C., & Paterson, A. H. (2012). MCScanX: A toolkit for detection and evolutionary analysis of gene synteny and collinearity. Nucleic Acids Research, 40(7), e49. https://doi.org/10.1093/nar/gkr1293

Weisenfeld, N. I., Kumar, V., Shah, P., Church, D. M., & Jaffe, D. B. (2017). Direct determination of diploid genome sequences. Genome Research, 27(5), 757–767. https://doi.org/10.1101/gr.214874.116

Wingett, S. W., & Andrews, S. (2018). FastQ Screen: A tool for multi-genome mapping and quality control. F1000Research, 7, 1338. https://doi.org/10.12688/f1000research.15931.2

Yoshido, A., Šíchová, J., Pospíšilová, K., Nguyen, P., Voleníková, A., Šafář, J., Provazník, J., Vila, R., & Marec, F. (2020). Evolution of multiple sex-chromosomes associated with dynamic genome reshuffling in Leptidea wood-white butterflies. Heredity, 125(3), 138–154. https://doi.org/10.1038/s41437-020-0325-9

