## Supplemental_material for "High-density linkage maps and chromosome level genome assemblies unveil direction and frequency of extensive structural rearrangements in wood white butterflies (*Leptidea* spp.)"

### Supplementary material

#### Supplementary figures

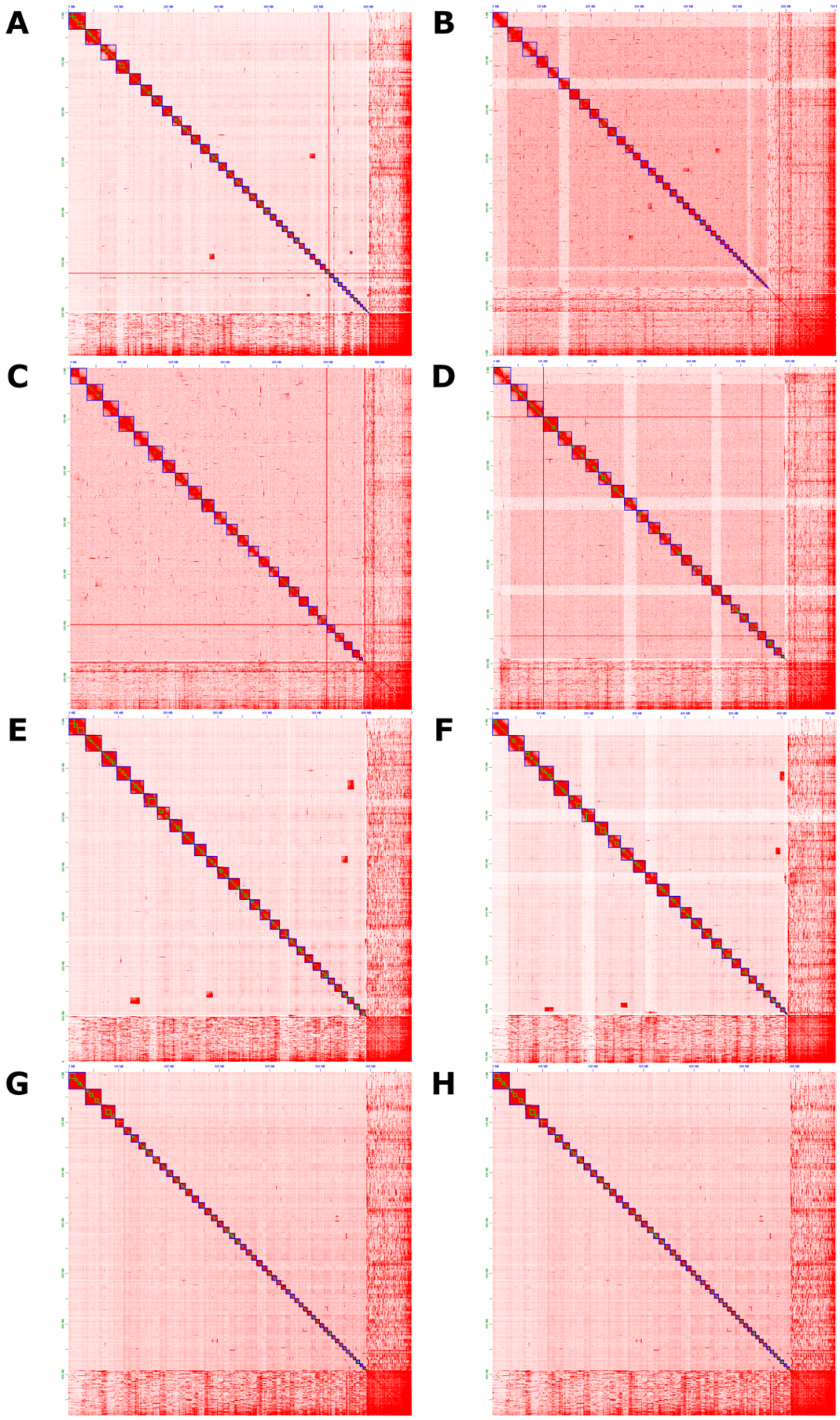

*Supplementary figure 1. Heatmaps of HiC interactions across the genomes. A: L. juvernica male, B: L. juvernica female, C: L. reali male, D: L. reali female, E: L. sinapis Swe male, F: L. sinapis Swe female, G: L. sinapis Cat male, H: L. sinapis female.*

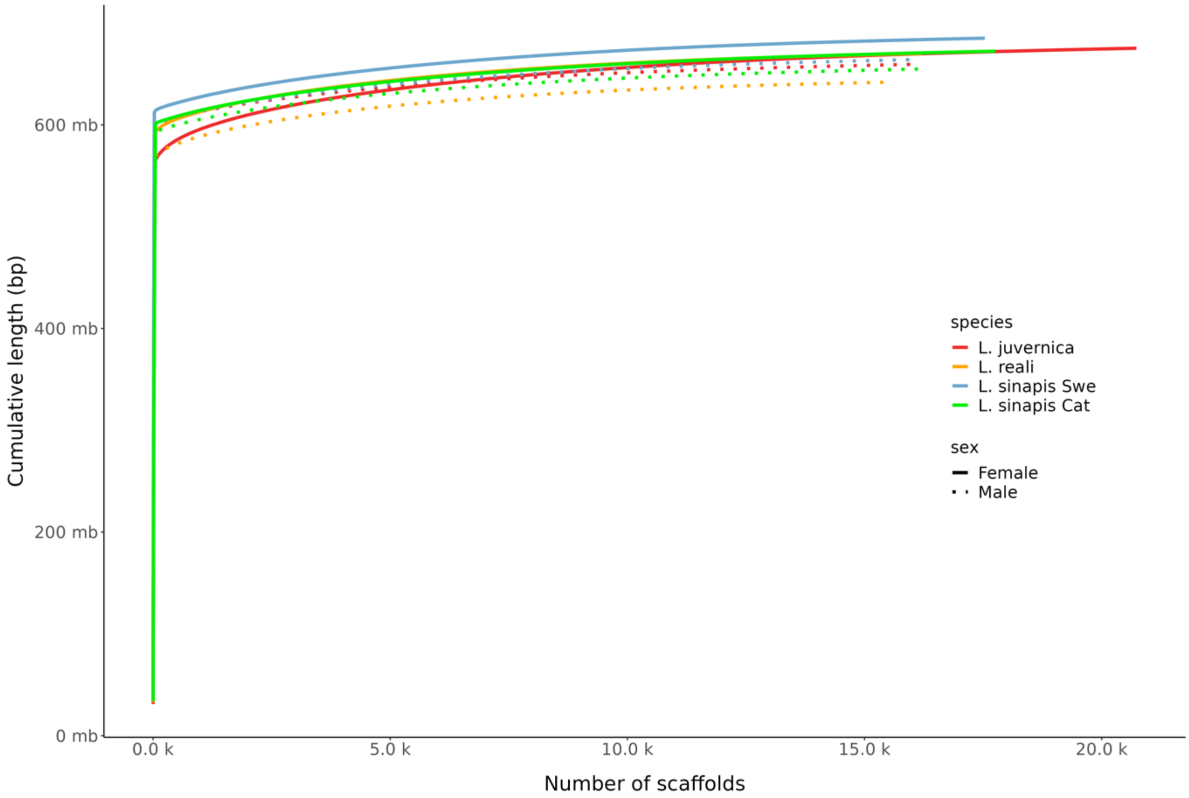

*Supplementary figure 2. Cumulative scaffold length. Scaffolds are ordered by size.*

A

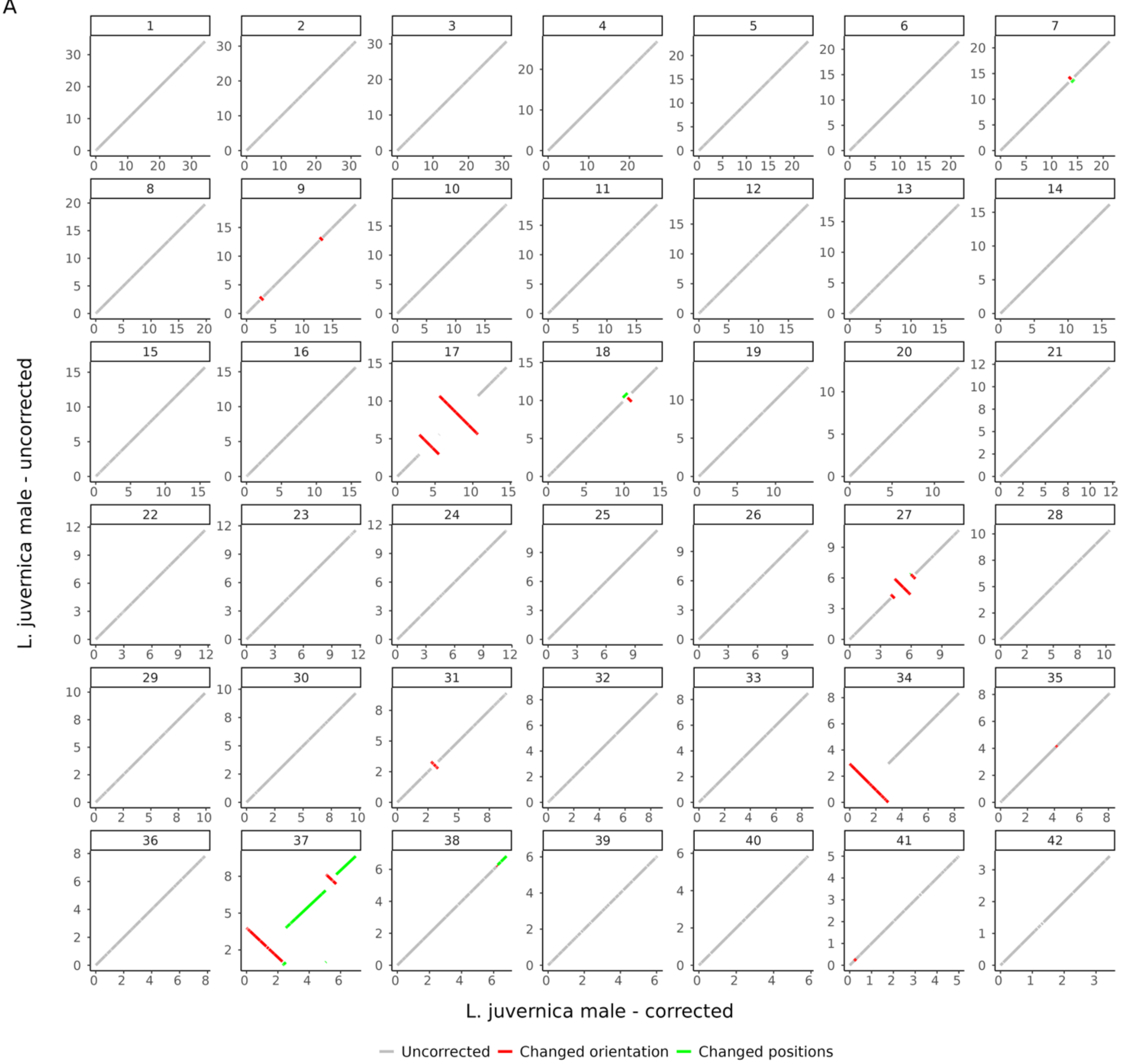

L. juvernica male - uncorrected
L. juvernica male - corrected
Uncorrected
Changed orientation
Changed positions

B

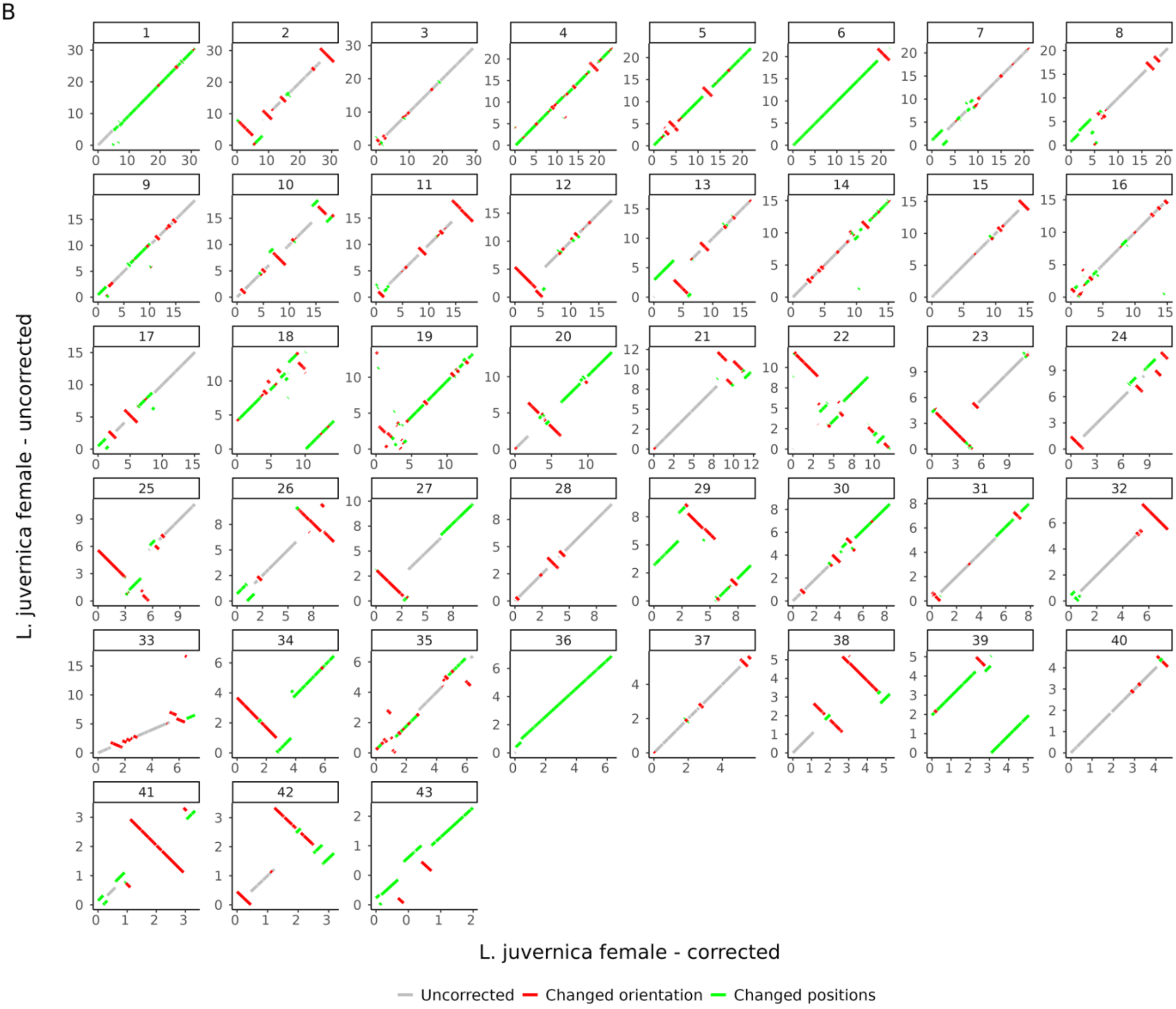
L. juvernica female - uncorrected
L. juvernica female - corrected
Uncorrected
Changed orientation
Changed positions

C

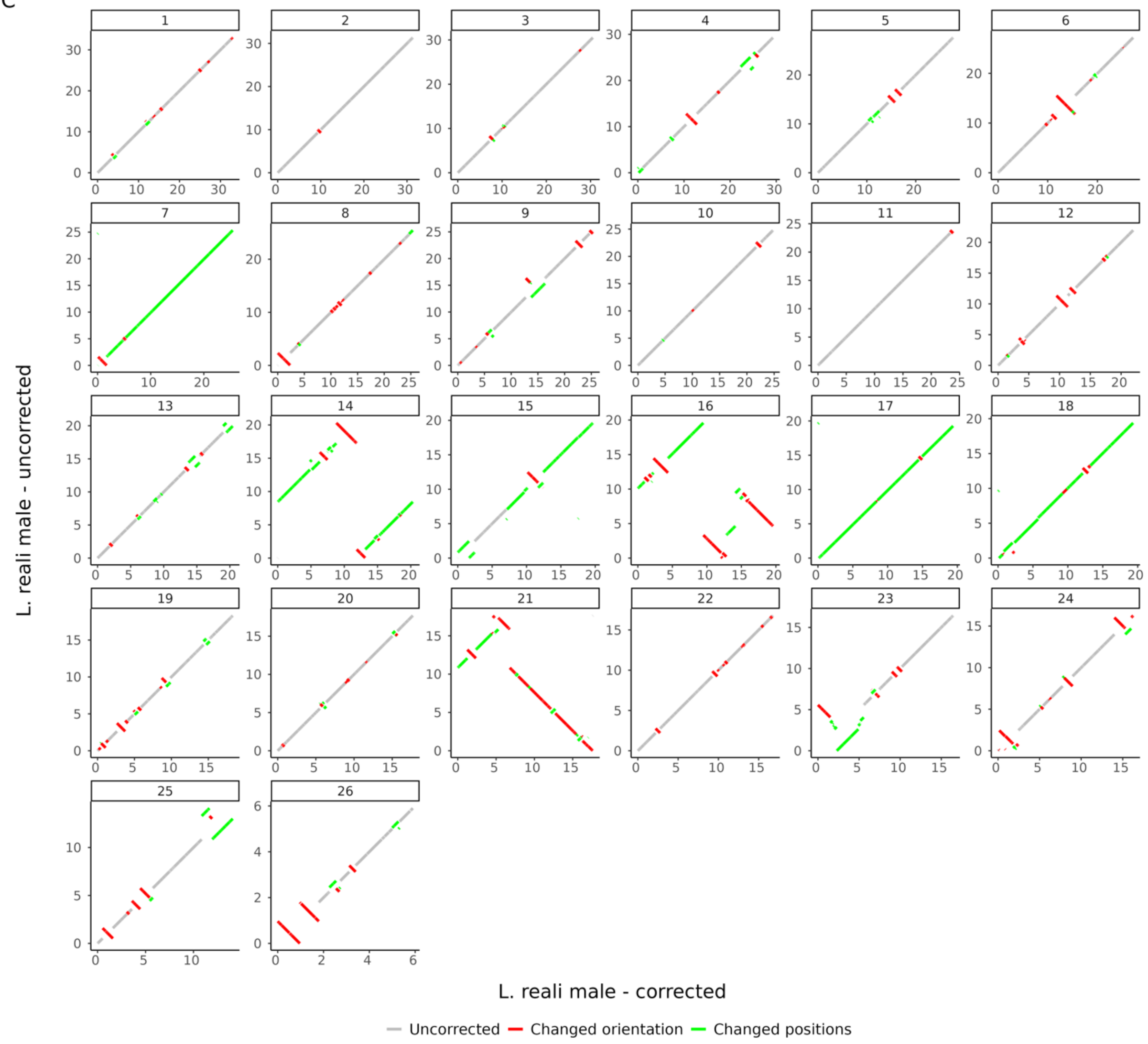
L. reali male - uncorrected
L. reali male - corrected
Uncorrected
Changed orientation
Changed positions

D

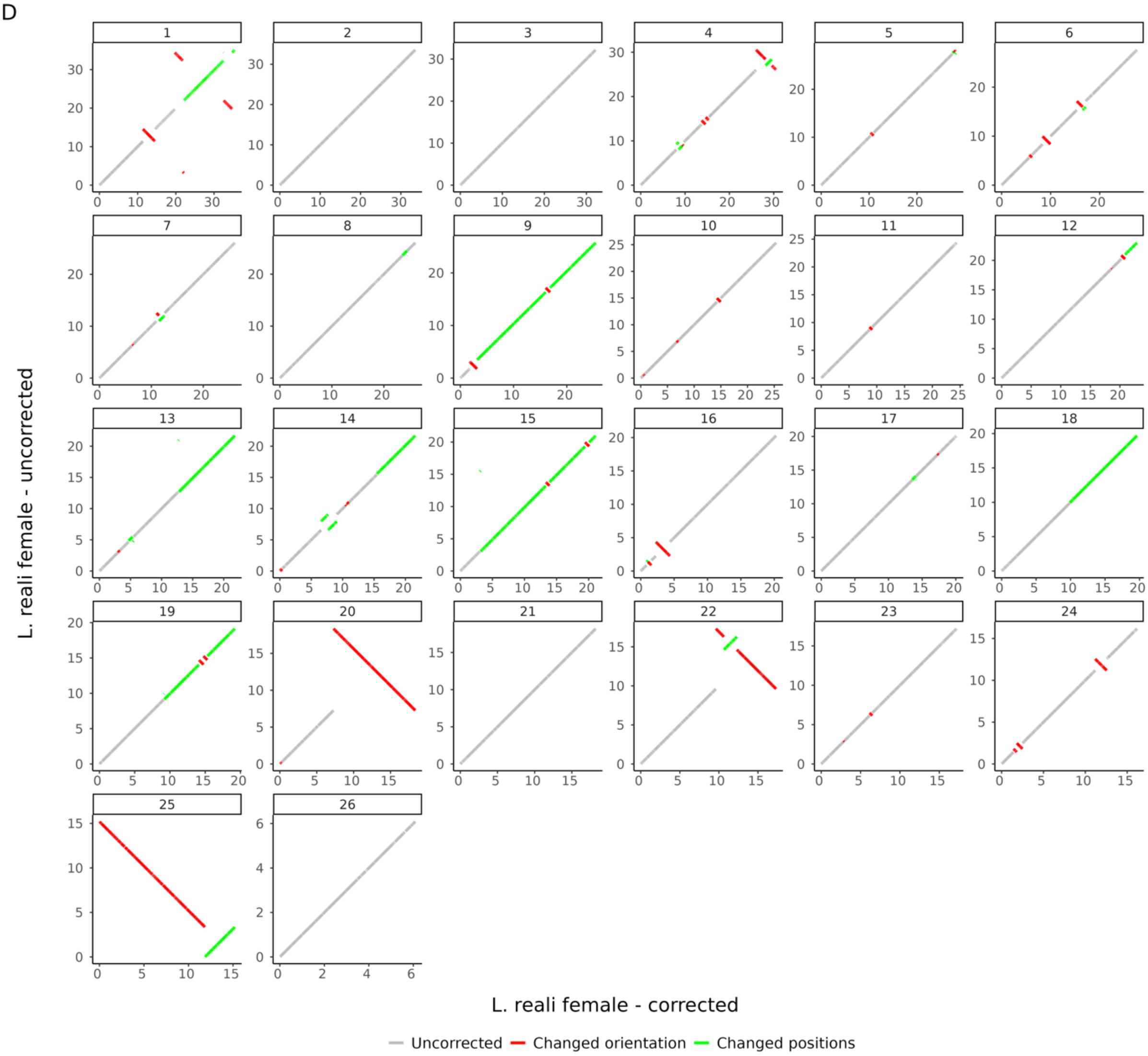

L. reali female - uncorrected
L. reali female - corrected
Uncorrected
Changed orientation
Changed positions

E

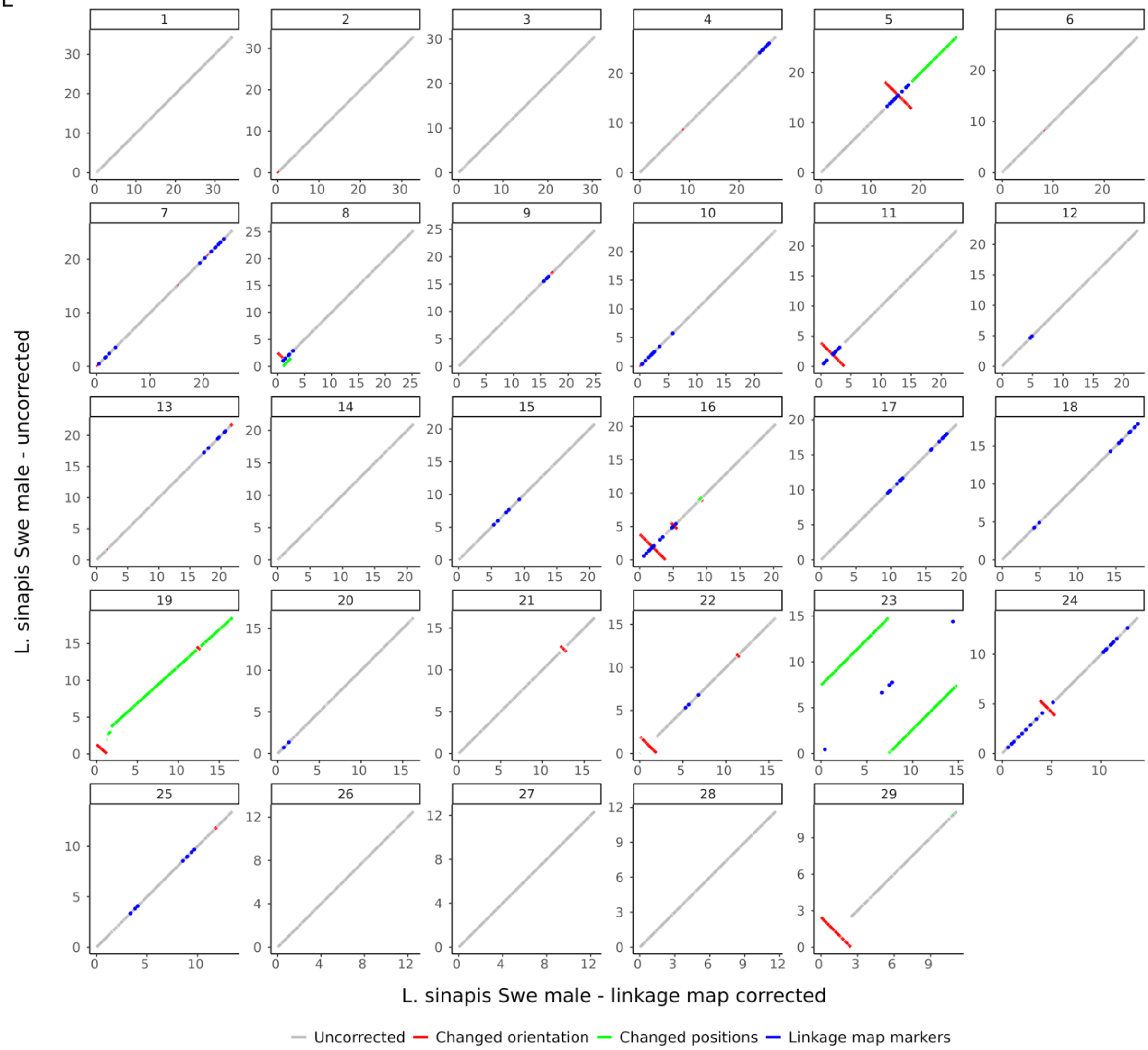

L. sinapis Swe male - uncorrected
L. sinapis Swe male - linkage map corrected
Uncorrected
Changed orientation
Changed positions
Linkage map markers

F

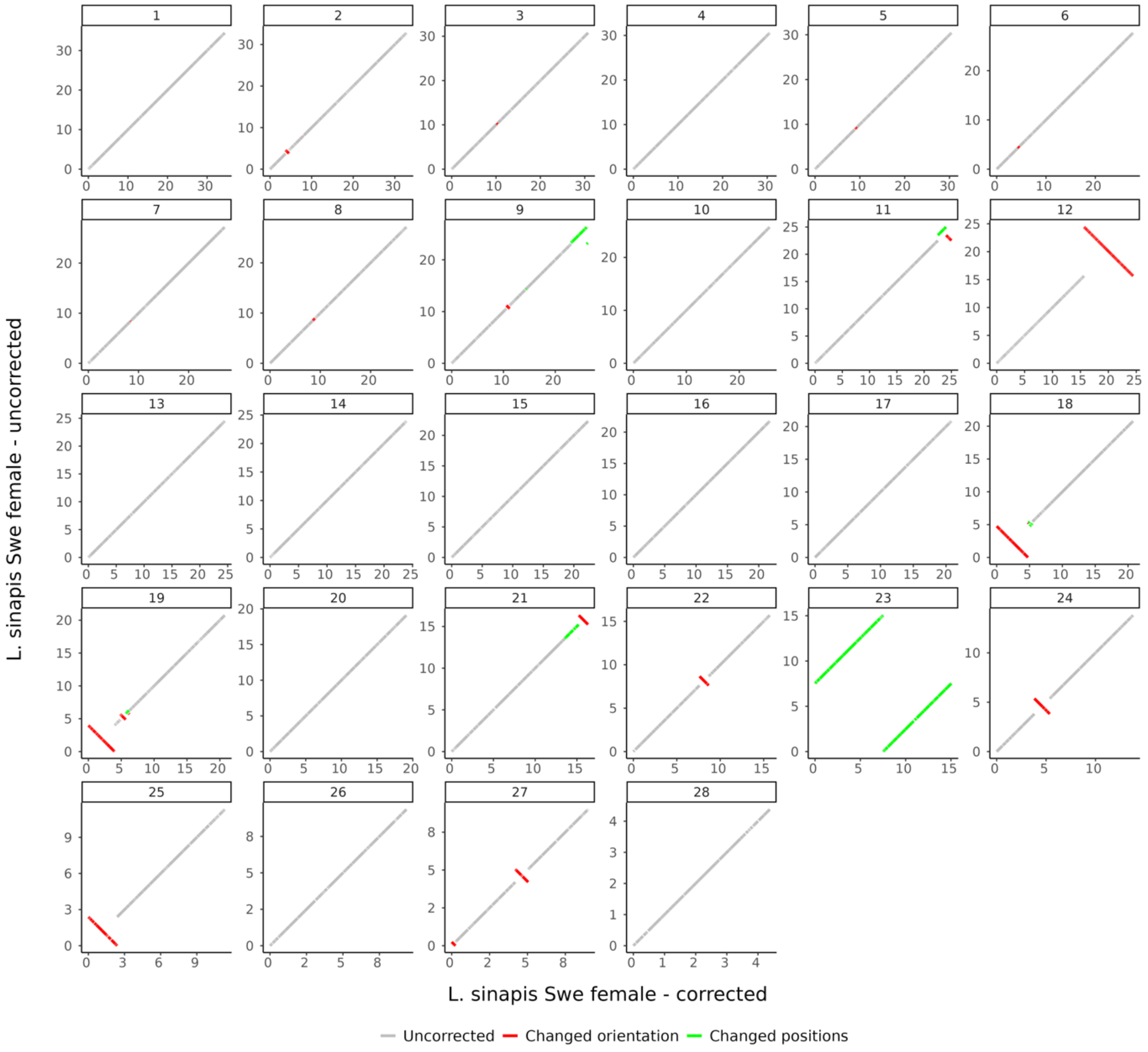

L. sinapis Swe female - uncorrected
L. sinapis Swe female - corrected
Uncorrected
Changed orientation
Changed positions

G

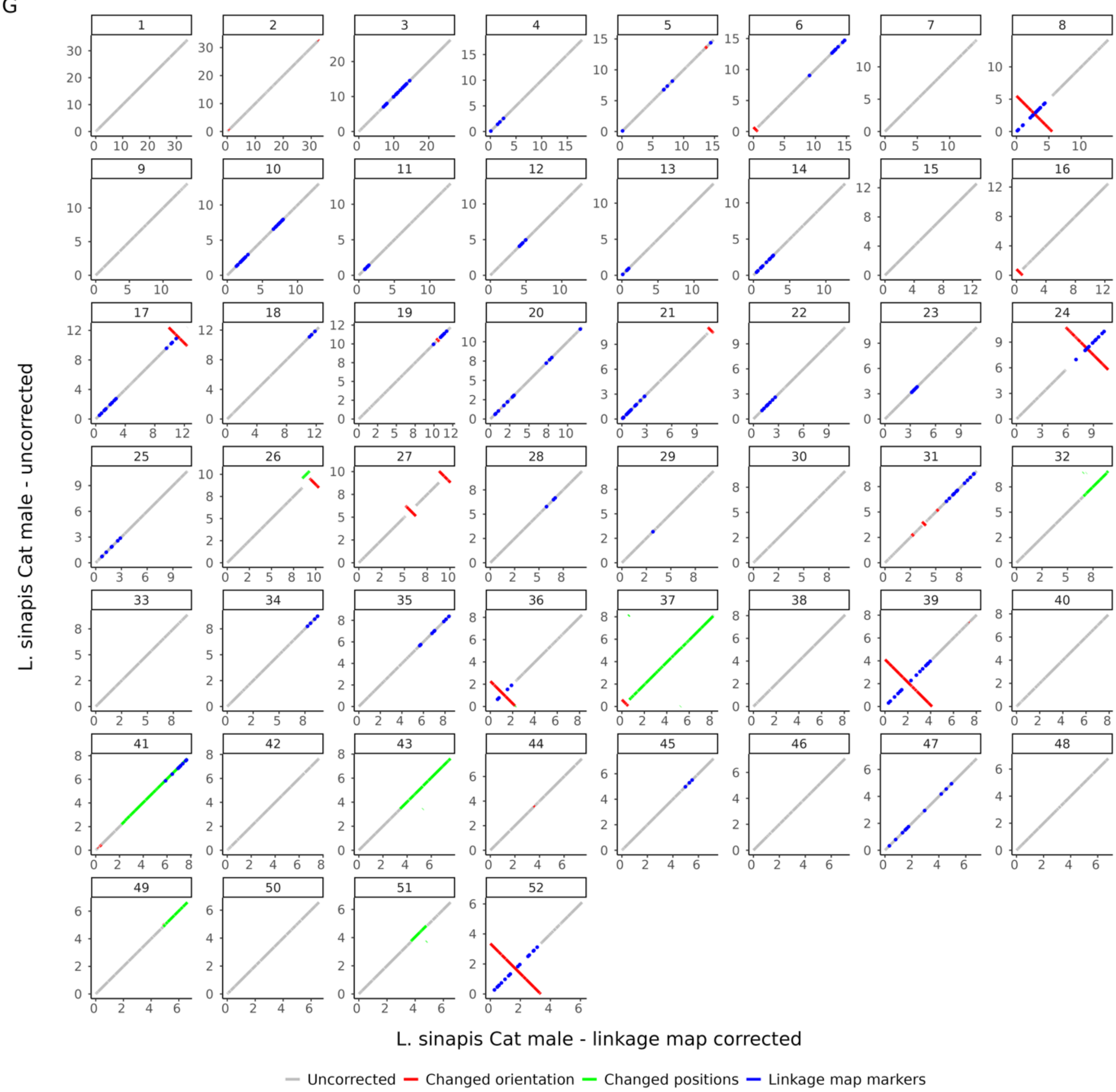

L. sinapis Cat male - uncorrected
L. sinapis Cat male - linkage map corrected
Uncorrected
Changed orientation
Changed positions
Linkage map markers

H

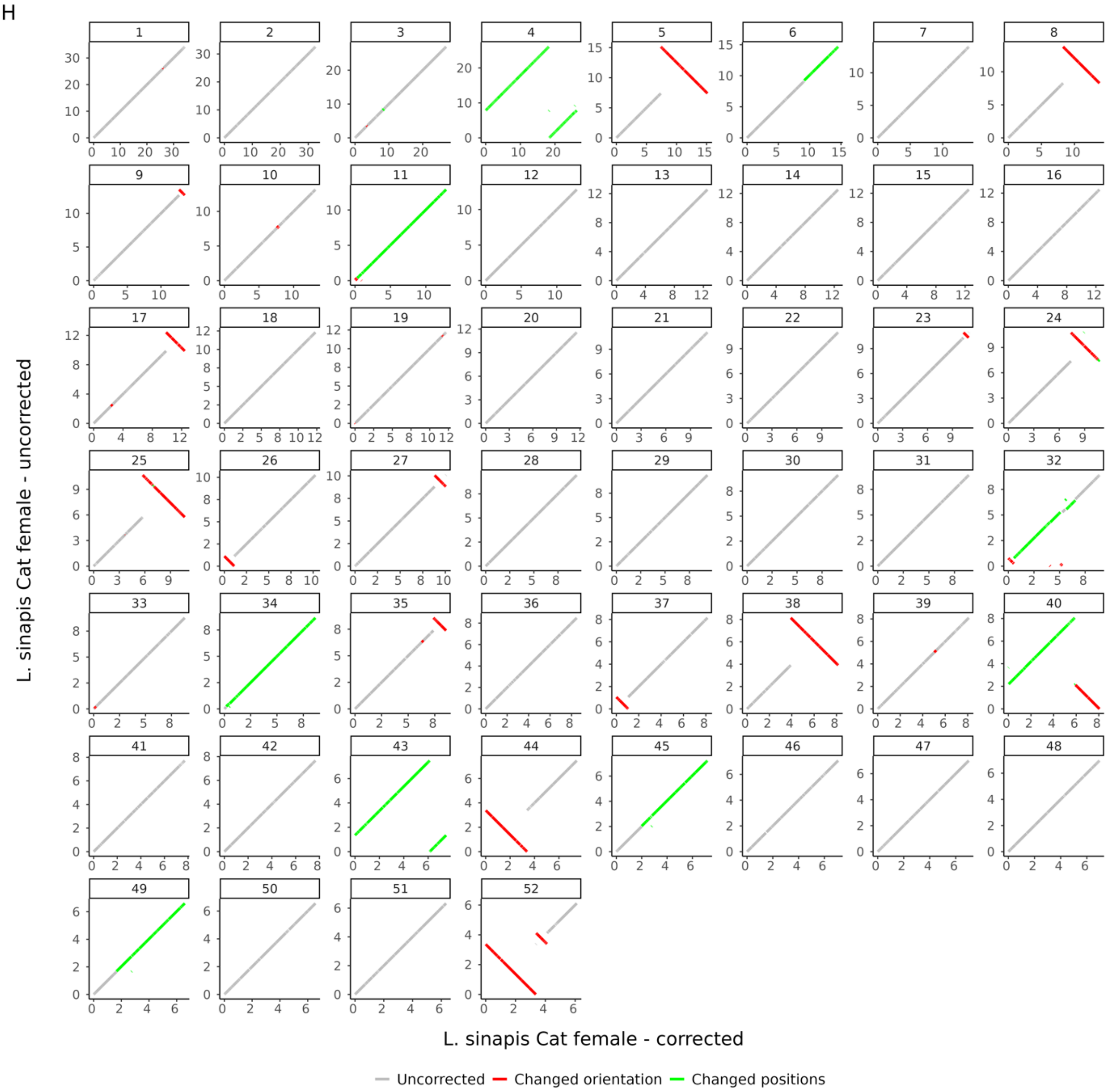

*Supplementary figure 3. Sequence alignments comparing before (query) and after (reference) assembly corrections guided by linkage map information (E and M), HiC data and pairwise alignments (A-H) between species and populations. Linkage map markers (highlighted in blue) show markers deviating from the physical assembly positions. Scale is in mb. A: L. juvernica male, B: L. juvernica female, C: L. reali male, D: L. reali female E: L. sinapis Swe male, F: L. sinapis Cat male, G: L. sinapis Swe female, H: L. sinapis Cat female.*

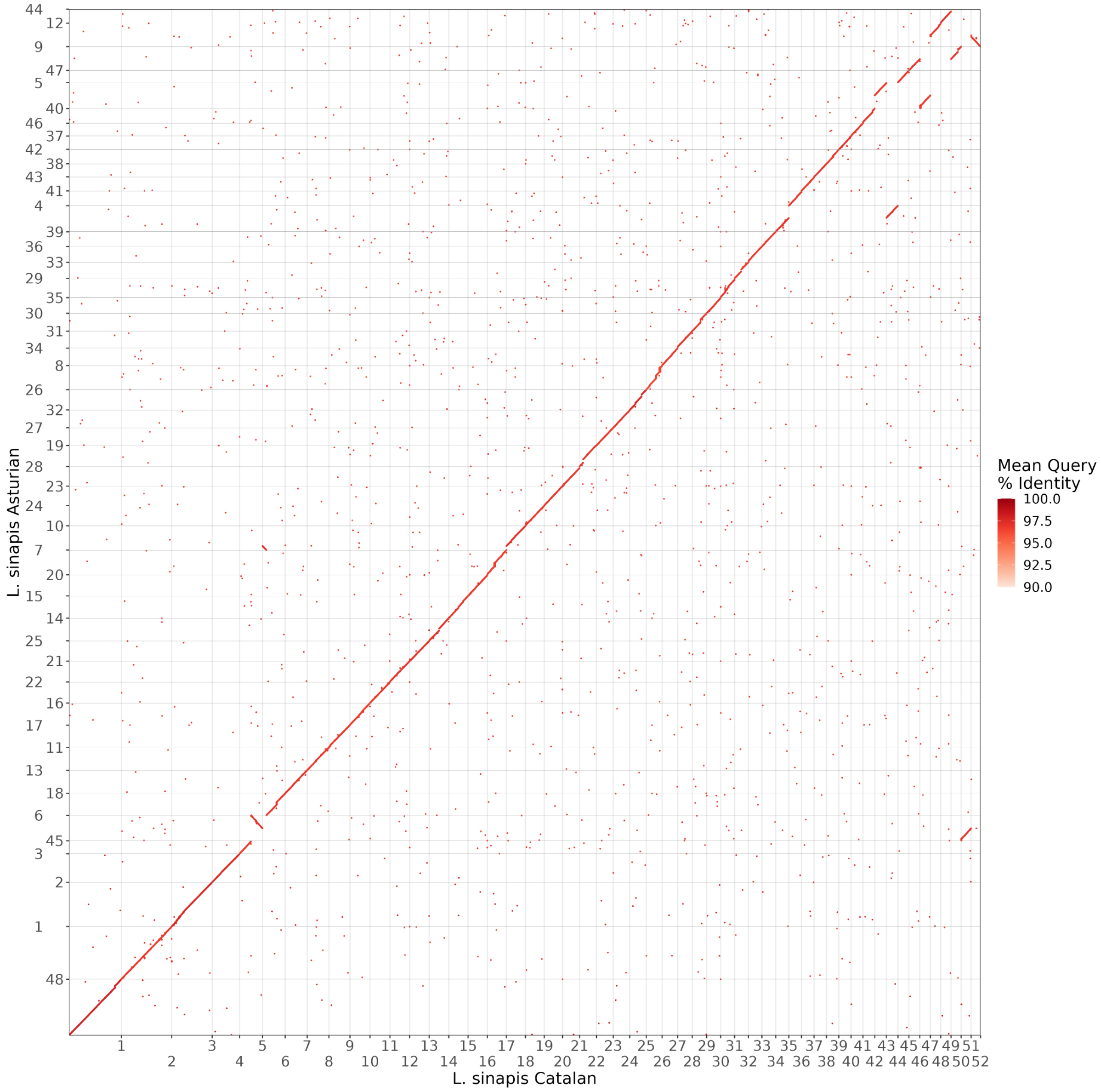

*Supplementary figure 4. Dotplot of alignments (>1kb) between L. sinapis Catalan male and L. sinapis Asturian (DToL) assemblies. Scaffolds in the query are oriented so the majority of alignments are in the same direction as the reference. Mean query % identity shows mean identity for all alignments in each scaffold.*

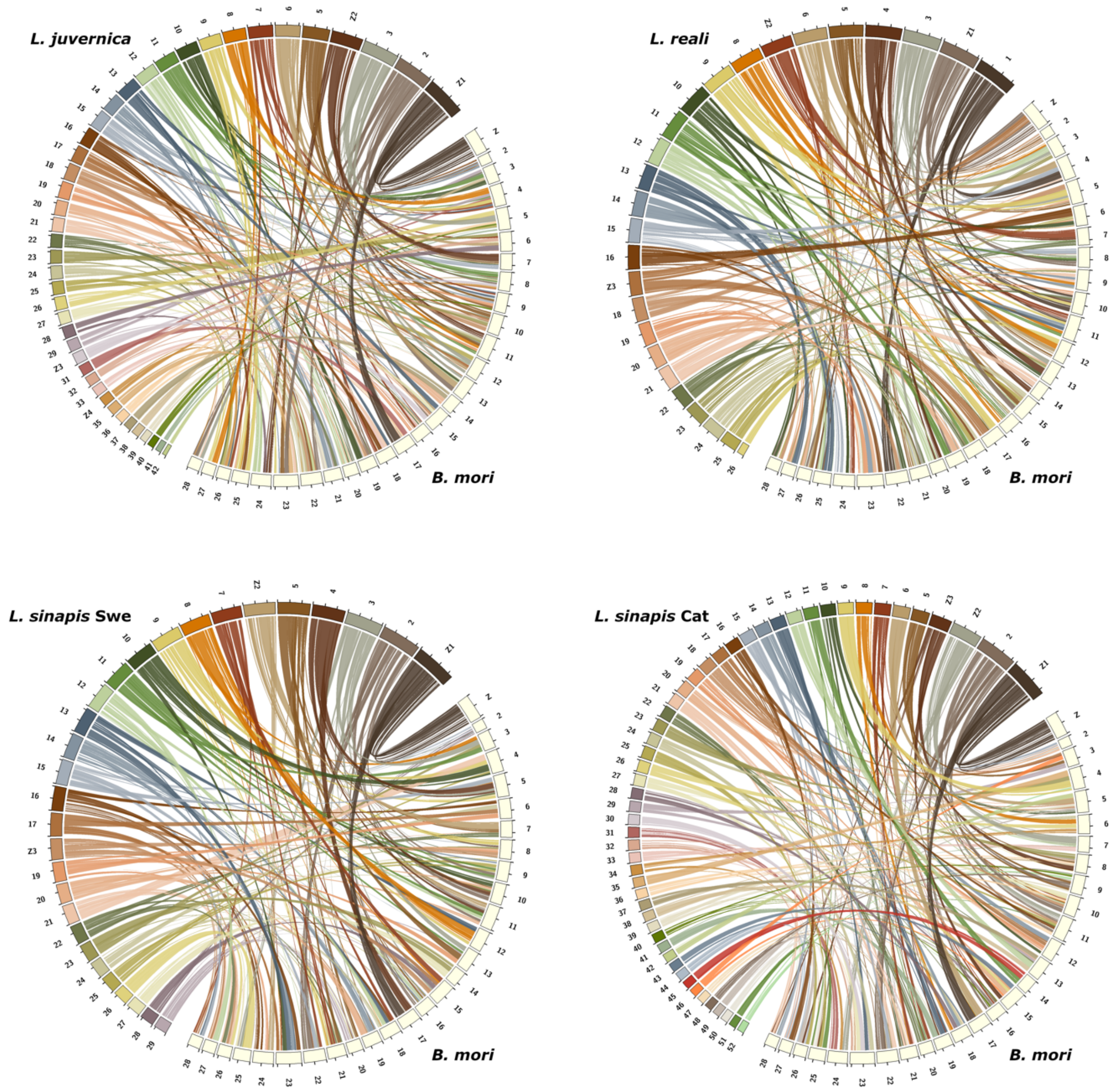

*Supplementary figure 5. Syntenic relationship between Leptidea species and Bombyx mori.*

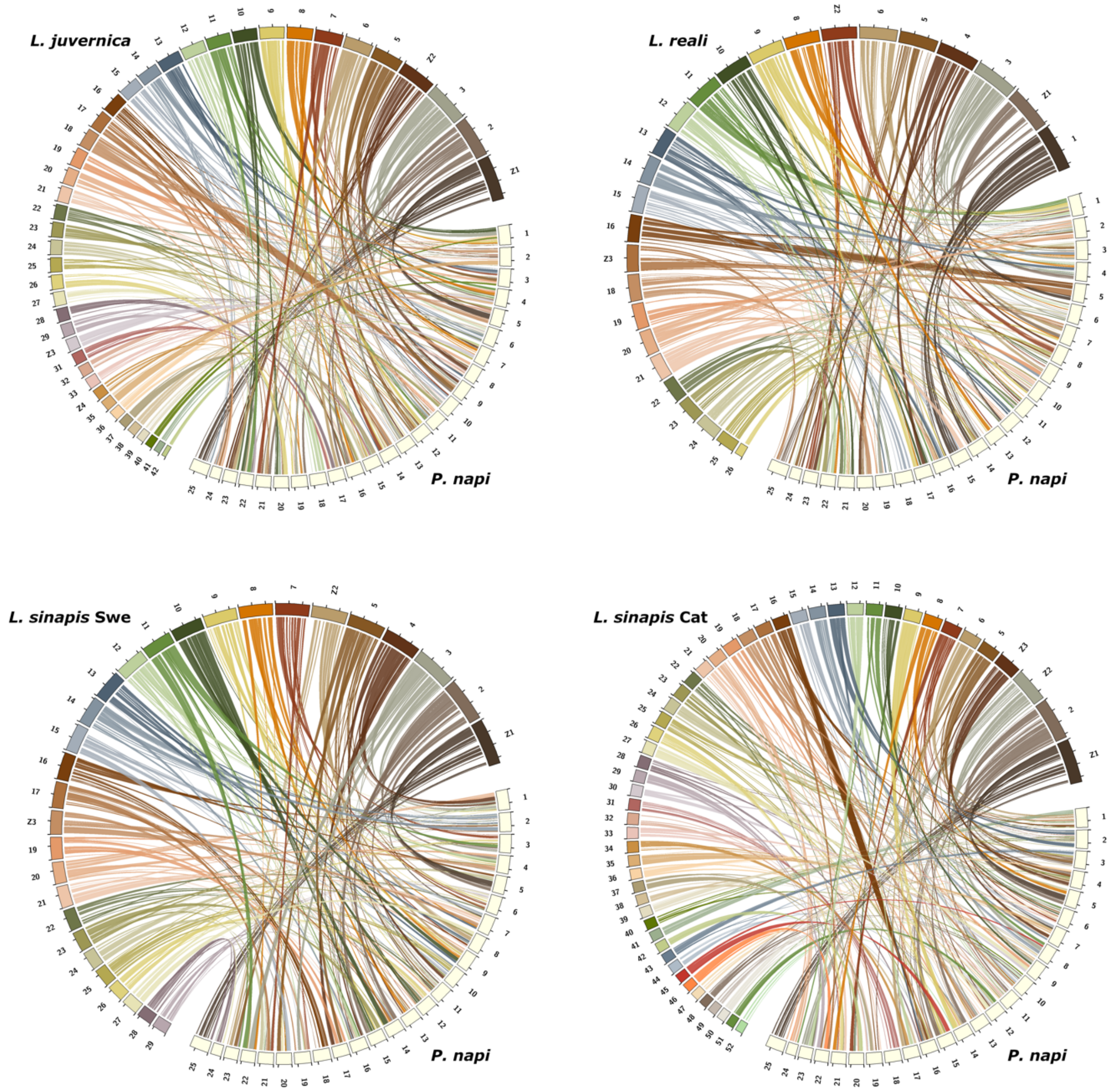

*Supplementary figure 6. Syntenic relationship between Leptidea species and Pieris napi.*

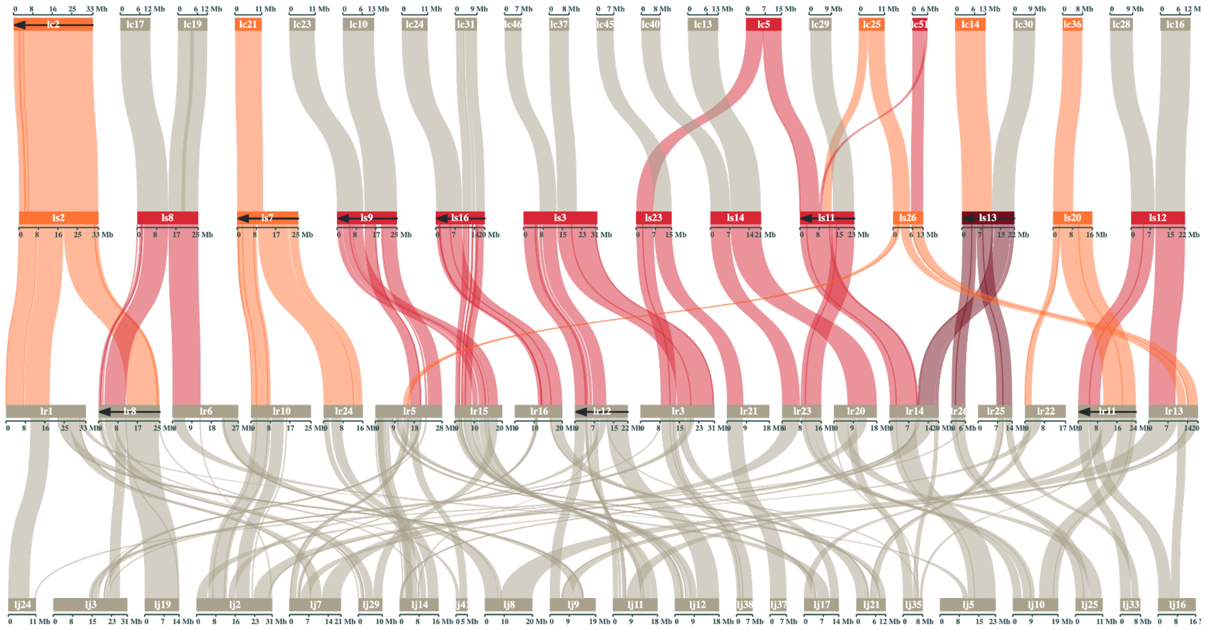

*Supplementary figure 7. Inferred fusion events in L. sinapis. Unique fusions are highlighted in red and ancestral fusions are highlighted in orange. Darker colour show chromosomes resulting from two events. Chromosomes with arrows have been rotated to simplify visualization. lc: L. sinapis Catalan karyotype, ls: L. sinapis Swedish karyotype, lr: L. reali and lj: L. juvernica.*

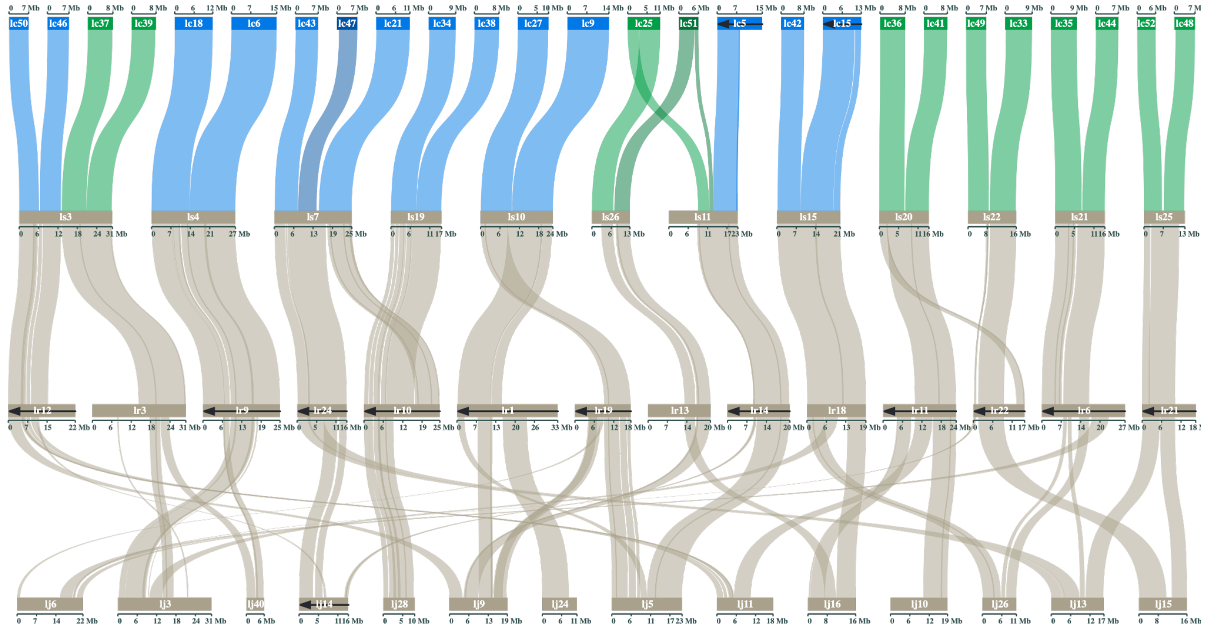

*Supplementary figure 8. Inferred fission events in L. sinapis. Unique fissions are highlighted in blue and fissions shared between L. sinapis Catalan karyotype and L. juvernica are highlighted in green. Darker colour show chromosomes resulting from two events. Chromosomes with arrows have been rotated to simplify visualization. lc: L. sinapis Catalan karyotype, ls: L. sinapis Swedish karyotype, lr: L. reali and lj: L. juvernica.*

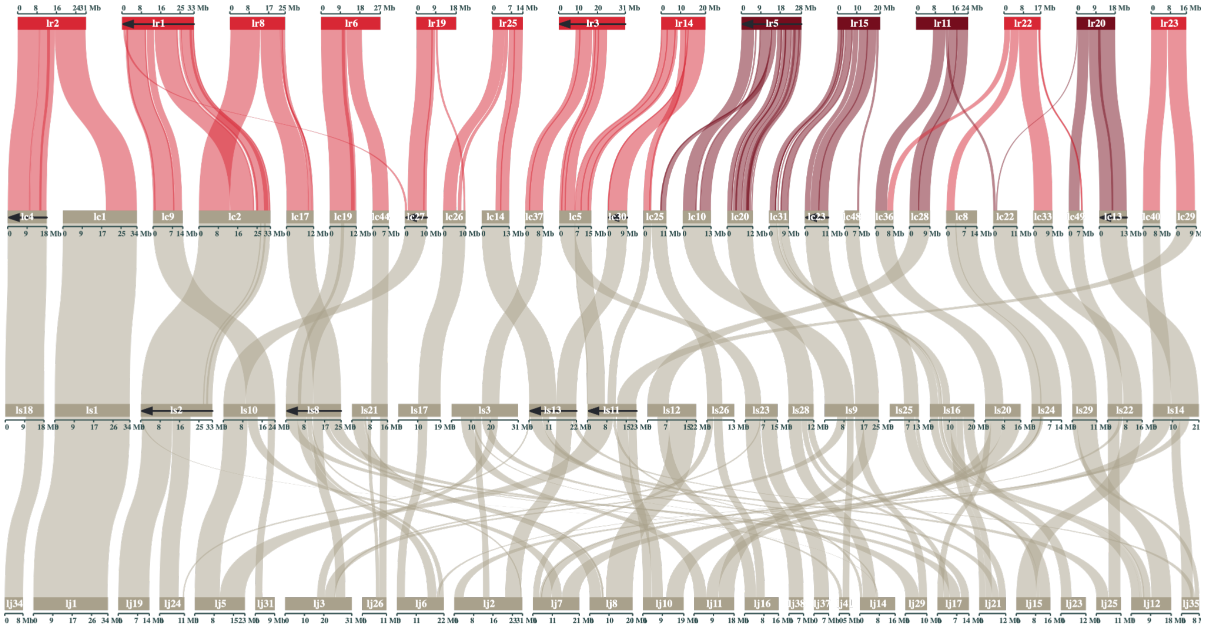

*Supplementary figure 9. Inferred fusion events in L. reali. Fusions are highlighted in red. Darker colour show chromosomes resulting from two events. Chromosomes with arrows have been rotated to simplify visualization. lr: L. reali , lc: L. sinapis Catalan karyotype, ls: L. sinapis Swedish karyotype and lj: L. juvernica.*

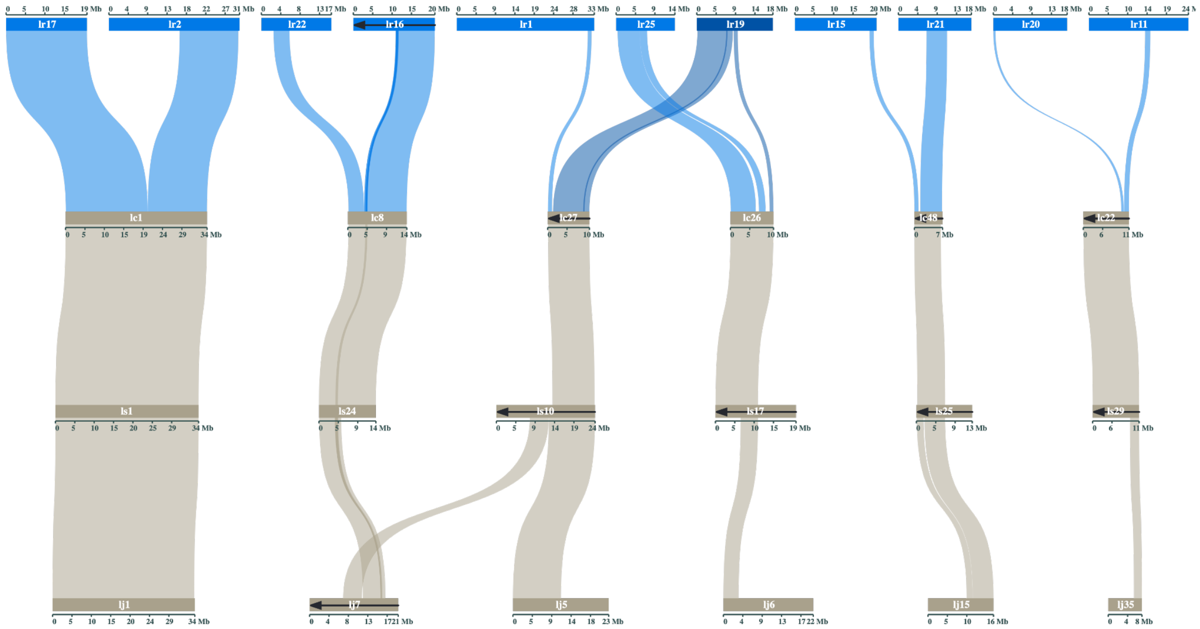

*Supplementary figure 10. Inferred fission events in L. reali. Fissions are highlighted in blue. Darker colour show chromosomes resulting from two events. Chromosomes with arrows have been rotated to simplify visualization. lr: L. reali, lc: L. sinapis Catalan karyotype, ls: L. sinapis Swedish karyotype and lj: L. juvernica.*

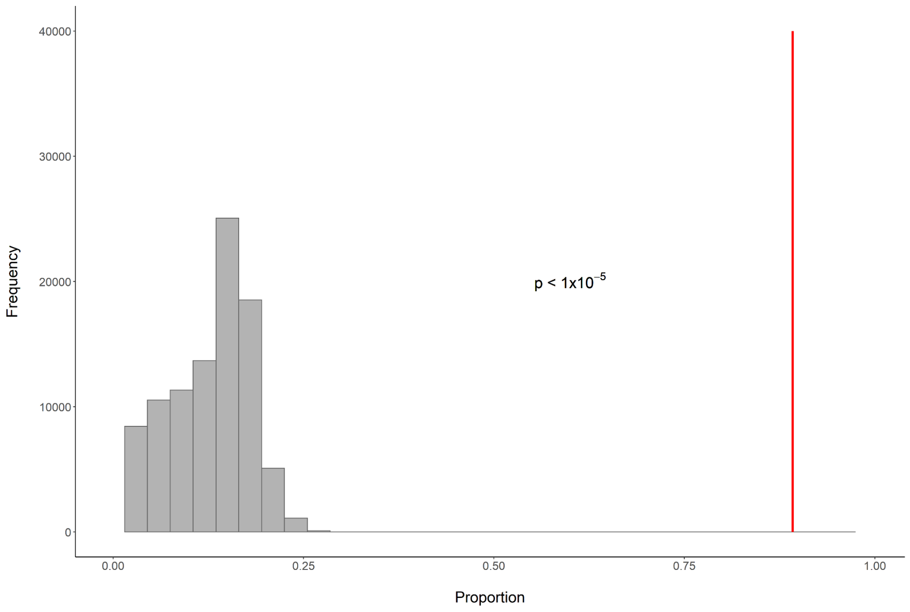

*Supplementary figure 11. Comparison of the observed proportion of identified fusions in Leptidea with overlapping breakpoints against M. cinxia to a distribution of random overlaps with breakpoints in Leptidea against M. cinxia. The distribution was generated by random sampling of non-overlapping windows across the combined Leptidea genomes and counting the proportion of windows that overlapped a breakpoint identified between Leptidea and M. cinxia. Each sample contained the equivalent number and sizes as all called breakpoints between Leptidea species and the sampling was repeated 100k times. The red line shows the observed proportion of called fusions that overlapped with breakpoints between Leptidea and M. cinxia. The p-value indicates the probability of observing the proportion given the distribution and was calculated with a one-tailed significance test.*

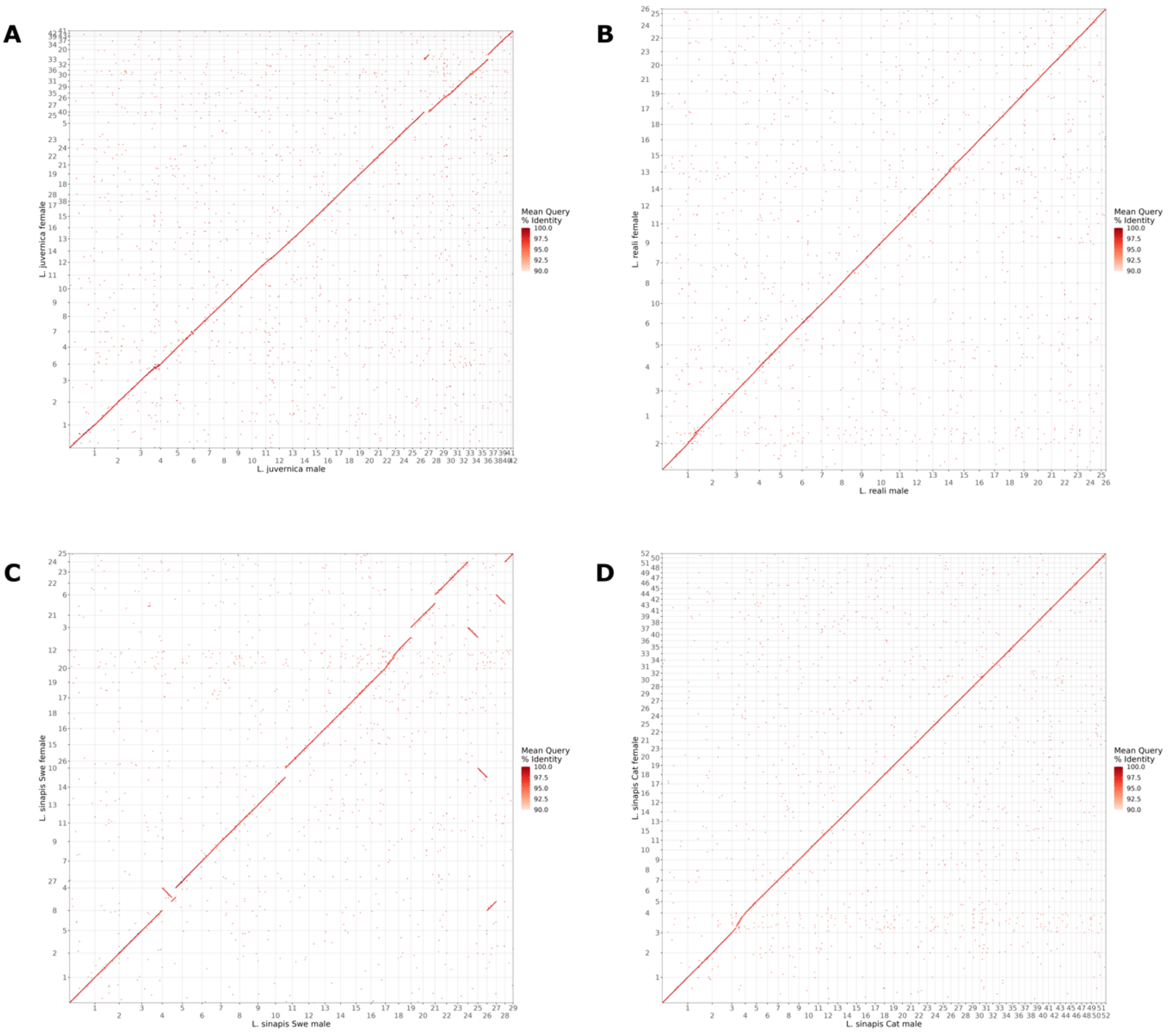

*Supplementary figure 12. Dotplots of alignments (>1kb) between Leptidea male and female assemblies. Scaffolds in the query are oriented so the majority of alignments are in the same direction as the reference. Mean query % identity shows mean identity for all alignments in each scaffold. A: L. juvernica, B: L. reali, C: L. sinapis Swedish population and D: L. sinapis Catalan population.*

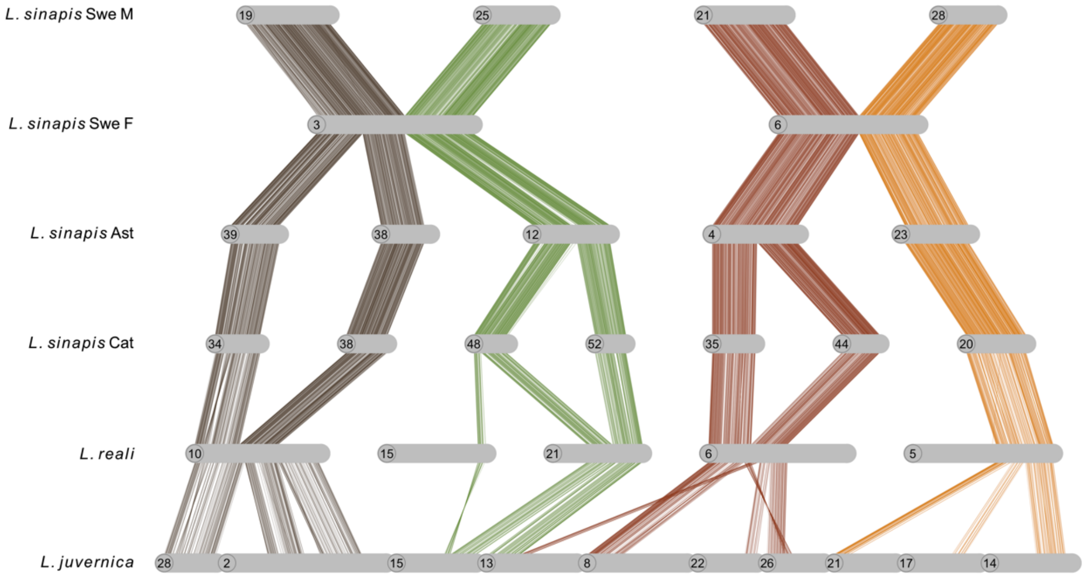

*Supplementary figure 13. Segregating chromosome fusions in the Swedish L. sinapis population identified in the genome assemblies. The two fusions were unique for the female individual. Chromosomes have been rotated to enhance visualization. Ast: Asturias population, Cat: Catalan population, Swe: Swedish population.*

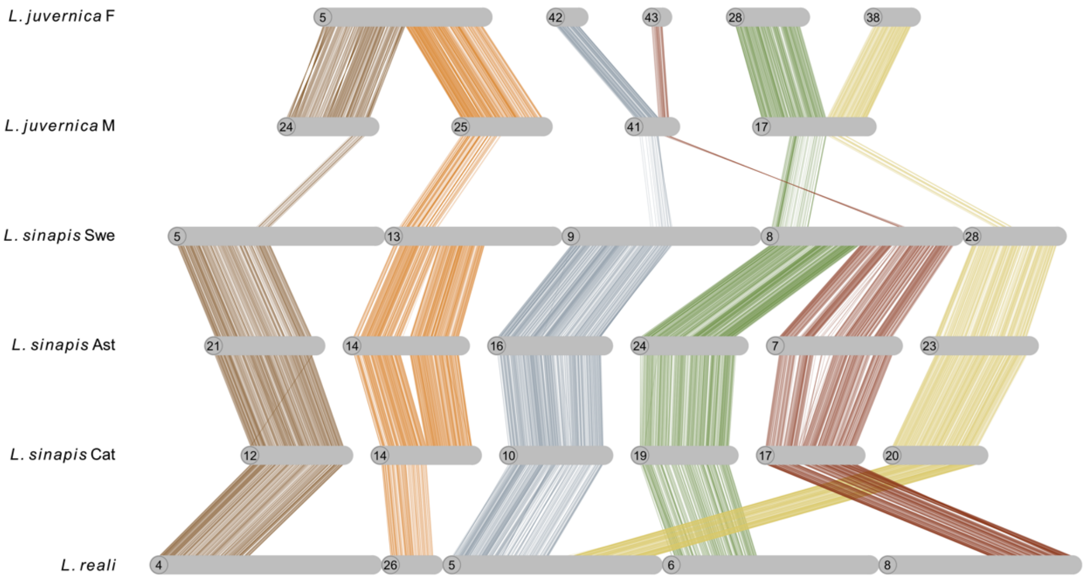

*Supplementary figure 14. Segregating chromosome fusions in L. juvernica. Lines show individual alignments (> 90% similarity) and colours represent homologous regions. Chromosomes have been rotated to enhance visualization. Ast: Asturias population, Cat: Catalan population, Swe: Swedish population.*

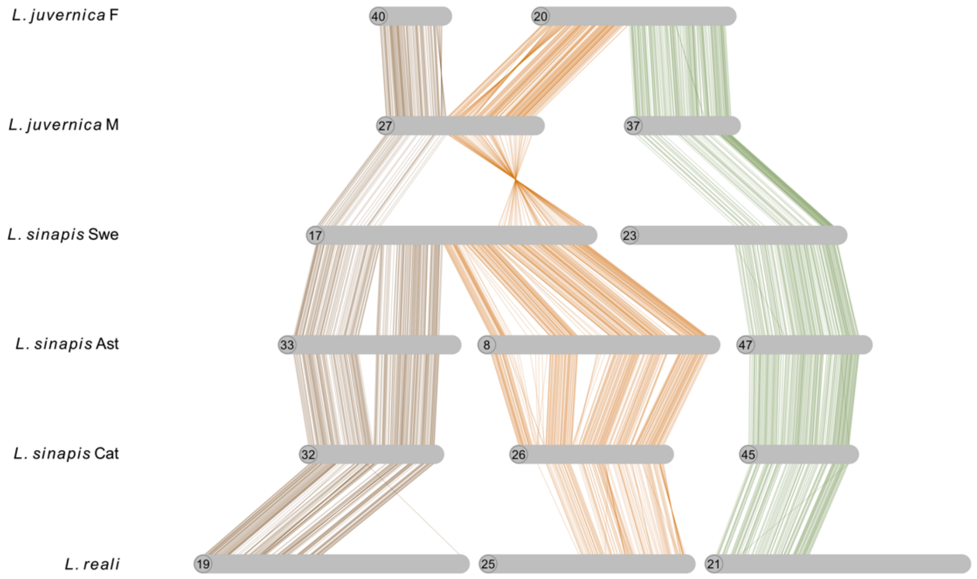

*Supplementary figure 15. Rearrangement polymorphism in L. juvernica. Lines show individual alignments (> 90% similarity) and colours represent homologous regions. Chromosomes have been rotated to enhance visualization. Ast: Asturias population, Cat: Catalan population, Swe: Swedish population.*

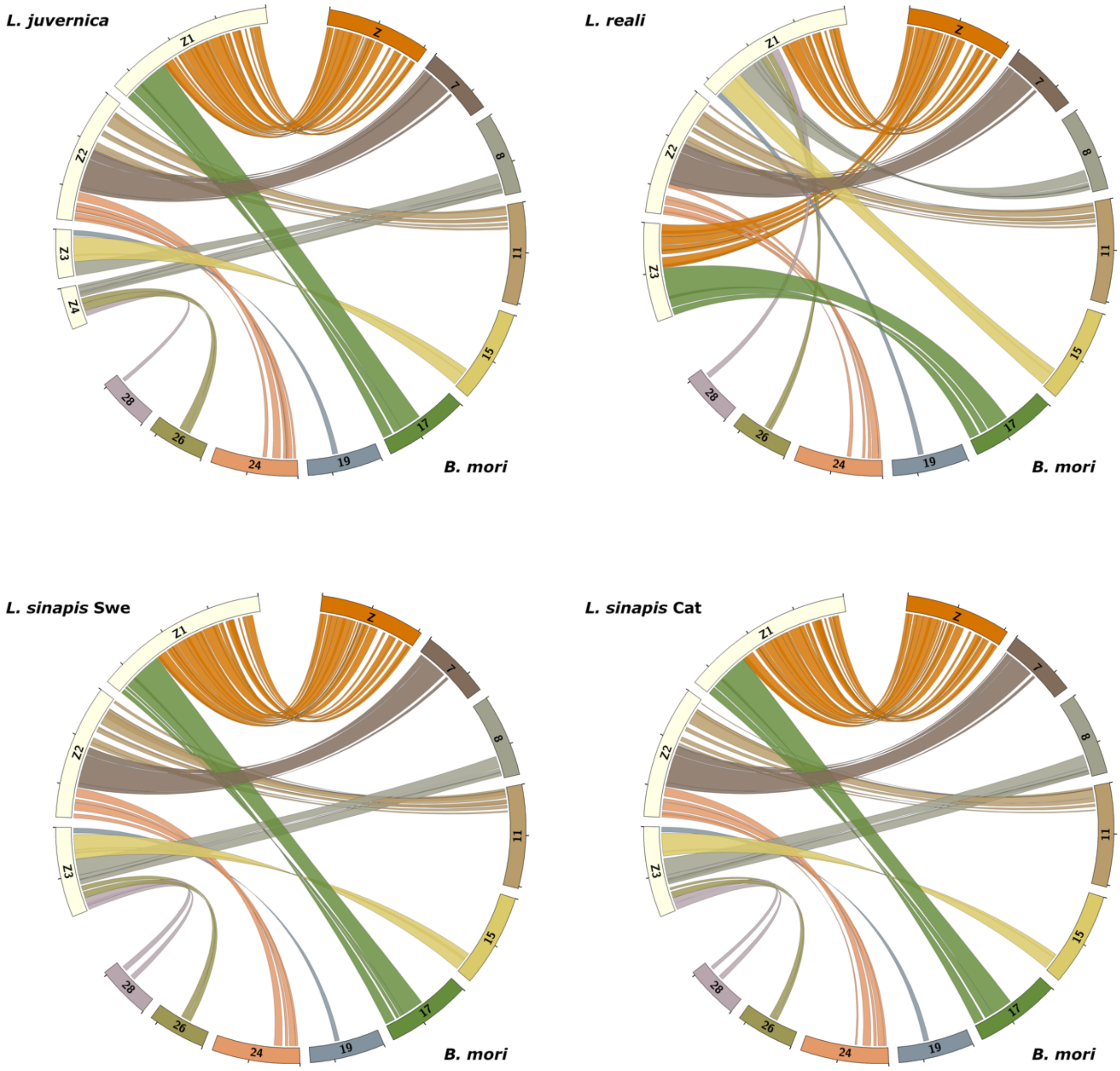

*Supplementary figure 16. Synteny between Leptidea sex chromosomes and homologous B. mori chromosomes. Individual Leptidea chromosomes have been rotated to simplify comparison between species.*

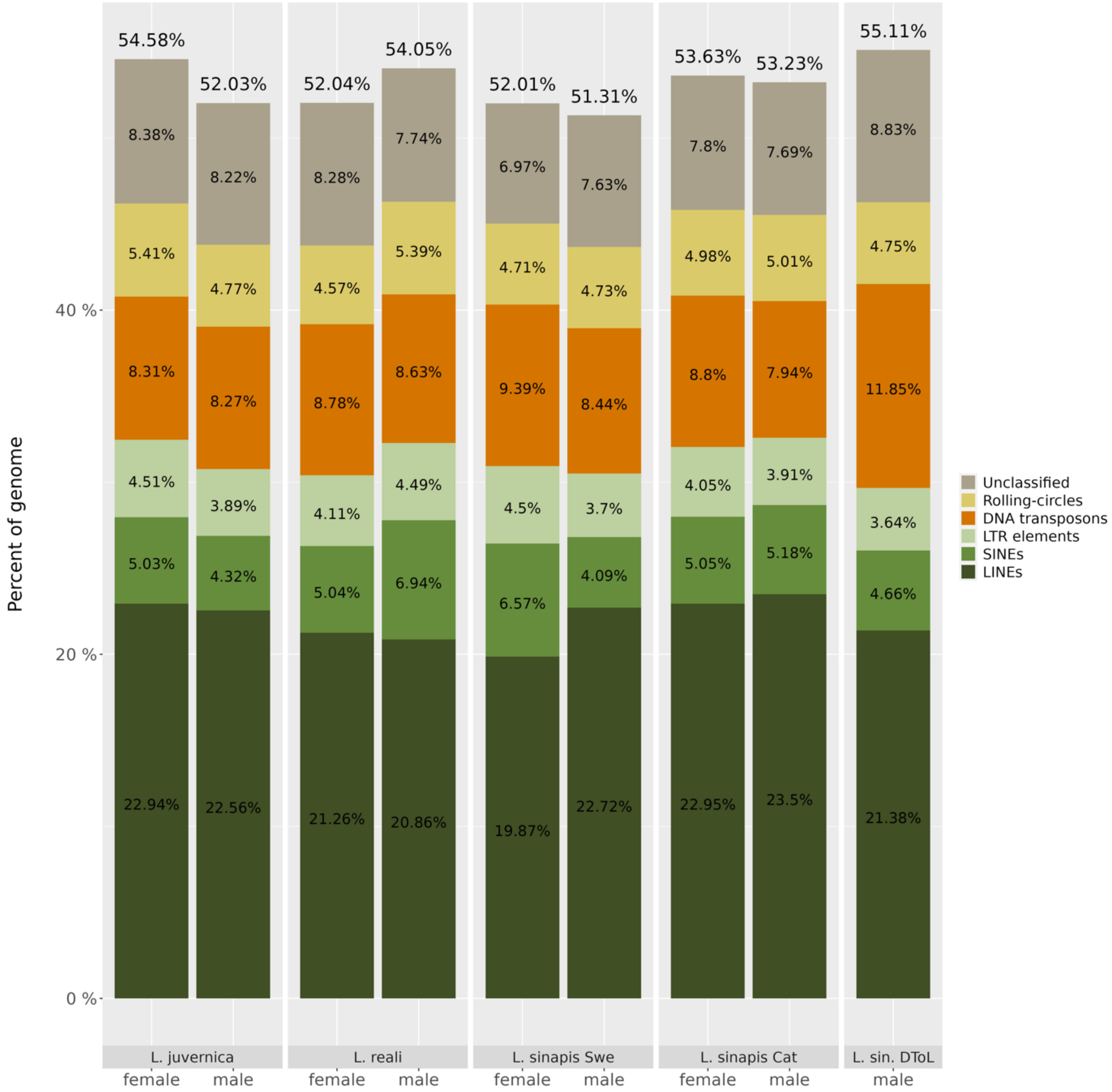

*Supplementary figure 17. Stacked barplot showing percent of different types of transposable element in the Leptidea genome assemblies. Percent values show assembly sequence content per repeat class and values on top of each bar shows total TE content. LINE = long interspersed elements, SINE = short interspersed elements, LTR = long terminal repeats.*

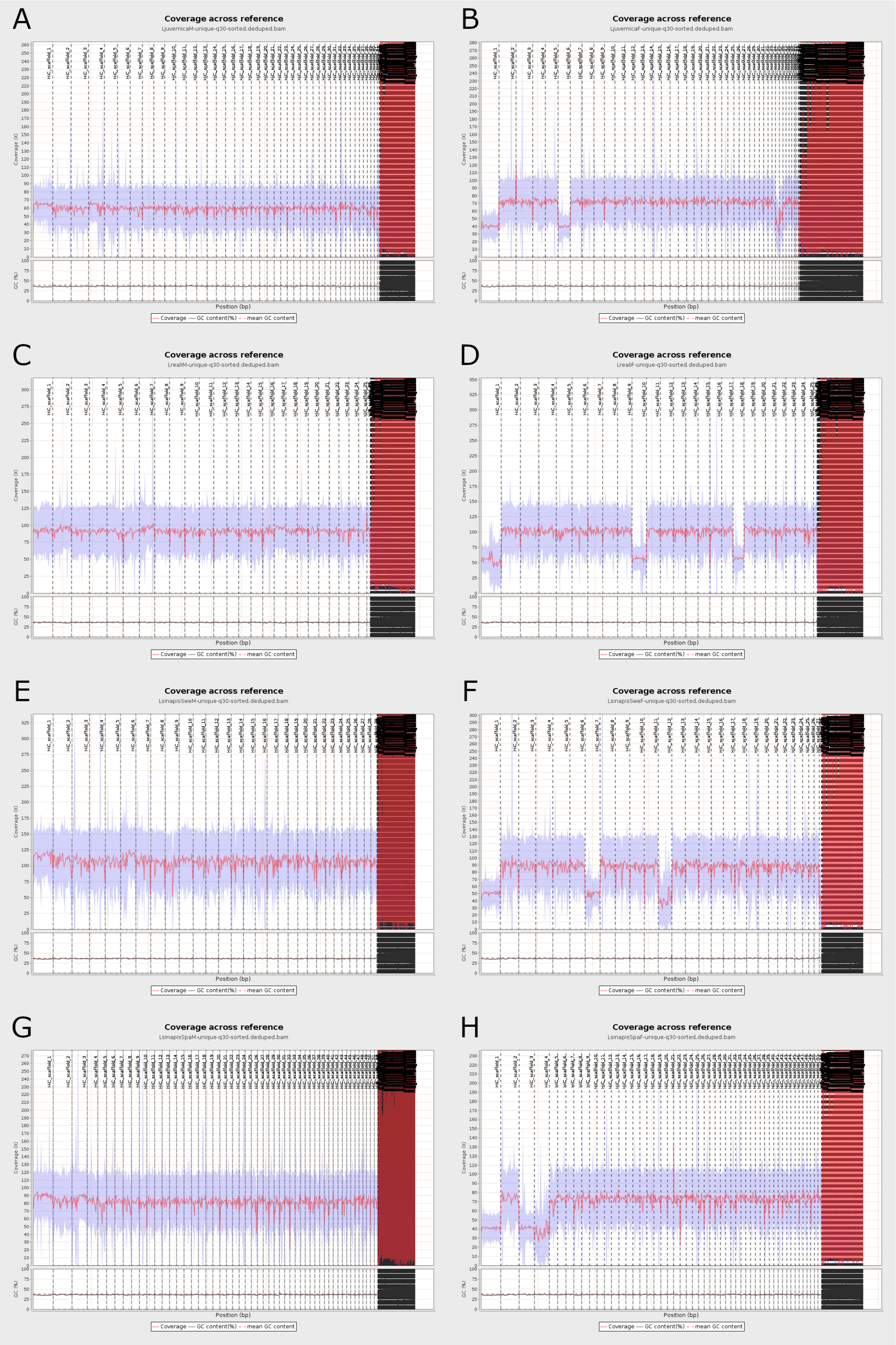

*Supplementary figure 18. Read mapping coverage and GC% across scaffolds. The red line shows mean coverage in 600 windows across the genome and the blue area shows standard deviation. Vertical dotted lines show scaffold boundaries. A: L. juvernica male, B: L. juvernica female, C: L. reali male, D: L. reali female, E: L. sinapis Swe male, F: L. sinapis Swe female, G: L. sinapis Cat male, H: L. sinapis female.*

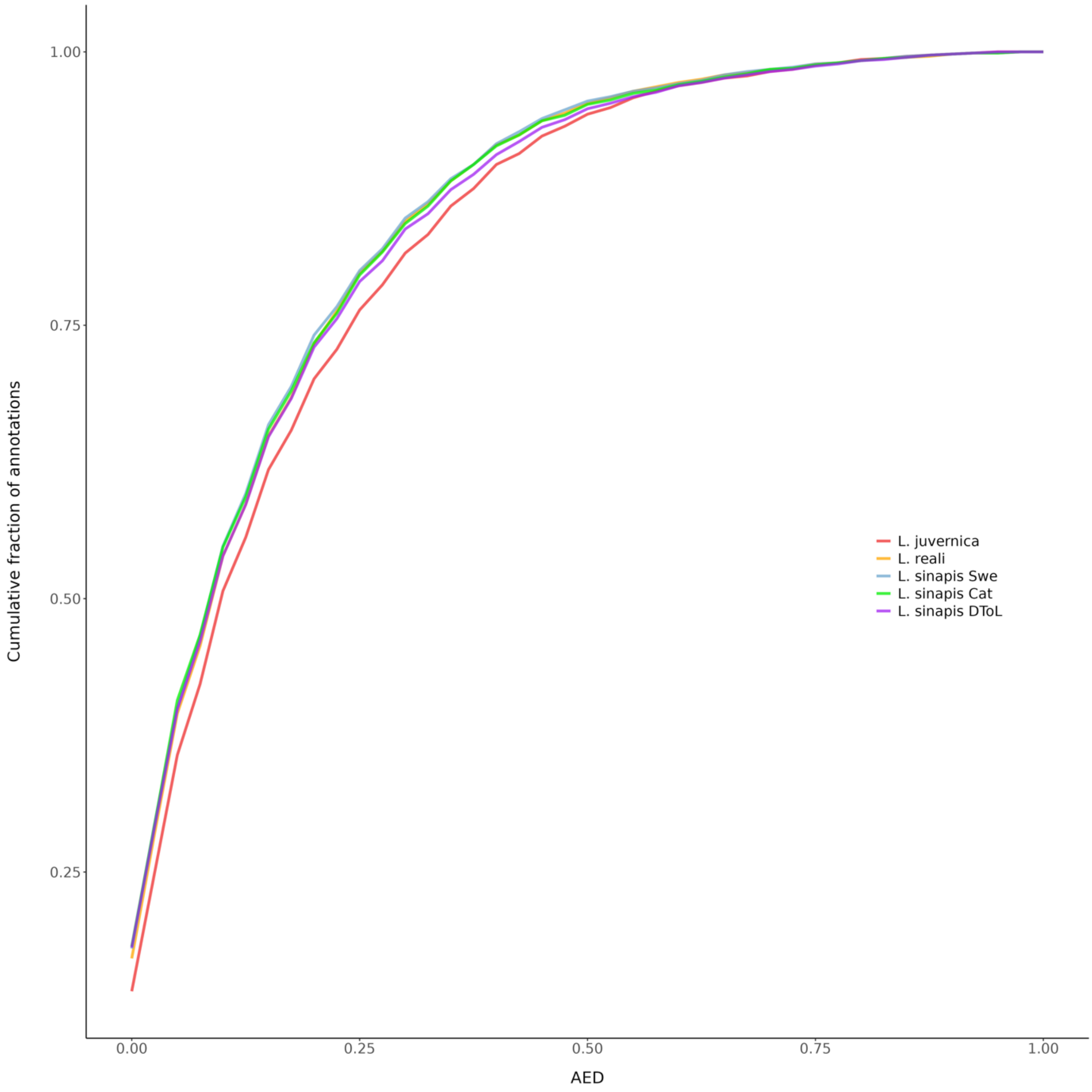

*Supplementary figure 19. Cumulative AED score for gene annotations. A score of 0 indicates full congruence and 1 indicates lack of congruence between the evidence and the final annotation.*

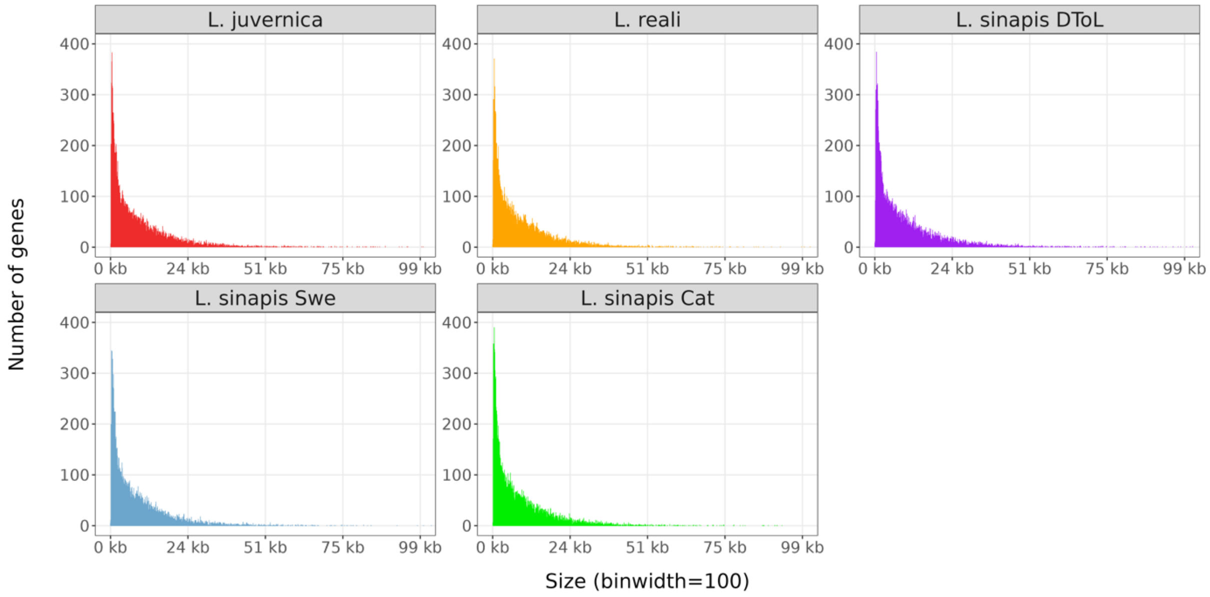

*Supplementary figure 20. Histogram showing size (bp) distribution of predicted genes. The x-axis has been capped at 100kb.*

#### Supplementary tables

Supplementary table 1. BUSCO scores in %, showing the estimated completeness of core insect genes. The database used was insecta_odb9 containing 1 658 genes. Values in parentheses show percent unit changes from uncorrected and unfiltered assemblies.

| Assembly | Complete (C) | Complete and single-copy (S) | Complete and duplicated (D) | Fragmented (F) | Missing (M) |
|---|---|---|---|---|---|
| *L. juvernica* male | 95.0% (+0.1%) | 93.7% | 1.3% (+0.1%) | 2.5% (-0.2) | 2.5% (+0.1%) |
| *L. juvernica* female | 93.7% (-0.1%) | 92.2% | 1.5% (-0.1%) | 3.5% (+0.1%) | 2.8% |
| *L. reali* male | 93.4% (+0.1%) | 92.2% (+0.2%) | 1.2% (-0.1%) | 3.8% (-0.1%) | 2.8% |
| *L. reali* female | 93.4% (+0.2%) | 92.2% (+0.8%) | 1.2% (-0.6%) | 3.8% | 2.8% (-0.2%) |
| *L. sinapis* Swe male | 93.8% (-0.1%) | 92.6% | 1.2% (-0.1%) | 3.1% | 3.1% (+0.1%) |
| *L. sinapis* Swe female | 93.9% | 92.6% | 1.3% | 3.1% | 3.0% |
| *L. sinapis* Cat male | 94.5% (+0.1%) | 93.1% | 1.4% (+0.1%) | 3.0% | 2.5% (-0.1%) |
| *L. sinapis* Cat female | 94.6% (+0.1%) | 93.2% (+0.3%) | 1.4% (-0.2%) | 2.5% | 2.9% (-0.1%) |

Supplementary table 2. Synteny statistics from comparison between *Leptidea* species and *B. mori*

| Statistic | *L. juvernica* | *L. reali* | *L. sinapis* Swe | *L. sinapis* Cat |
|---|---|---|---|---|
| Synteny blocks (N) | 410 | 398 | 372 | 390 |
| Median synteny blocks / chr | 14.00 | 14.50 | 13.00 | 12.50 |
| Median *B. mori* chr / chr | 4.00 | 5.00 | 5.00 | 3.00 |
| Median block size (mb) | 0.96 | 0.94 | 1.00 | 0.96 |
| Median genes / block | 25.00 | 24.00 | 24.00 | 24.00 |
| Median orthologs / block | 12.00 | 11.00 | 12.00 | 13.00 |

Supplementary table 3. Inferred fission and fusion events in *Leptidea*. Shaded chromosomes represent the reference for each inference. Columns with fusion events show the number of events for each respective reference chromosome and query chromosomes involved in fusions are in bold. Columns with fission events show which query chromosomes that result from an event. Values/chromosomes in parentheses show calls that change polarity after comparison with *M. cinxia* and detection of telomeric repeats. Direction of manual corrections are indicated by the last column.

| Chromosome number and homologous chromosomes in the other species | | | | Fusion | Fusion anc *L. sin* | Fission *L. sin* Cat | Fission *L. rea* | Fission *L.juv* / *L. sin* Cat | Manual Correction |
|---|---|---|---|---|---|---|---|---|---|
| *L. juvernica* | *L. reali* | *L. sinapis* Swe | *L. sinapis* Cat | | | | | | |
| 1 | 2, 17 | 1 | 1 | | | | 2-17 | | |
| 2 | 6, 10, 12, 14, 20 | 3, 7, 8, 13, 14, 19 | 13, 19, 21, 30, 38, 46 | | | | | | |
| 3 | 3, 4, 5, 8, 9, 18 | 3, 4, 5, 8, 9, 15 | 6, 7, 10, 15, 17, 18, 37, 42 | | | 6-18, 15-42 | | | |
| 4 | 7 | 6 | 3 | | | | | | |
| 5 | 1, 14, 19 | 10, 11 | 5, 9, 25, 27, 51 | | | (5-51), 9-27 | 1-19 | | |
| 6 | 4, 6, 19, 22, 25 | 5, 8, 17, 22 | 12, 19, 26, 49 | | | | 19-25 | | |
| 7 | 1, 5, 16, 22, 23 | 10, 14, 24, 26 | 8, 9, 25, 40 | | | | 16-22 | | |
| 8 | 6, 13 | 12, 21 | 16, 35 | | | | | | |
| 9 | 1, 12, 13, 19 | 3, 10, 17, 26 | 9, 32, 50, 51 | | | | | | |
| 10 | 4, 11, 23 | 5, 11, 20 | 7, 29, 41 | | | | | | |
| 11 | 11, 12, 15 | 3, 9, 20 | 23, 36, 46, 50 | | | 46-50 | | | |
| 12 | 12, 15 | 16, 29 | 22, 31 | | | | | | |
| 13 | 6, 21, 24 | 7, 21, 25 | 43, 44, 47, 52 | | | 43-47 | | | |
| 14 | 5, 12, 14, 24 | 3, 7, 11, 28 | 20, 21, 25, 46, 47 | | | 21-47 | | | |
| 15 | 15, 21, 22 | 22, 25 | 33, 48 | | | | 15-21 | | |
| 16 | 11, 13, 18 | 12, 15, 26 | 15, 25, 28 | | | | | | |
| 17 | 3, 5, 6, 22 | 3, 8, 20, 28 | 19, 20, 36, 37 | | | | | | |
| 18 | 1, 20 | 2, 14 | 2, 13 | | | | | | |
| 19 | 8 | 2 | 2 | | | | | | |
| 20 | 4 | 27 | 11 | | | | | | |
| 21 | 5, 16, 25 | 13, 16, 28 | 14, 20, 24 | | | | | | |
| 22 | 3, 6 | 5, 21 | 7, 44 | | | | | | |
| 23 | 9, 16 | 4, 16 | 18, 24 | | | | | | |
| 24 | 1, 4 | 2, 5 | 2, 12 | | | | | | |
| 25 | 20, 26 | 13, 22 | 14, 49 | | | | | | |
| 26 | 6, 18 | 15, 21 | 35, 42, 44 | | | | | | |
| 27 | 19, 25 | 17 | 26, 32 | | | | | | |
| 28 | 10 | 19 | 34, 38 | | | 34-38 | | | |
| 29 | 8, 15 | 8, 9 | 17, 23 | | | | | | |
| 30 | 2 | 18 | 4 | | | | | | |
| 31 | 1, 9 | 4, 10 | 6, 9 | | | | | | |
| 32 | 16 | 24 | 8 | | | | | | |
| 33 | 4, 11, 25 | 5, 12, 13 | 7, 14, 28 | | | | | | |
| 34 | 2 | 18 | 4 | | | | | | |

| | | | | | | | | | |
|---|---|---|---|---|---|---|---|---|---|
| 35 | 11, 14, 20 | 13, 14, 29 | 13, 22, 30 | | | | 11-20 | | |
| 36 | 3, 9 | 3, 4 | 6, 39 | | | | | | |
| 37 | 21 | 23 | 45 | | | | | | |
| 38 | 3 | 23 | 5 | | | | | | |
| 39 | 5 | 9 | 10 | | | | | | |
| 40 | 3, 9 | 3, 4 | 18, 39 | | | | | | |
| 41 | 5, 8 | 8, 9 | 10, 17 | | | | | | |
| 42 | 18 | 15 | 15 | | | | | | |
| 5, 7, 9, 18, **24**, **31** | 1 | **2**, **10** | **2**, **9**, 27 | 1 | | 9-27 | | | |
| **1**, 30, **34** | 2 | **1**, **18** | **1**, **4** | 1 | | | | | |
| 3, **17**, 22, 36, **38**, 40 | 3 | **3**, 5, **23** | **5**, 7, **37**, 39 | 1 | | | | 1 | |
| 3, 6, 10, 20, 24, 33 | 4 | 5, 27 | 7, 11, 12 | | | | | (1) | Fission -> Fusion |
| 3, **7**, 14, 17, **21**, 39, **41** | 5 | **9**, **26**, **28** | **10**, **20**, **25** | 2 | | | | | |
| **2**, 6, 8, 13, 17, 22, **26** | 6 | **8**, **21** | **19**, 35, **44** | 1 | | | | 1 | |
| 4 | 7 | 6 | 3 | | | | | | |
| **3**, **19**, 29, 41 | 8 | **2**, **8** | **2**, **17** | 1 | | | | | |
| 3, 23, 31, 36, 40 | 9 | 4 | 6, 18 | | | 6-18 | | | |
| 2, 28 | 10 | 7, 19 | 21, 34, 38 | | | 34-38 | | | |
| 10, **11**, **16**, 33, **35** | 11 | **12**, **20**, **29** | **22**, **28**, **36**, 41 | 2 | | | | 1 | |
| 2, 9, 11, 12, 14 | 12 | 3, 29 | 22, 46, 50 | | | 46-50 | | | |
| **8**, 9, **16** | 13 | **12**, **26** | **16**, **25**, 51 | (1) | | | | 1 | Fusion -> Fission |
| **2**, 5, **14**, 35 | 14 | **11**, **13** | 5, **25**, **30**, 51 | 1 | | 5-51 | | | |
| **11**, **12**, **15**, **29** | 15 | **9**, **16**, **25** | **23**, **31**, **48** | 2 | | | | | |
| **7**, 21, **23**, 32 | 16 | **16**, **24** | **8**, **24** | (1) | | | | | Fusion -> Fission |
| 1 | 17 | 1 | 1 | | | | | | |
| 3, 16, 26, 42 | 18 | 15 | 15, 42 | | | 15-42 | | | |
| **5**, **6**, 9, 27 | 19 | **10**, **17** | **26**, **27**, 32 | 1 | | | | (1) | Fission -> Fusion |
| 2, 18, **25**, **35** | 20 | **14**, **22**, **29** | **13**, **22**, **49** | 2 | | | | | |
| 13, **15**, **37** | 21 | **23**, **25** | **45**, **48**, 52 | (1) | | | | 1 | Fusion -> Fission |
| 6, **7**, **15**, 17 | 22 | 20, **22**, **24** | **8**, **33**, 36, 49 | 1 | | | | 1 | |
| **7**, **10** | 23 | **11**, **14** | **29**, **40** | 1 | | | | | |
| 13, 14 | 24 | 7 | 21, 43, 47 | | | 43-47, 21-47 | | | |
| **6**, **21**, 27, 33 | 25 | **13**, **17** | **14**, **26** | 1 | | | | | |
| 25 | 26 | 13 | 14 | | | | | | |
| 1 | 2, 17 | 1 | 1 | | | | 2 -17 | | |
| 18, **19**, **24** | **1**, **8** | 2 | 2 | | 1 | | | | |
| **2**, 3, 9, 11, 14, **17**, 36, 40 | **3**, **12** | 3 | **37**, 39, **46**, 50 | 1 | | 46-50 | | 1 | |
| 3, 23, 31, 36, 40 | 9 | 4 | 6,18 | | | 6-18 | | | |
| 3, 6, 10, 22, 24, 33 | 3, 4 | 5 | 7,12 | | | | | (1) | Fission -> Fusion |

| | | | | | | | | | |
|---|---|---|---|---|---|---|---|---|---|
| 4 | 7 | 6 | 3 | | | | | | |
| **2**, 13, **14** | **10**, **24** | 7 | 21, 43, 47 | | 1 | 21-47, 43-47 | | | |
| **2**, **3**, 6, 17, 29, 41 | **6**, **8** | 8 | **17**, **19** | 1 | | | | | |
| 3, 11, **29**, 39, **41** | **5**, **15** | 9 | **10**, **23** | 1 | | | | | |
| 5, 7, 9, 31 | 1, 19 | 10 | 9, 27 | | | 9-27 | 1-19 | | |
| 5, **10**, **14** | **14**, **23** | 11 | 5, **25**, **29**, 51 | 1 | | 5-51 | | | |
| **8**, **16**, 33 | **11**, **13** | 12 | **16**, **28** | 1 | | | | | |
| **2**, **21**, **25**, **33**, 35 | **14**, **25**, **26** | 13 | **14**, **30** | 1 | 1 | | | | |
| 2, **7**, 18, **35** | **20**, **23** | 14 | **13**, **40** | 1 | | | | | |
| 3, 16, 26, 42 | 18 | 15 | 15, 42 | | | 15-42 | | | |
| **12**, **21**, 23 | **15**, **16** | 16 | **24**, **31** | 1 | | | | | |
| 6, 9, 27 | 19, 25 | 17 | 26, 32 | | | | 19-25 | (1) | Fission -> Fusion |
| 30, 34 | 2 | 18 | 4 | | | | | | |
| 2, 28 | 10 | 19 | 34, 38 | | | 34-38 | | | |
| 10, **11**, **17** | **11**, **22** | 20 | 36, 41 | | 1 | | | 1 | |
| 8, 13, 22, 26 | 6 | 21 | 35, 44 | | | | | 1 | |
| 6, 15, 25 | 20, 22 | 22 | 33, 49 | | | | | 1 | |
| **37**, **38** | **3**, **21** | 23 | **5**, **45** | 1 | | | | | |
| 7, 32 | 16, 22 | 24 | 8 | | | | 16-22 | | |
| 13, 15 | 15, 21 | 25 | 48, 52 | | | | 15-21 | 1 | |
| **7**, 9, **16** | **5**, **13** | 26 | 25, 51 | | 1 | | | 1 | |
| 20 | 4 | 27 | 11 | | | | | | |
| 14, 17, 21 | 5 | 28 | 20 | | | | | | |
| 12, 35 | 11, 12, 20 | 29 | 22 | | | | 11-20 | | |
| 1 | 2, 17 | 1 | 1 | | | | 2-17 | | |
| 18, **19**, **24** | **1**, **8** | 2 | 2 | | 1 | | | | |
| 4 | 7 | 6 | 3 | | | | | | |
| 30, 34 | 2 | 18 | 4 | | | | | | |
| **5**, **38** | **3**, **14** | **11**, **23** | 5 | 1 | | | | | |
| 3, 31, 36 | 9 | 4 | 6 | | | | | | |
| 3, 10, 22, 33 | 3, 4 | 5 | 7 | | | | | | |
| 7, 32 | 16, 22 | 24 | 8 | | | | 16-22 | | |
| 5, 7, 9, 31 | 1 | 10 | 9 | | | | | | |
| 3, 39, 41 | 5 | 9 | 10 | | | | | | |
| 20 | 4 | 27 | 11 | | | | | | |
| 6, 24 | 4 | 5 | 12 | | | | | | |
| 2, 18, 35 | 20 | 14 | 13 | | | | | | |
| **21**, **25**, 33 | **25**, **26** | 13 | 14 | | 1 | | | | |
| 3, 16, 42 | 18 | 15 | 15 | | | | | | |
| 8 | 13 | 12 | 16 | | | | | | |
| 3, 29, 41 | 8 | 8 | 17 | | | | | | |
| 3, 23, 40 | 9 | 4 | 18 | | | | | | |
| 2, 6, 17 | 6 | 8 | 19 | | | | | | |
| 14, 17, 21 | 5 | 28 | 20 | | | | | | |

| | | | | | | | |
|---|---|---|---|---|---|---|---|
| **2**, **14** | **10**, **24** | 7 | 21 | | 1 | | |
| 12, 35 | 11, 12, 20 | 29 | 22 | | | | 11-20 |
| 11, 29 | 15 | 9 | 23 | | | | |
| 21, 23 | 16 | 16 | 24 | | | | |
| 5, **7**, 14, **16** | **5**, **13**, 14 | 11, 26 | 25 | | 1 | | |
| 6, 27 | 19, 25 | 17 | 26 | | | | 19-25 |
| 5 | 1, 19 | 10 | 27 | | | | 1-19 |
| 16, 33 | 11 | 12 | 28 | | | | |
| 10 | 23 | 11 | 29 | | | | |
| 2, 35 | 14 | 13 | 30 | | | | |
| 12 | 15 | 16 | 31 | | | | |
| 9, 27 | 19 | 17 | 32 | | | | |
| 15 | 22 | 22 | 33 | | | | |
| 28 | 10 | 19 | 34 | | | | |
| 8, 26 | 6 | 21 | 35 | | | | |
| **11**, **17** | **11**, **22** | 20 | 36 | | 1 | | |
| 3, 17 | 3 | 3 | 37 | | | | |
| 2, 28 | 10 | 19 | 38 | | | | |
| 36, 40 | 3 | 3 | 39 | | | | |
| 7 | 23 | 14 | 40 | | | | |
| 10 | 11 | 20 | 41 | | | | |
| 3, 26 | 18 | 15 | 42 | | | | |
| 13 | 24 | 7 | 43 | | | | |
| 13, 22, 26 | 6 | 21 | 44 | | | | |
| 37 | 21 | 23 | 45 | | | | |
| 2, 11, 14 | 12 | 3 | 46 | | | | |
| 13, 14 | 24 | 7 | 47 | | | | |
| 15 | 15, 21 | 25 | 48 | | | | 15-21 |
| 6, 25 | 20, 22 | 22 | 49 | | | | |
| 9, 11 | 12 | 3 | 50 | | | | |
| **5**, **9** | **13**, **14** | **11**, **26** | 51 | 1 | | | |
| 13 | 21 | 25 | 52 | | | | |

Supplementary table 4. Summary statistics of genetic element densities in fusion and fission breakpoints. Densities are shown as the mean percent of a breakpoint type covered by each sequence element type. P-values were calculated using two-tailed significance tests and indicate the probability of observing the mean given resampling distributions taken from the rest of the genomes. CDS = coding sequence, LINE = long interspersed elements, SINE = short interspersed elements, LTR = long terminal repeats, DNA = dna transposons, RC = rolling-circle TEs.

| Breakpoint | Inference type | Element | Mean (%) | Standard deviation (%) | FDR adjusted p-value | Raw p-value |
|---|---|---|---|---|---|---|
| fusion | reference | CDS | 2.54 | 1.45 | 0.932 | 0.918 |
| ” | “ | LINE | 29.33 | 7.23 | **< 2*10^-5** | < 2*10^-5 |
| “ | “ | SINE | 3.46 | 1.64 | **< 2*10^-5** | < 2*10^-5 |
| “ | “ | LTR | 6.30 | 3.02 | **< 2*10^-5** | < 2*10^-5 |
| “ | “ | DNA | 8.39 | 2.71 | 0.932 | 0.893 |
| “ | “ | RC | 4.15 | 2.26 | **0.003** | 0.001 |
| “ | query | CDS | 2.71 | 2.81 | 0.252 | 0.126 |
| “ | “ | LINE | 28.80 | 10.05 | **< 2*10^-5** | < 2*10^-5 |
| “ | “ | SINE | 3.63 | 3.07 | **< 2*10^-5** | < 2*10^-5 |
| “ | “ | LTR | 5.16 | 4.15 | 0.096 | 0.044 |
| “ | “ | DNA | 7.95 | 3.93 | 0.096 | 0.043 |
| “ | “ | RC | 4.50 | 2.92 | **< 2*10^-5** | < 2*10^-5 |
| fission | reference | CDS | 2.92 | 2.40 | 0.644 | 0.456 |
| ” | “ | LINE | 24.54 | 5.93 | 0.919 | 0.805 |
| “ | “ | SINE | 4.83 | 2.25 | 0.644 | 0.432 |
| “ | “ | LTR | 4.29 | 2.63 | 0.448 | 0.280 |
| “ | “ | DNA | 9.05 | 3.43 | 0.370 | 0.205 |
| “ | “ | RC | 5.21 | 2.33 | 0.853 | 0.711 |
| “ | query | CDS | 4.40 | 5.34 | 0.077 | 0.029 |
| “ | “ | LINE | 24.24 | 7.15 | 0.853 | 0.699 |
| “ | “ | SINE | 5.18 | 2.44 | 0.932 | 0.932 |
| “ | “ | LTR | 4.65 | 3.15 | 0.216 | 0.370 |
| “ | “ | DNA | 9.05 | 4.01 | 0.758 | 0.569 |
| “ | “ | RC | 4.32 | 1.92 | **0.040** | 0.013 |

Supplementary table 5. LINE element families enriched in telomeric regions in the Asturian *L. sinapis* assembly, identified with Fisher's exact test. The table is limited to the top five candidate families which are defined by occurring in more than half of the chromosomes and at least the same number of copies as the number of telomeres (n=96).

| LINE family | Odds ratio | FDR-adjusted p-value | Raw p-value |
|---|---|---|---|
| rnd-4_family-1088 LINE/R1 | 120.57 | $< 1 \times 10^{-189}$ | $< 1 \times 10^{-191}$ |
| rnd-4_family-5536 LINE/R1 | 58.41 | $< 1 \times 10^{-189}$ | $< 1 \times 10^{-191}$ |
| rnd-4_family-980 LINE/R1 | 33.93 | $1 \times 10^{-189}$ | $1 \times 10^{-191}$ |
| rnd-5_family-614 LINE/R2 | 39.99 | $1 \times 10^{-136}$ | $1 \times 10^{-138}$ |
| rnd-5_family-1424 LINE/I-Jockey | 6.30 | $2 \times 10^{-57}$ | $2 \times 10^{-59}$ |

Supplementary table 6. Female identity and number of offspring per female (# offspring) used in the pedigree for constructing linkage maps in the Swedish and Catalonian *L. sinapis* populations.

| Swedish | | Catalonian | |
|---|---|---|---|
| Female | # offspring | Female | # offspring |
| T2 | 41 | 3c9 | 38 |
| T3 | 31 | 4C | 35 |
| T4 | 30 | 7C | 34 |
| T5 | 58 | 7CB | 20 |
| T6 | 14 | 8C | 27 |
| S25 | 10 | 9C | 24 |
| Total | 184 | Total | 178 |

Supplementary table 7. Sex chromosome identification and statistics. Note that Z chromosomes with similar chromosome names are not necessarily homologous between species.

| Assembly | Chromosome name | Scaffold name | Size (mb) | Chr:genome mean read depth ratio |
|---|---|---|---|---|
| *L. juvernica* male | Z1 | HiC_scaffold_1 | 34.1 | 1.0533 |
| | Z2 | HiC_scaffold_4 | 26.8 | 1.0216 |
| | Z3 | HiC_scaffold_30 | 9.7 | 0.9931 |
| | Z4 | HiC_scaffold_34 | 8.3 | 1.0031 |
| *L. juvernica* female | Z1 | HiC_scaffold_1 | 31.2 | 0.5847 |
| | Z2 | HiC_scaffold_6 | 22.0 | 0.5793 |
| | Z3 | HiC_scaffold_35 | 6.4 | 0.5816 |
| | Z4 | HiC_scaffold_36 | 6.3 | 0.7444 |
| *L. reali* male | Z1 | HiC_scaffold_2 | 31.4 | 1.0356 |
| | Z2 | HiC_scaffold_7 | 25.4 | 1.0480 |
| | Z3 | HiC_scaffold_17 | 19.4 | 1.0585 |
| *L. reali* female | Z1/W | HiC_scaffold_1 | 35.4 | 0.5650 |
| | Z2 | HiC_scaffold_10 | 25.4 | 0.5988 |
| | Z3 | HiC_scaffold_18 | 19.6 | 0.6036 |
| *L. sinapis* Swe male | Z1 | HiC_scaffold_1 | 34.5 | 1.1004 |
| | Z2 | HiC_scaffold_6 | 26.6 | 1.0702 |
| | Z3 | HiC_scaffold_18 | 17.8 | 1.0444 |
| *L. sinapis* Swe female | Z1 | HiC_scaffold_1 | 34.5 | 0.6211 |
| | Z2 | HiC_scaffold_7 | 27.2 | 0.6058 |
| | Z3/W | HiC_scaffold_12 | 24.4 | 0.5066 |
| | W | HiC_scaffold_28 | 4.4 | 0.4936 |
| *L. sinapis* Cat male | Z1 | HiC_scaffold_1 | 34.0 | 1.1052 |
| | Z2 | HiC_scaffold_3 | 26.2 | 1.0653 |
| | Z3 | HiC_scaffold_4 | 17.9 | 1.0207 |
| *L. sinapis* Cat female | Z1 | HiC_scaffold_1 | 34.2 | 0.6146 |
| | Z2 | HiC_scaffold_3 | 26.9 | 0.6140 |
| | Z3/W | HiC_scaffold_4 | 26.3 | 0.5441 |

Supplementary table 8. Gene annotation statistics.

| Assembly | Predicted genes (N) | Median gene size (bp) | Median CDS size (bp) | Median number of exons (N) | Median exon size (bp) | Genes on chromosome scaffolds (%) | Genes with functional predictions (N) | Predicted protein domains (N) |
|---|---|---|---|---|---|---|---|---|
| *L. juvernica* | 16 149 | 5 444 | 789 | 4 | 148 | 92.41% | 9 441 | 22 676 |
| *L. reali* | 15 689 | 5702 | 768 | 4 | 144 | 87.51% | 9 256 | 21 452 |
| *L. sinapis* Swe | 15 915 | 5 453 | 762 | 4 | 144 | 90.34% | 9 348 | 21 790 |
| *L. sinapis* Cat | 16 478 | 5 375 | 762 | 4 | 145 | 91.69% | 9 723 | 22 724 |
| *L. sinapis* DToL | 17 229 | 6018 | 819 | 4 | 145 | 100% | 9 935 | 23 684 |
